# The Parkinson’s Disease Mendelian Randomization Research Portal

**DOI:** 10.1101/604033

**Authors:** Alastair J Noyce, Sara Bandres-Ciga, Jonggeol Kim, Karl Heilbron, Demis Kia, Gibran Hemani, Angli Xue, Debbie A Lawlor, George Davey Smith, Raquel Duran, Ziv Gan-Or, Cornelis Blauwendraat, J Raphael Gibbs, 23andMe Research Team, International Parkinson’s Disease Genomics Consortium (IPDGC), David A Hinds, Jian Yang, Peter Visscher, Jack Cuzick, Huw Morris, John Hardy, Nicholas W Wood, Mike A Nalls, Andrew B Singleton

**Affiliations:** Preventive Neurology Unit, Wolfson Institute of Preventive Medicine, Queen Mary University of London, London EC1M 6BQ, UK; Department of Clinical and Movement Neurosciences, University College London, Institute of Neurology, London WC1N 1PJ, UK; Molecular Genetics Section, Laboratory of Neurogenetics, National Institute on Aging, National Institutes of Health, Bethesda, MD 20892, USA; Instituto de Investigación Biosanitaria de Granada (ibs.GRANADA), Granada, Spain; 23andMe, Inc., 899 W Evelyn Avenue, Mountain View, CA 94041 USA; MRC Integrative Epidemiology Unit, University of Bristol, Bristol BS8 2BN, UK; Institute for Molecular Bioscience, The University of Queensland, Brisbane 4072, Australia; Queensland Brain Institute, The University of Queensland, Brisbane, Queensland 4072, Australia; Population Health Science, Bristol Medical School, University of Bristol, Bristol, UK; Centro de Investigacion Biomedica and Departamento de Fisiologia, Facultad de Medicina, Universidad de Granada, Granada, Spain; Department of Neurology & Neurosurgery, McGill University, Montreal, Quebec, Canada; Montreal Neurological Institute, McGill University, Montreal, Quebec, Canada; Department of Human Genetics, McGill University, Montreal, Quebec, Canada; Institute for Advanced Research, Wenzhou Medical University, Wenzhou, Zhejiang 325027, China; Data Tecnica International, Glen Echo, MD 20812, USA

**Author notes:** Equal contribution (Alastair J Noyce, Sara Bandres-Ciga, Jonggeol Kim). **Corresponding author:** Dr Alastair Noyce, Preventive Neurology Unit, Wolfson Institute of Preventive Medicine, Barts and the London School of Medicine and Dentistry, Queen Mary University of London, London, EC1M 6BQ UK.

**Keywords:** Mendelian randomization, Parkinson’s disease, risk factor, public resource

## Abstract

**Background:** Mendelian randomization (MR) is a method for exploring observational associations to find evidence of causality.

**Objective:** To apply MR between multiple risk factors/phenotypic traits (exposures) and Parkinson’s disease (PD) in a large, unbiased manner, and to create a public resource for research.

**Methods:** We used two-sample MR in which the summary statistics relating to SNPs from genome wide association studies (GWASes) of 5,839 exposures curated on *MR Base* were used to assess causal relationships with PD. We selected the highest quality exposure GWASes for this report (n=401). For the disease outcome, summary statistics from the largest published PD GWAS were used. For each exposure, the causal effect on PD was assessed using the inverse variance weighted (IVW) method, followed by a range of sensitivity analyses. We used a false discovery rate (FDR) corrected p-value of <0.05 from the IVW analysis to prioritize traits of interest.

**Results:** We observed evidence for causal associations between twelve exposures and risk of PD. Of these, nine were causal effects related to increasing adiposity and decreasing risk of PD. The remaining top exposures that affected PD risk were tea drinking, time spent watching television and forced vital capacity, but the latter two appeared to be biased by violations of underlying MR assumptions.

**Discussion:** We present a new platform which offers MR analyses for a total of 5,839 GWASes versus the largest PD GWASes available (https://pdgenetics.shinyapps.io/pdgenetics/). Alongside, we report further evidence to support a causal role for adiposity on lowering the risk of PD.

## INTRODUCTION

The risk factors and determinants of Parkinson’s disease (PD) remain incompletely understood. While monogenic forms are responsible for approximately 5% of PD, the vast majority of disease is considered to be sporadic and due to a number of genetic and non-genetic risk factors ^1^. Approximately one third of the genomic liability in PD risk has been explained by common genetic variants uncovered by current genome-wide association studies (GWAS) ^2,3^. Total heritability estimates are estimated to be roughly 22-27%, meaning that a substantial proportion of genetic risk is still to be discovered ^4,5^. The remainder of PD risk likely comes from environmental factors ^4,5^, aging ^6,7^, and stochastic events occuring in an unpredictable and non-modifiable manner ^8^.

Mendelian randomization (MR) is an epidemiological method that can be used to provide support for causality between a modifiable exposure/risk factor/phenotypic trait (henceforth collectively termed *exposure*) and a disease outcome ^9^. Put simply, genetic variants (usually single nucleotide polymorphisms or SNPs) that explain variation in an exposure can be used as proxies to determine how a change in that exposure might influence a disease outcome. A ratio of the genetically-estimated change in an exposure and the genetically-estimated change in the outcome using the same individual SNP is calculated and then pooled across all SNPs that are independently associated with the exposure trait of interest. The pooled ratio (usually a Wald ratio) is an estimate of change in the outcome for a given change in the exposure, as long as certain instrumental variable (IV) assumptions are upheld (see **Supplemental Material**).

A common approach to MR involves the use of summary statistics from published GWASes of exposures and the summary statistics of a GWAS of an outcome to determine causal estimates; an approach known as two-sample MR. Recently, the summary statistics from GWAS for a large range of exposures have been curated in *MR Base* (http://www.mrbase.org), which enables targeted (hypothesis-driven) exploration of causal associations or hypothesis-generating approaches to MR ^10^. Many MR studies are underpowered, either as a consequence of relative low samples sizes or the small amount of variation in an exposure trait which is explained by common genetic variation. We have undertaken a high-throughput approach to two-sample MR for a large range of exposures and PD. The principal goal was to provide a resource for the research community to add causal insights to associations arising from traditional epidemiological approaches and to support pursuit of new interventions to reduce risk of PD.

## METHODS

### Exposure data

*MR Base* is an online resource which, at the time of the analysis, contained summary results from 7,956 GWASes across multiple exposures, encompassing a wide range of physiological characteristics and disease phenotypes. *MR Base* was accessed on 14th January 2019. Each exposure was tested separately to determine if it altered risk of PD. All analyses were performed using the R package *TwoSampleMR* (version 3.2.2; https://github.com/MRCIEU/TwoSampleMR). The instrumental variables used for each binary exposure consisted of the per-allele log-odds ratio (or the beta estimate for continuous exposure traits) and standard errors for all independent SNPs reaching genome-wide significance.

We used the following stringent criteria for any exposure GWAS to be included in our analysis: (i) the GWAS had to report SNPs with p-values less than 5.0×10^−8^ for their association with a given exposure; (ii) these SNPs or their proxies (linkage disequilibrium R^2^ value >= 0.8) had to be present in both the exposure and outcome (PD) datasets; (iii) these SNPs were independent signals that were generated through a process called ‘clumping’. In order to ‘clump’, index SNPs were identified by ranking exposure associations from the smallest to largest p-value (but still with a cutoff value of p=5×10^−8^). Clumped SNPs were those in linkage disequilibrium (LD) with index SNPs (R^2^ threshold of 0.001) or within 10,000 kb physical distance. Hence, each index SNP represented a number of clumped SNPs that were all associated with or near to the index SNP, and the index SNPs were all independent of one another (according to the stringent parameters defined here). A total of 5,839 GWASes surpassed these criteria and were tested against the outcome. Then, we further expanded our filtering approach as follows: (iv) in order to use MR sensitivity analyses designed to identify pleiotropy, each GWAS had to include a minimum of ten associated SNPs; (v) the number of cases was > 250 for GWASes of a binary exposure or > 250 individuals for GWASes of a continuous exposure; and (vi) both the exposure and the outcome data were drawn from European populations. A total of 401 traits met our filtering criteria (7% of 5,839), consisting of 175 published GWASes and 226 unpublished GWASes from UK Biobank (UKB; www.ukbiobank.ac.uk). For UKB GWASes, some of the exposures are reported in an ordinal fashion, but treated as continuous when calculating betas for the effect allele at each SNP. This means that some of the effect estimates that arise are difficult to interpret quantitatively, both in the GWAS and in the subsequent MR analysis.

### Outcome data

Summary statistics from the largest, published PD GWAS meta-analysis involving 26,035 PD cases and 403,190 controls of European ancestry were used as the outcome data for the primary analysis ^11^. In this study there were 7,909,453 genotyped and imputed SNPs tested for association with PD with a mean allele frequency (MAF) > 3%. Recruitment and genotyping quality control procedures were described in the original report ^11^.

A newer PD GWAS includes a total of 37,688 cases, 1,417,791 controls and 18,618 ‘proxy cases’ from UKB (individuals that reported having a parent with PD) ^2^. However, there is substantial overlap in control subjects between each of the UKB exposures and the Nalls et al., 2019 meta-analysis, which can in turn lead to bias in causal effect estimates. For this reason, we repeated the analyses using only 5,851 clinically-diagnosed PD cases and 5,866 matched controls as the outcome, after excluding UKB samples and self-reported PD cases and controls. Finally, we used an earlier PD GWAS that included 13,708 cases and 95,282 controls as the outcome ^12^.

### Mendelian randomization analyses

Harmonization was undertaken to rule out strand mismatches and to ensure alignment of SNP effect sizes. Within each exposure GWAS, Wald ratios were calculated for each extracted SNP by dividing the per-allele log-odds ratio (or beta) of that variant in the PD GWAS data by the log-odds ratio (or beta) of the same variant in the exposure data.

First, the inverse-variance weighted (IVW) method was implemented to examine the relationship between the individual exposures and PD. In this method, the Wald ratio for each SNP was weighted according to its inverse variance and the effect estimates were meta-analysed using random effects. This approach is equivalent to plotting SNP-exposure/SNP-outcome associations on scatter plot and fitting a regression line (inverse-variance weighted regression), which is constrained to pass through the origin. The slope of the linear regression represents the pooled-effect estimate of the individual SNP Wald ratios^13^. For the purpose of demonstrating the use of this new platform, we used an false discovery rate (FDR) adjusted p-value of <0.05 to define exposures of interest as showing potential evidence of a causal effect. The IVW estimate is valid when the three core assumptions that underpin MR are upheld (see **Supplemental Material**). However, simulation studies show that up to 90% of MR analysis may be affected by pleiotropy, which in turn may bias the IVW estimate ^14^. Effects of pleiotropy for each analysis were studied by first looking for evidence of heterogeneity in the individual SNP Wald ratios and then undertaking a range of sensitivity analyses, each with different underlying assumptions.

Heterogeneity in the IVW estimates was tested using the Cochran’s Q test, quantified using the I^2^ statistic, and displayed in forest plots. Heterogeneity in the IVW estimate may indicate that alternative pathways exist from some of the SNPs to the outcome (known as horizontal pleiotropy), which can violate the third MR assumption ^15^, but as long as overall heterogeneity is balanced it does not necessarily bias the pooled IVW estimate ^16^. After calculation of the IVW estimates, three sensitivity analyses were applied to evaluate the core assumptions of MR and which rely on instruments containing multiple SNPs (in this case a minimum of 10 SNPs per exposure phenotype) ^17^.

MR-Egger was used to demonstrate evidence of a net unbalanced horizontal pleiotropic effect which might bias the IVW estimate. With MR-Egger, the regression line fitted to the data is not constrained to pass through the origin and a non-zero intercept indicates whether there is a net horizontal pleiotropic effect which may bias the IVW estimate ^18^. The weighted median (WM) MR method gives consistent effect estimates under the assumption that no more than 50% of the weight of the MR effect estimate comes from invalid (e.g. pleiotropic) SNPs, where weight is determined by the strength of their association with the exposure ^19^. Finally, IVW radial analysis was performed as a complementary method to account for SNPs acting as heterogeneous outliers and to determine the effect of resulting bias on the IVW estimate ^16^.

### Exploring directionality, single SNP effects and reverse causality

For effect estimate directionality, odds ratios were scaled on a standard deviation increase in genetic risk for the exposure from that population mean. We evaluated the possibility that the overall estimate was driven by a single SNP using leave-one-out (LOO) analyses for each of the phenotypic traits associated with PD. Finally, we tested for reverse causation by using SNPs tagging the independent loci described in the latest PD GWAS as exposure instrumental variables, and exposure GWASes as the outcomes. Note that this analysis measures the causal effect of genetic liability towards PD on each of the exposure traits included in the main analysis, which is independent of PD actually occurring (in a case-control setting such as this).

## RESULTS

The PD MR portal is hosted at https://pdgenetics.shinyapps.io/pdgenetics. Users can search for evidence that PD is causally affected by a broad range of exposures from 5,839 GWASes. Here, we explore some of the top results from our filtered analysis of 401 exposures to assist users in understanding how to interpret these data (see Figure 1 for flowchart of analysis). To our knowledge, there are no previous reports where a large-scale approach has been implemented to explore the causal contribution of multiple exposures to PD risk. Such approaches require appropriate multiple testing correction in order to avoid false-positive associations, which we have undertaken. That noted, at present most MR analyses are not sufficiently well-powered to detect truly causal associations after correcting for multiple testing, hence context should be applied when weighing the evidence for causality.

**Figure 1.**
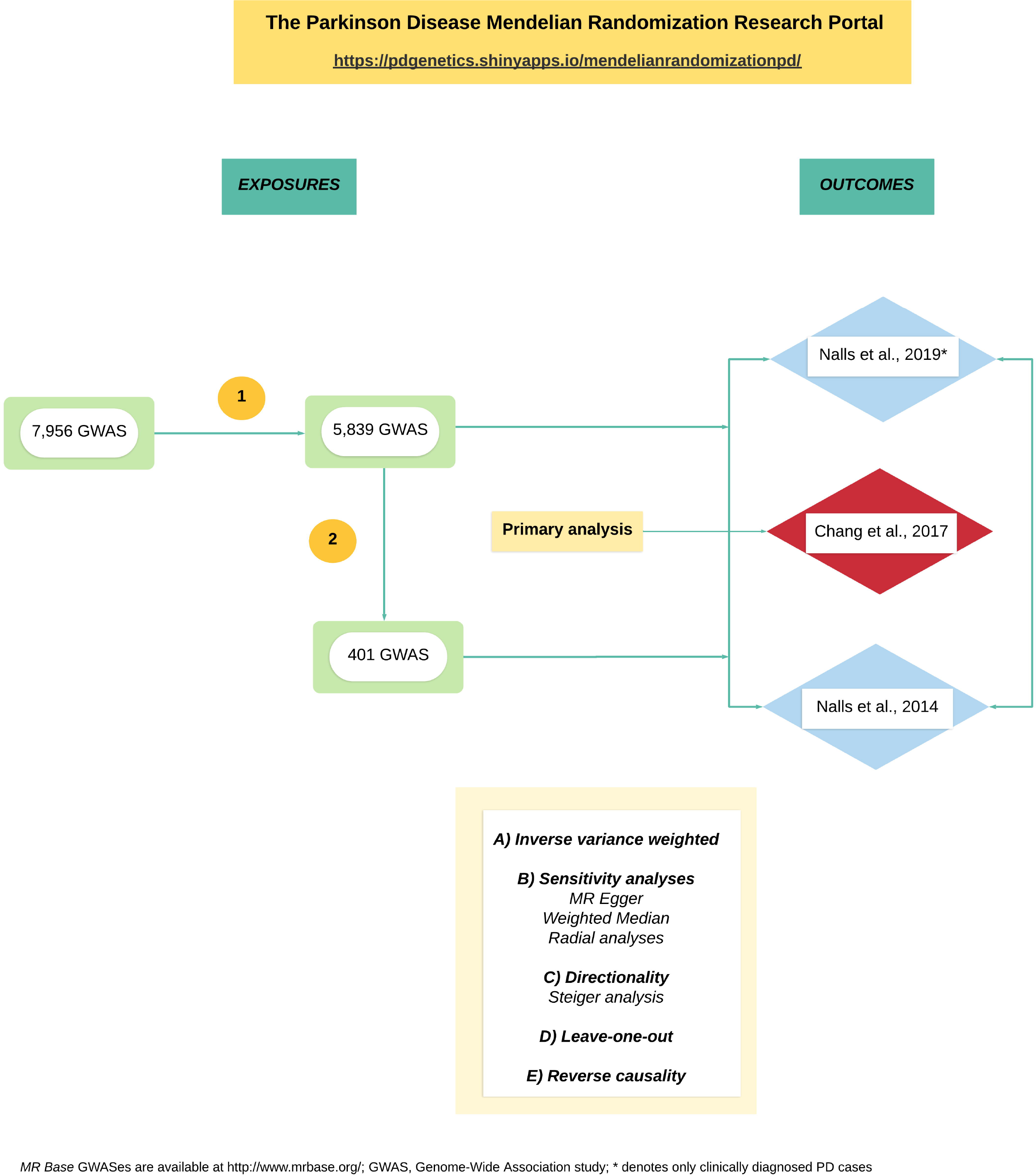
Flowchart of analysis. The PD MR Research Portal is an interactive tool where the user can explore causal associations across multiple traits. (1) The inclusion criteria used includes (i) GWASes with at least two associated SNPs with p-values less than 5.0×10^−8^, (ii) SNPs present in both the exposure and outcomes (Chang et al., 2017 as the primary analysis, Nalls et al., 2019 with only clinically diagnosed cases and Nalls et al., 2014) datasets or when not present their linkage-disequilibrium proxies (R2 value >= 0.8); and, (iii) independent SNPs (R2 < 0.001 with any other associated SNP within 10 Mb), considered as the most stringent clumping threshold used when performing MR analyses; (B) A total of 5,839 GWASes surpassed this criteria and were tested against the 3 outcomes. Then, we further expanded our filtering approach as follows: (iv) each GWAS had to include a minimum of 10 associated SNPs in order to use MR sensitivity analyses designed to identify pleiotropy, (v) the number of cases had to be > 250 for each GWAS of a given binary exposure or > 250 individuals for each GWAS of a given continuous trait; and (vi) the exposure and the outcome datasets were drawn from European populations. A total of 401 traits surpassed our filtering approach consisting of 175 published GWASes and 226 unpublished GWASes from UK Biobank (UKB; www.ukbiobank.ac.uk);

Of the 401 exposures that we included in this report, we found twelve exposures with potentially causal effects on PD (i.e. IWV FDR-adjusted p-values <0.05) (see Table 1 and **Figure S1** for forest plots displaying individual SNP-level estimates and pooled estimates). Of these exposures, eight were measures of adiposity that implied an inverse causal effect of increase adiposity and a lowering of PD risk. Four additional exposure traits met the FDR-adjusted p-value threshold for a possible causal effect: time spent watching television (inverse), tea drinking (positive), forced vital capacity (FVC; positive) and impedance of the right leg (positive). In each case, to explore the possibility that results were biased due to the violation of core MR assumptions, we looked for heterogeneity in the individual Wald ratios and performed the three additional MR sensitivity analyses: MR-Egger, WM and radial IVW (see Table 2).

**Table 1.** Mendelian randomization analyses of exposures with IVW FDR < 0.05 vs Chang et al., 2017

| Exposure | no. of SNPs | IVW |  |  |  | MR Egger |  |  | Weighted Median |  |  | Radial IVW |  |  |
| --- | --- | --- | --- | --- | --- | --- | --- | --- | --- | --- | --- | --- | --- | --- |
|  |  | beta | se | p-value | FDR | beta | se | p-value | beta | se | p-value | beta | se | p-value |
| Arm fat percentage (right) \|\| id:UKB-a:282 | 214 | -0.431 | 0.098 | 1.10E-05 | 0.005 | -0.657 | 0.324 | 0.044 | -0.372 | 0.143 | 0.009 | -0.454 | 0.079 | 3.35E-08 |
| Arm fat percentage (left) \|\| id:UKB-a:286 | 233 | -0.417 | 0.103 | 4.94E-05 | 0.010 | -0.790 | 0.333 | 0.018 | -0.201 | 0.137 | 0.143 | -0.434 | 0.077 | 5.41E-08 |
| Arm fat mass (right) \|\| id:UKB-a:283 | 248 | -0.276 | 0.072 | 1.32E-04 | 0.011 | -0.305 | 0.218 | 0.164 | -0.299 | 0.100 | 0.003 | -0.271 | 0.056 | 2.44E-06 |
| Leg fat percentage (right) \|\| id:UKB-a:274 | 228 | -0.482 | 0.126 | 1.33E-04 | 0.011 | -0.944 | 0.479 | 0.050 | -0.436 | 0.158 | 0.006 | -0.465 | 0.096 | 2.54E-06 |
| Whole body fat mass \|\| id:UKB-a:265 | 258 | -0.269 | 0.070 | 1.35E-04 | 0.011 | -0.362 | 0.220 | 0.102 | -0.241 | 0.102 | 0.018 | -0.281 | 0.056 | 7.95E-07 |
| Arm fat mass (left) \|\| id:UKB-a:287 | 248 | -0.279 | 0.074 | 1.58E-04 | 0.011 | -0.492 | 0.222 | 0.027 | -0.307 | 0.105 | 0.004 | -0.288 | 0.056 | 6.07E-07 |
| Trunk fat mass \|\| id:UKB-a:291 | 257 | -0.259 | 0.070 | 2.10E-04 | 0.012 | -0.438 | 0.227 | 0.055 | -0.256 | 0.094 | 0.007 | -0.255 | 0.054 | 4.37E-06 |
| Time spent watching television (TV) \|\| id:UKB-b:5192 | 104 | -0.775 | 0.211 | 2.40E-04 | 0.013 | -0.166 | 1.016 | 0.870 | -0.586 | 0.234 | 0.012 | -0.772 | 0.156 | 2.81E-06 |
| Trunk fat percentage \|\| id:UKB-a:290 | 216 | -0.307 | 0.085 | 3.20E-04 | 0.015 | -0.554 | 0.307 | 0.073 | -0.217 | 0.111 | 0.050 | -0.325 | 0.066 | 1.49E-06 |
| Forced vital capacity (FVC) Best measure \|\| id:UKB-a:232 | 179 | 0.441 | 0.128 | 5.80E-04 | 0.024 | 0.940 | 0.381 | 0.015 | 0.241 | 0.123 | 0.050 | 0.455 | 0.075 | 6.27E-09 |
| Tea intake \|\| id:UKB-b:6066 | 35 | 0.554 | 0.163 | 6.58E-04 | 0.025 | 0.710 | 0.334 | 0.041 | 0.543 | 0.223 | 0.015 | 0.570 | 0.143 | 2.97E-04 |
| Impedance of leg (right) \|\| id:UKB-a:270 | 294 | 0.220 | 0.066 | 9.17E-04 | 0.032 | 0.212 | 0.189 | 0.263 | 0.315 | 0.091 | 0.001 | 0.233 | 0.054 | 2.14E-05 |

id, specific code attributed to each trait by MR Base; se, standard error; No. of SNPs, number of SNPs; MR, Mendelian randomization; NA: not applicable due to limited no. of SNPs; IVW, Inverse Variance Weighted

**Table 2.**
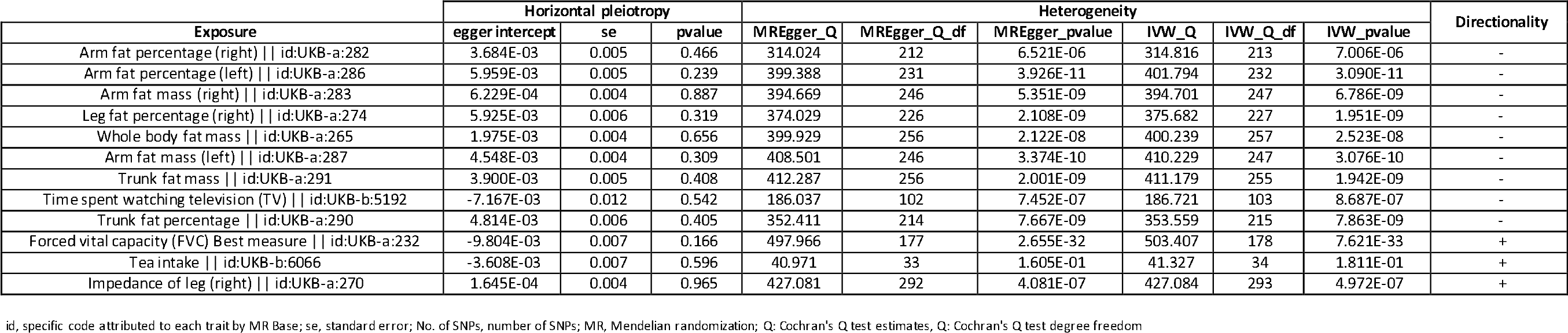
Heterogeneity, horizontal pleiotropy and directionality analyses for exposures with FDR < 0.05 vs Chang et al. 2017

The eight adiposity GWASes all contained >200 SNPs and were highly correlated with one another. The strongest causal effects were observed for arm fat percentage which was measured quantitatively using tissue impedance (https://biobank.ctsu.ox.ac.uk/crystal/field.cgi?id=23119). A unit increase in arm fat percentage (right and left) yielded a pooled odds ratio of 0.65 (95% CI 0.54-0.79, *FDR adjusted* p=0.005) and a pooled odds ratio of 0.66 (95% CI 0.54-0.79, *FDR adjusted* p=0.01) respectively. Although the individual Wald ratios showed significant heterogeneity (arm fat percentage (right) Q=314.8, p=7.01×10^−6^; arm fat percentage (left) Q=399.38, p=3.93×10^−11^), the sensitivity analyses did not suggest significant bias in the causal effect estimate from the IVW analysis (see Table 2). In general, all effect estimates for the adiposity traits supported a protective effect of increased adiposity on risk of PD (ORs ranged between 0.62 and 0.77) (see Table 1 and Table 2).

Another body composition exposure called ‘impedance of right leg’ surpassed the IVW FDR-adjusted p-value threshold. As described above, impedance is the method by which percentage fat is calculated, with higher impedance indicative of higher fat percentage (https://biobank.ctsu.ox.ac.uk/crystal/field.cgi?id=23107). Given the results for adiposity, we expected that a unit change in impedance would result in a causal lowering of PD risk. In contrast to the results for adiposity, the impedance exposure trait gave rise to an IVW OR 1.25 (95% CI 1.10-1.42, *FDR adjusted* p=0.032). There was significant heterogeneity in the individual Wald ratios (Q=428.4, p=7.25×10^−7^), but the sensitivity analyses did not suggest bias in the IVW effect estimate (see Table 2). We sought to explain why the direction of effect for impedance was different to that for percentage fat, when it was expected that it should be the same. Genetic correlations between impedance and percentage fat were run (see **Table S1**), and in all cases the genetic association between the two traits was negative.

The ‘tea drinking’ exposure was captured by an instrument containing 35 SNPs. The IVW OR was 1.74 (95% CI 1.26-2.39, *FDR adjusted* p=0.025) and there was no significant heterogeneity in the individual Wald ratios (Q=41.3, p=0.181). The three sensitivity analysis did not imply bias in the IVW estimate. Of note, coffee intake was not causally associated with increased risk of PD (OR 1.37, p=0.197), but similar to the ‘tea drinking’ exposure, the direction of the effect was also not consistent with the widely-accepted negative observational effect.

‘Time spent watching television’ was inversely linked to PD risk in the IVW analysis (OR 0.46, *FDR adjusted* p=0.013), suggesting that more time spent watching television caused a lower risk of PD. However, the MR Egger sensitivity analysis gave a very different pooled effect estimate (OR 0.85, p=0.870) suggesting that the IVW result may have been biased by directional pleiotropy. The intercept term can be used to test for net directional pleiotropy (here it was −0.007, p=0.542), but this test is generally underpowered. The difference in the slope from the IVW analysis and the MR Egger analysis is shown graphically in Figure S2 and suggests that, in the presence of directional pleiotropy, the IVW effect for ‘time spent watching television’ may be overestimated. From the other scatter and forest plots, it is clear that for the adiposity traits, that the slope (or magnitude of effect) the MR Egger regression is greater than the IVW slope (effect), which suggests that in the presence of directional pleiotropy the IVW may be underestimated.

‘Forced vital capacity’ also showed a positive causal effect on PD risk that was similarly observed in the sensitivity analyses (see Table 1 and Table 2). However, the effects appeared to be largely driven by two specific SNPs that are known to be pleiotropic for PD (see below).

The LOO analysis showed that none of the results described for the twelve exposures were being driven by a single SNP in each of the instruments (**Table S2**). The most precisely estimated Wald ratio for most of the adiposity exposures (7/8) came from a single SNP in the *FTO* gene (rs11642015), but dropping this SNP from the analyses did not affect the overall results. Similarly, the most precisely estimated Wald ratio in the impedance instrument was for a different SNP in the *FTO* gene (rs62048402), but again dropping this SNP did not affect the overall results. Importantly, and in support of the observations relating to negative genetic correlation between percentage fat and impedance described above, the direction of the Wald ratio for the *FTO* SNP in adiposity exposures was negative and for impedance was positive.

For tea drinking, the two Wald ratios with the greatest influence on the pooled effect estimates came from SNPs rs4410790 and rs2472297, which are located in the *AHR* and *CYP1A2* gene loci respectively, and are known to be strongly associated with caffeine-consumption behaviours ^20^. Leaving either SNP out from the analysis did not change the overall result, but the pooled effect estimate weakened when rs4410790 (*AHR*) was removed (the OR 1.74 changed to 1.61). In the ‘forced vital capacity’ analysis SNPs rs1991556 and rs13146142 were most precisely estimated and appeared to influence the magnitude of the causal effect. Closer examination revealed that rs1991556 is in the *MAPT* locus and rs13146142 is the *LCORL* locus, and both are known to be associated with PD, likely biasing the causal effect of FVC on PD ^12,2^.

The reverse causation analyses revealed no clear evidence that a liability towards PD was causally linked with any of the twelve exposures, but this analysis was restricted to only 18 of the 43 PD GWAS significant hits and may have been underpowered to detect reverse causal effects **(Table S3)**.

Of interest, the next seven exposures with the strongest associations in the IVW analysis, but not surpassing the FDR adjusted p-value threshold, were four adiposity traits (all showing a negative causal effect), current tobacco smoking (negative causal effect), increased alcohol intake (negative causal effect), and increased education (having a college or university degree; positive causal effect). The FDR adjusted p-values for each of these were <0.1 and unadjusted p-values were all <0.004.

Finally, we used clinically diagnosed cases (and controls) from the 2019 PD GWAS dataset as the outcome (**Table S4**). Given the smaller sample size, none of the MR analyses surpassed an FDR adjusted p-value, but the top hit was for a marker of adiposity (hip circumference). When an even earlier iteration of the PD GWAS was used (~13.5k cases and ~95k controls)^12^, there were 11 exposures that surpassed the FDR-adjusted p-value threshold and all 11 were related to adiposity (**Table S5**).

## Discussion

Here we present the PD MR research portal, hosted at https://pdgenetics.shinyapps.io/pdgenetics/. We envisage this being a valuable and evolving resource for the global research community, which will be updated as new data emerge. It should be used to provide evidence to support, and over time, evidence against, causality when undertaking observational studies or pursuing interventions aimed at reducing the risk of PD (hypothesis-driven research). Clearly there are too many associations presented in the portal to explore each one in detail. So, for the purpose of demonstrating the use of the tool, we undertook a data-driven approach to identify those exposures with the strongest causal signals (FDR adjusted IVW p-value <0.05).

We have previously reported an inverse causal association between BMI and risk of PD - a genetically-estimated 5kg/m^2^ higher BMI was associated with a reduced risk of PD (IVW OR=0.82) ^21^. Here we found further evidence to support an inverse relationship between increased adiposity and PD, given that eight of the top twelve exposures were measures of adiposity. These exposures were objectively and quantitatively ascertained and each GWAS in question identified >200 SNPs that were associated with increased adiposity. The results of the IVW analyses and all sensitivity analyses were broadly consistent, and effect across all traits was consistent with there being a protective effect of higher BMI and/or higher percentage fat on PD. Impedance gave rise to a causal effect opposite to that of increasing adiposity, which was initially unexpected. We have no reason to be believe that the scale upon which SNPs were associated with impedance was inverted. However, the difference in the direction of effect may be because the equation that links percentage fat and impedance requires adjustment for height and weight, which would not have been done routinely in the impedance GWAS ^22^.

Of note, although BMI specifically was not one of the top twelve exposures, the causal effect of BMI on PD was also negative in this analysis (OR 0.81; p=0.037) using the same BMI instrument that we have previously published about ^21,23^. In that earlier analysis, we included simulations that indicated the protective effect was unlikely to be explained by survival bias (that is, people with higher BMI dying of other diseases before they would usually be diagnosed with PD). The relationship between BMI and adiposity with PD clearly warrants further study. It is not clear at what point across the BMI scale protection against PD is conveyed since all we have observed is an averaged linear association between BMI (and other markers of adiposity) and PD.

Tea drinking emerged as being causally linked to PD in that genetically-estimated tea drinking status appeared to increase the risk of PD. This result was largely unexpected because the observational study associations between caffeine-containing drinks (both coffee and tea) have almost always suggested a negative association between caffeine and PD ^4,24^. This has meant that caffeine has been explored as a potentially therapeutic option for PD ^25^. It is not clear what would account for observing consistently negative observational associations between caffeine and PD, and a potentially detrimental effects of caffeine on PD in MR analyses, but the effect for coffee drinking in this analysis was also not in favour of a true protective effect. The remaining two exposures (time spent watching television and forced vital capacity) that appeared to be causally-linked to PD should be regarded with additional caution. Television watching appeared to fail in one of the sensitivity analyses and the effect of forced vital capacity appeared to be largely driven by two SNPs that are known to be associated with PD.

Data sharing restrictions relating to some of the data in PD GWASes meant that the full summary statistics cannot be released to libraries such as *MR Base* and the presentation of the results in the PD MR portal has been tailored to suit our specific aims. Alongside a standard method used to pool ratio estimates from individual SNPs (the IVW method), we also demonstrated the use of three sensitivity analyses that provide more valid estimates when core instrumental variable assumptions have been violated (namely the MR-Egger, WM and radial methods) ^19,16^. These methods will allow researchers to further appraise the validity of the instruments presented and whether IVW estimates may be biased. In the current version of the portal, we have provided the opportunity of undertaking analyses with three different PD outcome datasets. There was little qualitative difference in the ‘top MR hits’ across the three iterations of the PD outcome data, particularly when comparing the larger datasets. The portal also contains a huge number of exposures that have been curated by *MR Base*. However, as highlighted in this report and in a warning message in the portal, we advise caution with the interpretation of results for exposure traits that do not surpass considered filtering criteria (for our criteria this was 5,438 out of 5,839 or 93% of exposures). These included GWASes for which the case numbers were small and the number of variants in the exposure instruments was too few to permit the use of the sensitivity analyses.

## Limitations

Here we have presented a hypothesis-free approach using two-sample MR to study causal associations between PD and a range of exposures. In most instances, the variation in a given exposure that is explained by genome-wide significant GWAS signals is small (i.e 0.5%-8%), meaning that sample sizes often need to be extremely large to detect causal effects using this design. The resulting p-values that arise from MR hypothesis tests are often not small and in the various sensitivity analyses, often border on or exceed nominal statistical significance. Given the approach, traditional methods of adjusting for multiple comparisons (i.e. Bonferroni or false discovery rate correction) render many potentially causal associations obsolete and limit the conclusions that one can draw. While we have observed further evidence to support the role of increased adiposity on lower risk of PD, in general we recommend using this tool to confirm or add to evidence to associations identified in observational studies or towards a particular intervention. In any MR analysis, it is generally recommended that emphasis should be placed on the confidence intervals and consistency of point estimates across the IVW estimate and the sensitivity analyses, rather than just the p-values *per se*. In addition, we recommend the calculation of F-statistics for determining instrument strength and post-hoc power calculations, particularly in the instance of null results ^26^. This is particularly important when making claims of no evidence to support a causal association (such as in some of the reverse causation analyses presented in this manuscript). These additions and others are planned for future versions of the portal.

It is important to remember that for all analyses presented here, the outcome is ‘risk of PD’ because the outcome data come from a GWAS of PD cases versus controls. In order to make causal inferences about the effect of various exposures on disease progression, one would need to have an outcome GWAS of PD progression. As mentioned in the Methods, an important source of bias in MR studies can occur when there is overlap in the control groups in the exposure and outcome data. For this reason, all UK Biobank data was removed from the outcome PD GWAS summary statistics.

In summary, we present a new portal for use by the research community that will help assess causality where observational associations exist and prioritize (or de-prioritize) interventions aimed at reducing risk of PD overall. We think that it will be of value and will continue to evolve over time.

## Funding agencies

This research was supported in part by the Intramural Research Program of the National Institutes of Health (National Institute on Aging, National Institute of Neurological Disorders and Stroke; project numbers: project numbers 1ZIA-NS003154, Z01-AG000949-02 and Z01-ES101986. The QMUL Preventive Neurology Unit is funded by the Barts Charity: grant reference number MGU0364. George Davey Smith and Debbie Lawlor work in the Medical Research Council Integrative Epidemiology Unit at the University of Bristol, which is supported by the Medical Research Council (MC_UU_00011/1). University College London Hospitals and University College London receive support from the Department of Health’s National Institute for Health Research (NIHR) Biomedical Research Centres (BRC). Debbie Lawlor and Nicholas Wood are NIHR senior Investigators.

## AUTHORS’ ROLES

1) **Research project:** A. Conception (AJN, SB, JJ), B. Organization (AJN, SB, JJ), C. Execution (AJN, SB, JJ)

2) **Data generation:** A. Experimental (IPDGC).

3) **Statistical Analysis:** A. Design (AJN, SB, JJ), B. Execution (AJN, SB, JJ)

4) **Manuscript Preparation:** A. Writing of the first draft (AJN, SB), B. Review and Critique (all authors)

## ACKNOWLEDGEMENTS

We thank the research participants for their contribution to the GWAS used here. We would like to thank Zhihong Zhu (Queensland) who contributed to the discussion on MR methods. We thank the following members of the 23andMe Research Team: Michelle Agee, Babak Alipanahi, Adam Auton, Robert K. Bell, Katarzyna Bryc, Sarah L. Elson, Pierre Fontanillas, Nicholas A. Furlotte, Barry Hicks, Karen E. Huber, Ethan M. Jewett, Yunxuan Jiang, Aaron Kleinman, Keng-Han Lin, Nadia K. Litterman, Jennifer C. McCreight, Matthew H. McIntyre, Kimberly F. McManus, Joanna L. Mountain, Elizabeth S. Noblin, Carrie A.M. Northover, Steven J. Pitts, G. David Poznik, J. Fah Sathirapongsasuti, Janie F. Shelton, Suyash Shringarpure, Chao Tian, Joyce Y. Tung, Vladimir Vacic, Xin Wang, Catherine H. Wilson.

## REFERENCES

1. Billingsley KJ, Bandres-Ciga S, Saez-Atienzar S, Singleton AB. Genetic risk factors in Parkinson’s disease. Cell Tissue Res. 2018;373(1):9–20.

2. Nalls MA, Blauwendraat C, Vallerga CL, et al. Parkinson’s disease genetics: identifying novel risk loci, providing causal insights and improving estimates of heritable risk. bioRxiv. August 2018:388165. doi:10.1101/388165

3. Wray NR, Yang J, Goddard ME, Visscher PM. The genetic interpretation of area under the ROC curve in genomic profiling. PLoS Genet. 2010;6(2):e1000864.

4. Noyce AJ, Bestwick JP, Silveira-Moriyama L, et al. Meta-analysis of early nonmotor features and risk factors for Parkinson disease. Ann Neurol. 2012;72(6):893–901.

5. Berg D, Postuma RB, Adler CH, et al. MDS research criteria for prodromal Parkinson’s disease. Mov Disord. 2015;30(12):1600–1611.

6. Collier TJ E al. Aging and Parkinson’s disease: Different sides of the same coin? - PubMed - NCBI. https://www.ncbi.nlm.nih.gov/pubmed/28520211. Accessed March 12, 2019.

7. Reeve A E al. Ageing and Parkinson’s disease: why is advancing age the biggest risk factor? - PubMed - NCBI. https://www.ncbi.nlm.nih.gov/pubmed/24503004. Accessed March 12, 2019.

8. Smith GD. Epidemiology, epigenetics and the “Gloomy Prospect”: embracing randomness in population health research and practice. Int J Epidemiol. 2011;40(3):537–562.

9. Lawlor DA, Harbord RM, Sterne JAC, Timpson N, Davey Smith G. Mendelian randomization: using genes as instruments for making causal inferences in epidemiology. Stat Med. 2008;27(8):1133–1163.

10. Hemani G, Zheng J, Elsworth B, et al. The MR-Base platform supports systematic causal inference across the human phenome. Elife. 2018;7. doi:10.7554/eLife.34408

11. Chang D, Nalls MA, Hallgrímsdóttir IB, et al. A meta-analysis of genome-wide association studies identifies 17 new Parkinson’s disease risk loci. Nat Genet. 2017;49(10):1511–1516.

12. Nalls MA, Pankratz N, Lill CM, et al. Large-scale meta-analysis of genome-wide association data identifies six new risk loci for Parkinson’s disease. Nat Genet. 2014;46(9):989–993.

13. Burgess S, Small DS, Thompson SG. A review of instrumental variable estimators for Mendelian randomization. Stat Methods Med Res. 2017;26(5):2333–2355.

14. Hemani G, Bowden J, Haycock PC, et al. Automating Mendelian randomization through machine learning to construct a putative causal map of the human phenome. doi:10.1101/173682

15. Davey Smith G, Hemani G. Mendelian randomization: genetic anchors for causal inference in epidemiological studies. Hum Mol Genet. 2014;23(R1):R89–R98.

16. Bowden J, Spiller W, Del Greco M F, et al. Improving the visualization, interpretation and analysis of two-sample summary data Mendelian randomization via the Radial plot and Radial regression. Int J Epidemiol. 2018;47(4):1264–1278.

17. Hemani G, Bowden J, Davey Smith G. Evaluating the potential role of pleiotropy in Mendelian randomization studies. Hum Mol Genet. 2018;27(R2):R195–R208.

18. Bowden J, Davey Smith G, Burgess S. Mendelian randomization with invalid instruments: effect estimation and bias detection through Egger regression. Int J Epidemiol. 2015;44(2):512–525.

19. Bowden J, Davey Smith G, Haycock PC, Burgess S. Consistent Estimation in Mendelian Randomization with Some Invalid Instruments Using a Weighted Median Estimator. Genet Epidemiol. 2016;40(4):304–314.

20. Coffee and Caffeine Genetics Consortium, Cornelis MC, Byrne EM, et al. Genome-wide meta-analysis identifies six novel loci associated with habitual coffee consumption. Mol Psychiatry. 2015;20(5):647–656.

21. Noyce AJ, Kia DA, Hemani G, et al. Estimating the causal influence of body mass index on risk of Parkinson disease: A Mendelian randomisation study. PLoS Med. 2017;14(6):e1002314.

22. Jebb SA, Cole TJ, Doman D, Murgatroyd PR, Prentice AM. Evaluation of the novel Tanita body-fat analyser to measure body composition by comparison with a four-compartment model. Br J Nutr. 2000;83(2):115–122.

23. Locke AE, Kahali B, Berndt SI, et al. Genetic studies of body mass index yield new insights for obesity biology. Nature. 2015;518(7538):197–206.

24. Qi H, Li S. Dose-response meta-analysis on coffee, tea and caffeine consumption with risk of Parkinson’s disease. Geriatr Gerontol Int. 2014;14(2):430–439.

25. Postuma RB, Anang J, Pelletier A, et al. Caffeine as symptomatic treatment for Parkinson disease (Café-PD). Neurology. 2017;89(17):1795–1803. doi:10.1212/wnl.0000000000004568

26. Brion M-JA, Shakhbazov K, Visscher PM. Calculating statistical power in Mendelian randomization studies. Int J Epidemiol. 2013;42(5):1497–1501.

