## Supplemental information for "The Parkinson’s Disease Mendelian Randomization Research Portal"

* Equal contribution

**Figure S1.** Forest plots showing point estimates of the exposures of interest…………………….………………………….………………… 1

**Figure S2.** Funnel plots showing point estimates of the exposures of interest………………………………………………………………… 14

**Table S1.** Genetic correlations between fat related traits and impedance related traits……………………………………………….. 27

**Table S2**. Leave-one-out analyses of exposures of interest…………………………………………………………………………………………… 28

**Table S3.** Reverse causality analyses……………………………………………………………………………………………………………………….. 95

**Table S4.** Mendelian Randomization analyses of exposures significantly linked to PD at IVW p-value < 0.05 vs Nalls et al., 2019 (only clinically diagnosed cases)……………………………..……………………………………………………………………………………….. 97

**Table S5.** Mendelian Randomization analyses of exposures significantly linked to PD at IVW p-value < 0.05 vs Nalls et al., 2014……………………………………………………………………………………………………………………………………………………………..………………….. 98

Mendelian Randomization assumptions…………………………………………………………………………………………………………….. 99

Consortia members…………………………………………………………………………………………………………………………………………… 100

### Figure S1. Forest plots showing point estimates of the exposures of interest

(The trait appears at the bottom of each panel)

Within each panel, the black points represent the causal estimate (log-odds ratio) of each SNP on the risk of developing PD. Red points represent the causal estimate when combining all SNPs together, using the inverse variance weighted and MR Egger methods. Horizontal lines denote 95% confidence intervals.

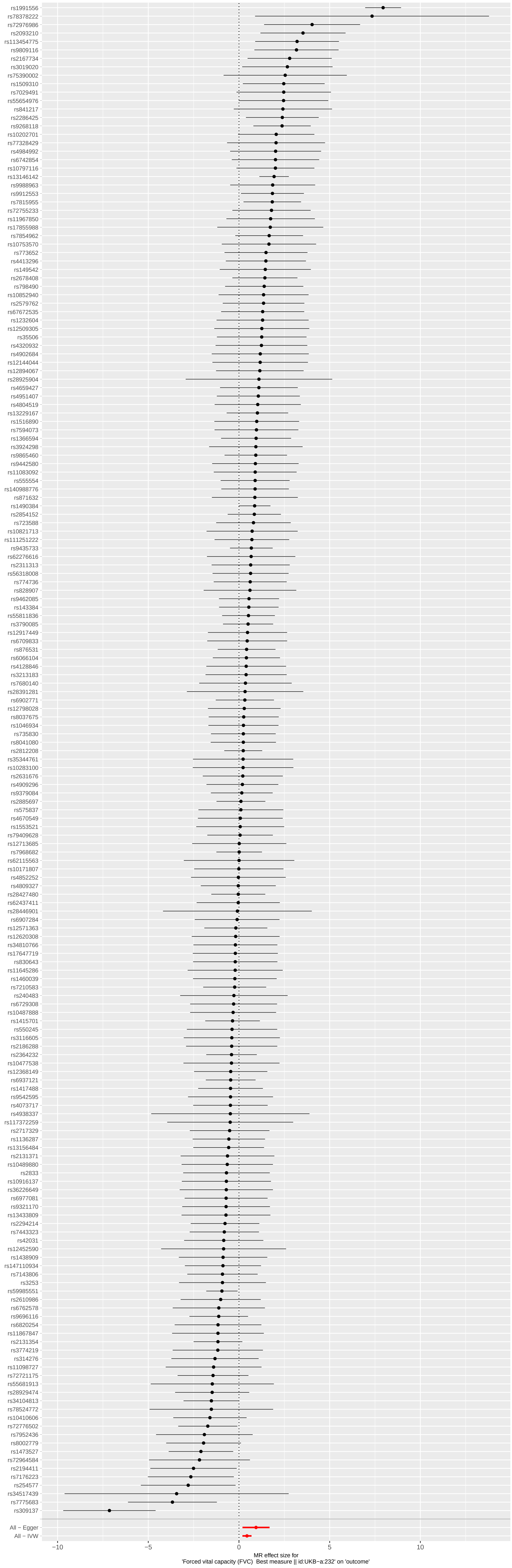
rs1991556
rs78378222
rs72976986
rs2093210
rs113454775
rs9809116
rs2167734
rs3019020
rs75390002
rs1509310
rs7029491
rs55654976
rs841217
rs2286425
rs9268118
rs10202701
rs77328429
rs4984992
rs6742854
rs10797116
rs13146142
rs9988963
rs9912553
rs7815955
rs72755233
rs11967850
rs17855988
rs7854962
rs10753570
rs773652
rs4413296
rs149542
rs2678408
rs798490
rs10852940
rs2579762
rs67672535
rs1232604
rs12509305
rs35506
rs4320932
rs4902684
rs12144044
rs12894067
rs28925904
rs4659427
rs4951407
rs4804519
rs13229167
rs1516890
rs7594073
rs1366594
rs3924298
rs9865460
rs9442580
rs11083092
rs555554
rs140988776
rs871632
rs1490384
rs2854152
rs723588
rs10821713
rs111251222
rs9435733
rs62276616
rs2311313
rs56318008
rs774736
rs828907
rs9462085
rs143384
rs55811836
rs3790085
rs12917449
rs6709833
rs876531
rs6066104
rs4128846
rs3213183
rs7680140
rs28391281
rs6902771
rs12798028
rs8037675
rs1046934
rs735830
rs8041080
rs2812208
rs35344761
rs10283100
rs2631676
rs4909296
rs9379084
rs2885697
rs575837
rs4670549
rs1553521
rs79409628
rs12713685
rs7968682
rs62115563
rs10171807
rs4852252
rs4809327
rs28427480
rs62437411
rs28446901
rs6907284
rs12571363
rs12620308
rs34810766
rs17647719
rs830643
rs11645286
rs1460039
rs7210583
rs240483
rs6729308
rs10487888
rs1415701
rs550245
rs3116605
rs2186288
rs2364232
rs10477538
rs12368149
rs6937121
rs1417488
rs9542595
rs4073717
rs4938337
rs117372259
rs2717329
rs1136287
rs13156484
rs2131371
rs10489880
rs2833
rs10916137
rs36226649
rs6977081
rs9321170
rs13433809
rs2294214
rs7443323
rs42031
rs12452590
rs1438909
rs147110934
rs7143806
rs3253
rs59985551
rs2610986
rs6762578
rs9696116
rs6820254
rs11867847
rs2131354
rs3774219
rs314276
rs11098727
rs72721175
rs55681913
rs28929474
rs34104813
rs78524772
rs10410606
rs72776502
rs7952436
rs8002779
rs1473527
rs72964584
rs2194411
rs7176223
rs254577
rs34517439
rs7775683
rs309137
All − Egger
All − IVW
−10
−5
0
5
10
MR effect size for
'Forced vital capacity (FVC)  Best measure || id:UKB−a:232' on 'outcome'

3

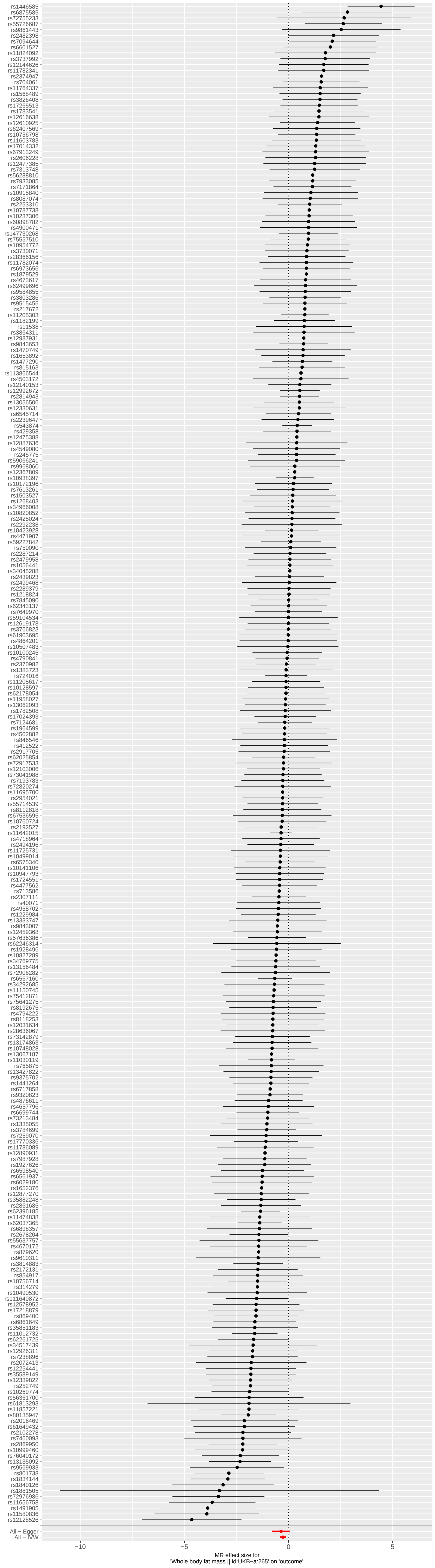
All − Egger
All − IVW
−10
−5
0
5
MR effect size for
'Whole body fat mass || id:UKB−a:265' on 'outcome'

4

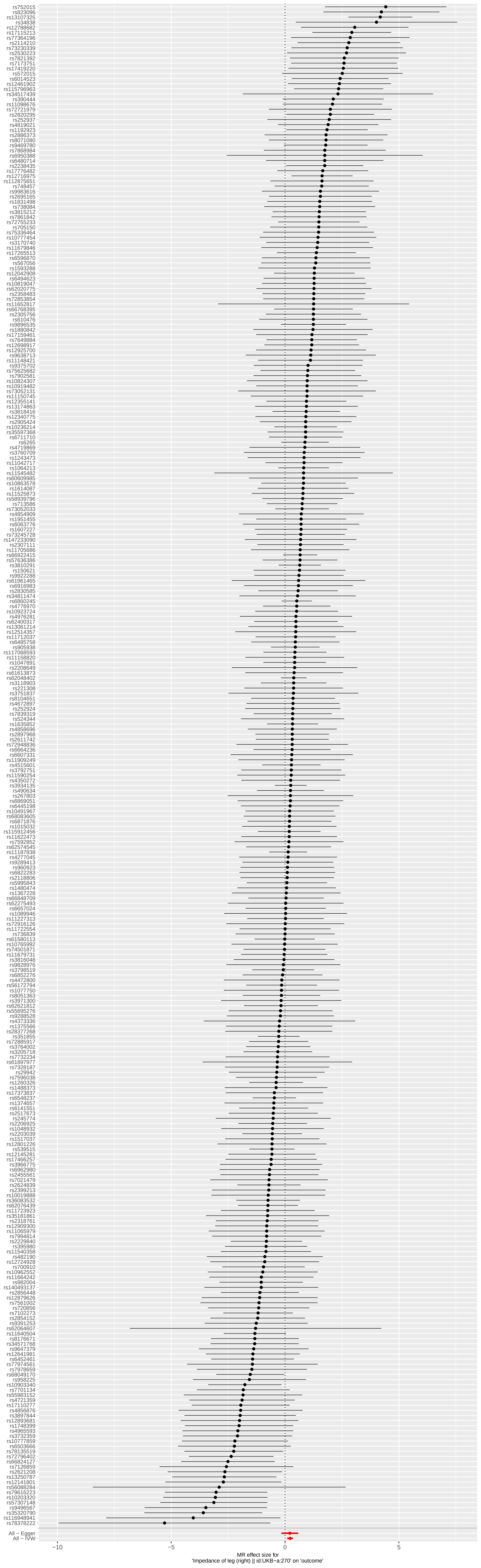
All − Egger
All − IVW
-10
-5
0
5
MR effect size for
'Impedance of leg (right) || id:UKB-a:270' on 'outcome'

5

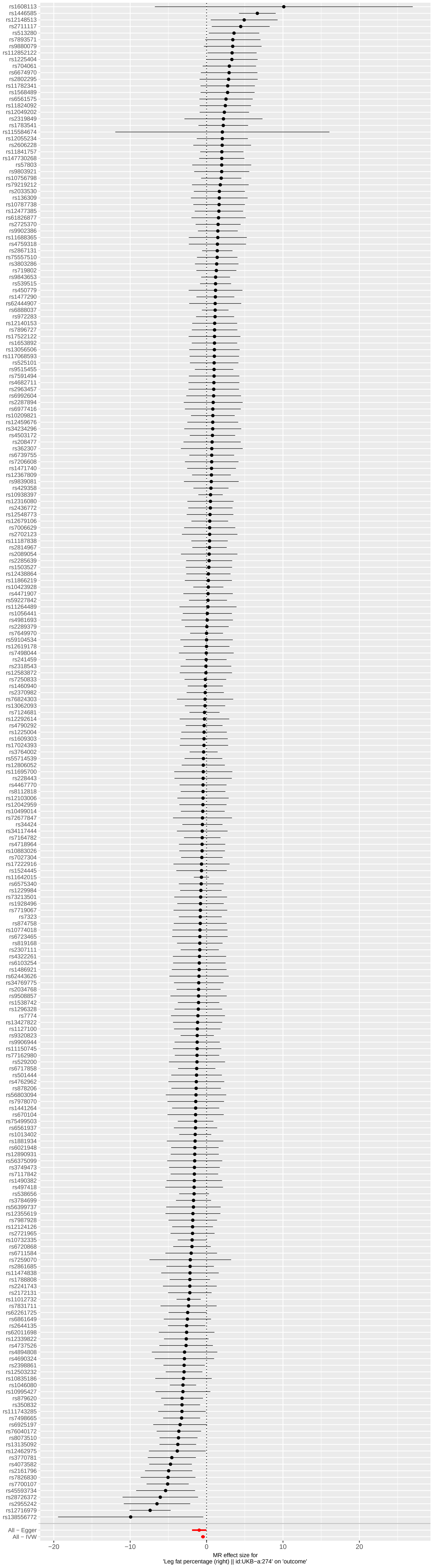
rs1608113
rs1446585
rs12148513
rs2711117
rs513280
rs7893571
rs9880079
rs112852122
rs1225404
rs704061
rs6674970
rs2802295
rs11782341
rs1568489
rs6561575
rs11824092
rs12049202
rs2319849
rs1783541
rs115584674
rs12055234
rs2606228
rs11841757
rs147730268
rs57803
rs9803921
rs10756798
rs79219212
rs2033530
rs136309
rs10787738
rs12477385
rs61826877
rs2725370
rs9902386
rs11688365
rs4759318
rs2867131
rs75557510
rs3803286
rs719802
rs9843653
rs539515
rs450779
rs1477290
rs62444907
rs6888037
rs972283
rs12140153
rs7896727
rs17522122
rs1653892
rs13056506
rs117068593
rs525101
rs9515455
rs7591494
rs4682711
rs2963457
rs6992604
rs2287894
rs6977416
rs10209821
rs12459676
rs34234296
rs4503172
rs208477
rs362307
rs6739755
rs7206608
rs1471740
rs12367809
rs9839081
rs429358
rs10938397
rs12316080
rs2436772
rs12548773
rs12679106
rs7006629
rs2702123
rs11187838
rs2814967
rs2089054
rs2285639
rs1503527
rs12438864
rs11866219
rs10423928
rs4471907
rs59227842
rs11264489
rs1056441
rs4981693
rs2289379
rs7649970
rs59104534
rs12619178
rs7498044
rs241459
rs2318543
rs12583872
rs7250833
rs1460940
rs2370982
rs76824303
rs13062093
rs7124681
rs12292614
rs4790292
rs1225004
rs1609303
rs17024393
rs3764002
rs55714539
rs12806052
rs11695700
rs228443
rs4467770
rs8112818
rs12103006
rs12042959
rs10499014
rs72677847
rs34424
rs34117444
rs7164782
rs4718964
rs10883026
rs7027304
rs17222916
rs1524445
rs11642015
rs6575340
rs1229984
rs73213501
rs1928496
rs7719067
rs7323
rs874758
rs10774018
rs6723465
rs819168
rs2307111
rs4322261
rs6103254
rs1486921
rs62443626
rs34769775
rs2034768
rs9508857
rs1538742
rs1296328
rs7774
rs13427822
rs1127100
rs9320823
rs9906944
rs11150745
rs77162980
rs529200
rs6717858
rs501444
rs4762962
rs878206
rs56803094
rs7978070
rs1441264
rs670104
rs75499503
rs6561937
rs1013402
rs1881934
rs6021948
rs12890931
rs56375099
rs3749473
rs7117842
rs1490382
rs497418
rs538656
rs3784699
rs56399737
rs12355619
rs7987928
rs12124126
rs2721965
rs10732335
rs6720868
rs6711594
rs7259070
rs2861685
rs11474838
rs1788808
rs2241743
rs2172131
rs11012732
rs7831711
rs62261725
rs6861649
rs2644135
rs62011698
rs12339822
rs4737526
rs4894808
rs4690324
rs2398861
rs12503232
rs10835186
rs1046080
rs10995427
rs879620
rs350832
rs111743285
rs7498665
rs6925197
rs76040172
rs8073510
rs13135092
rs12462975
rs3770781
rs4073582
rs2161796
rs7826830
rs7700107
rs45593734
rs28726372
rs2955242
rs12716979
rs138556772
All − Egger
All − IVW
−20
−10
0
10
20
MR effect size for
'Leg fat percentage (right) || id:UKB−a:274' on 'outcome'

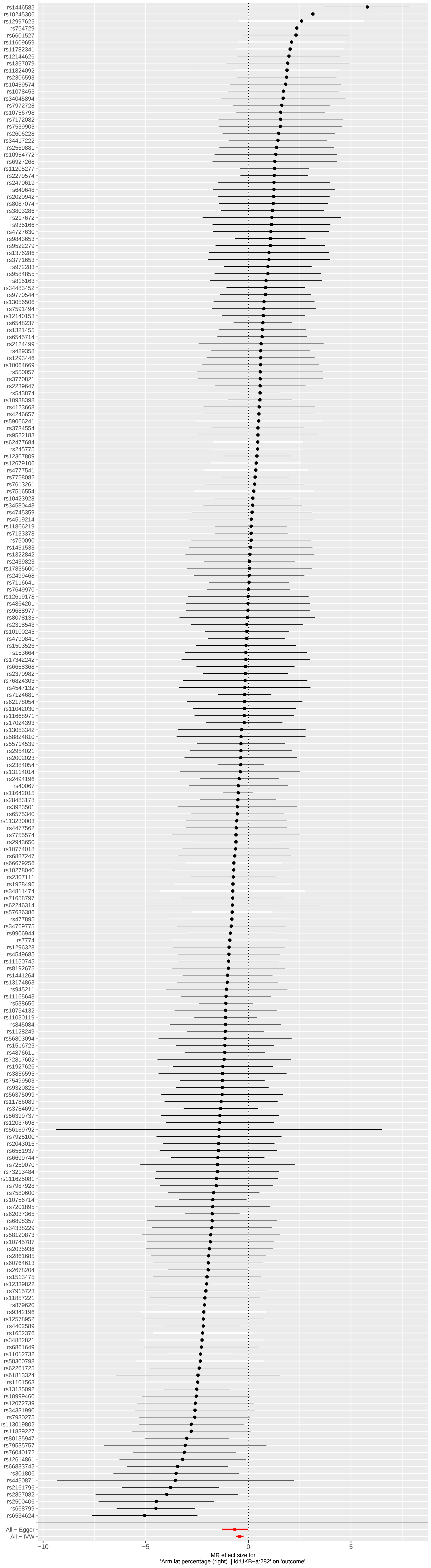
rs1446585
rs10245306
rs12997625
rs764729
rs6601527
rs11609659
rs11782341
rs12144626
rs1357079
rs11824092
rs2306593
rs10459574
rs1078455
rs34045894
rs7972728
rs10756798
rs7172082
rs7539903
rs2606228
rs34417222
rs2569881
rs10954772
rs6927268
rs11205277
rs2279574
rs2470619
rs649648
rs2020942
rs8087074
rs3803286
rs217672
rs935166
rs4727630
rs9843653
rs9522279
rs1376286
rs3771653
rs972283
rs9584855
rs815163
rs34483452
rs9770544
rs13056506
rs7591494
rs12140153
rs6548237
rs1321455
rs6545714
rs2124499
rs429358
rs1293446
rs10064669
rs550057
rs3770821
rs2239647
rs543874
rs10938398
rs4123668
rs4246657
rs59066241
rs3734554
rs9522183
rs62477684
rs245775
rs12367809
rs12679106
rs4777541
rs7758082
rs7613261
rs7516554
rs10423928
rs34580448
rs4745359
rs4519214
rs11866219
rs7133378
rs750090
rs1451533
rs1322842
rs2439823
rs17835600
rs2499468
rs7116641
rs7649970
rs12619178
rs4864201
rs9688977
rs8078135
rs2318543
rs10100245
rs4790841
rs1503526
rs153664
rs17342242
rs6658368
rs2370982
rs76824303
rs4547132
rs7124681
rs62178054
rs11042030
rs11668971
rs17024393
rs13053342
rs58824810
rs55714539
rs2954021
rs2002023
rs2384054
rs13114014
rs2494196
rs40067
rs11642015
rs28483178
rs3923501
rs6575340
rs113230003
rs4477562
rs7755574
rs2943650
rs10774018
rs6887247
rs66679256
rs10278040
rs2307111
rs1928496
rs34811474
rs71658797
rs62246314
rs57636386
rs477895
rs34769775
rs9906944
rs7774
rs1296328
rs4549685
rs11150745
rs8192675
rs1441264
rs13174863
rs945211
rs11165643
rs538656
rs10754132
rs11030119
rs845084
rs1128249
rs56803094
rs1516725
rs4876611
rs72817602
rs1927626
rs3856595
rs75499503
rs9320823
rs56375099
rs11786089
rs3784699
rs56399737
rs12037698
rs56169792
rs7925100
rs2043016
rs6561937
rs6699744
rs7259070
rs73213484
rs111625081
rs7987928
rs7580600
rs10756714
rs7201895
rs62037365
rs6898357
rs34338229
rs58120873
rs10745787
rs2035936
rs2861685
rs60764613
rs2678204
rs1513475
rs12339822
rs7915723
rs11857221
rs879620
rs9342196
rs12578952
rs4402589
rs1652376
rs34882821
rs6861649
rs11012732
rs58360798
rs62261725
rs61813324
rs1101563
rs13135092
rs10999460
rs12072739
rs34331990
rs7930275
rs113019802
rs11839227
rs80135947
rs79535757
rs76040172
rs12614861
rs66833742
rs301806
rs4450871
rs2161796
rs2857082
rs2500406
rs668799
rs6534624
All − Egger
All − IVW
−10
−5
0
5
MR effect size for
'Arm fat percentage (right) || id:UKB−a:282' on 'outcome'

7

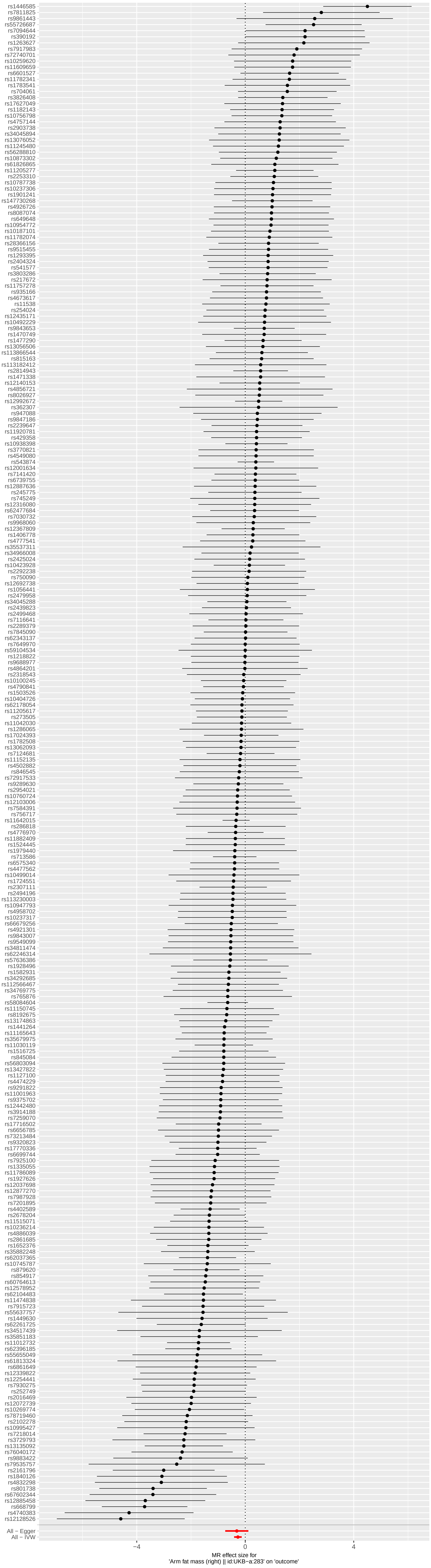
rs1446585
rs7811825
rs9861443
rs55726687
rs7094644
rs390192
rs1263627
rs7917983
rs72740701
rs10259620
rs11609659
rs6601527
rs11782341
rs1783541
rs704061
rs3826408
rs17627049
rs1182143
rs10756798
rs4757144
rs2903738
rs34045894
rs13076052
rs11245480
rs56288810
rs10873302
rs61826865
rs11205277
rs2253310
rs10787738
rs10237306
rs1901241
rs147730268
rs4926726
rs8087074
rs649648
rs10954772
rs10187101
rs11782074
rs28366156
rs9515455
rs1293395
rs2404324
rs541577
rs3803286
rs217672
rs11757278
rs935166
rs4673617
rs11538
rs254024
rs12435171
rs10492229
rs9843653
rs1470749
rs1477290
rs13056506
rs113866544
rs815163
rs113182412
rs2814943
rs1471338
rs12140153
rs4856721
rs8026927
rs12992672
rs362307
rs947088
rs9847186
rs2239647
rs11920781
rs429358
rs10938398
rs3770821
rs4549080
rs543874
rs12001634
rs7141420
rs6739755
rs12887636
rs245775
rs745249
rs12316080
rs62477684
rs7030732
rs9968060
rs12367809
rs1406778
rs4777541
rs35537311
rs34966008
rs2425024
rs10423928
rs2292238
rs750090
rs12692738
rs1056441
rs2479958
rs34045288
rs2439823
rs2499468
rs7116641
rs2289379
rs7845090
rs62343137
rs7649970
rs59104534
rs1218822
rs9688977
rs4864201
rs2318543
rs10100245
rs4790841
rs1503526
rs10404726
rs62178054
rs11205617
rs273505
rs11042030
rs1286065
rs17024393
rs1782508
rs13062093
rs7124681
rs11152135
rs4502882
rs846545
rs72917533
rs9289630
rs2954021
rs10760724
rs12103006
rs7584391
rs756717
rs11642015
rs286818
rs4776970
rs11882409
rs1524445
rs1979440
rs713586
rs6575340
rs4477562
rs10499014
rs1724551
rs2307111
rs2494196
rs113230003
rs10947793
rs4958702
rs10237317
rs66679256
rs4921301
rs9843007
rs9549099
rs34811474
rs62246314
rs57636386
rs1928496
rs1582931
rs34292685
rs112566467
rs34769775
rs765876
rs58084604
rs11150745
rs8192675
rs13174863
rs1441264
rs11165643
rs35679975
rs11030119
rs1516725
rs845084
rs56803094
rs13427822
rs1127100
rs4474229
rs9291822
rs11001963
rs9375702
rs12442480
rs3914188
rs7259070
rs17716502
rs6656785
rs73213484
rs9320823
rs17770336
rs6699744
rs7925100
rs1335055
rs11786089
rs1927626
rs12037698
rs12877270
rs7987928
rs7201895
rs4402589
rs2678204
rs11515071
rs10236412
rs4886039
rs2861685
rs1652376
rs35882248
rs62037365
rs10745787
rs879620
rs854917
rs60764613
rs12578952
rs62104483
rs11474838
rs7915723
rs55637757
rs1449630
rs62261725
rs34517439
rs35851183
rs11012732
rs62396185
rs55655049
rs61813324
rs6861649
rs12339822
rs12254441
rs7930275
rs252749
rs2016469
rs12072739
rs10269774
rs78719460
rs2102278
rs10995427
rs7218014
rs3729793
rs13135092
rs76040172
rs9883422
rs79535757
rs2161796
rs1840126
rs4832298
rs801738
rs67602344
rs12885458
rs668799
rs4740383
rs12128526
All − Egger
All − IVW
−4
0
4
MR effect size for
'Arm fat mass (right) || id:UKB−a:283' on 'outcome'

8

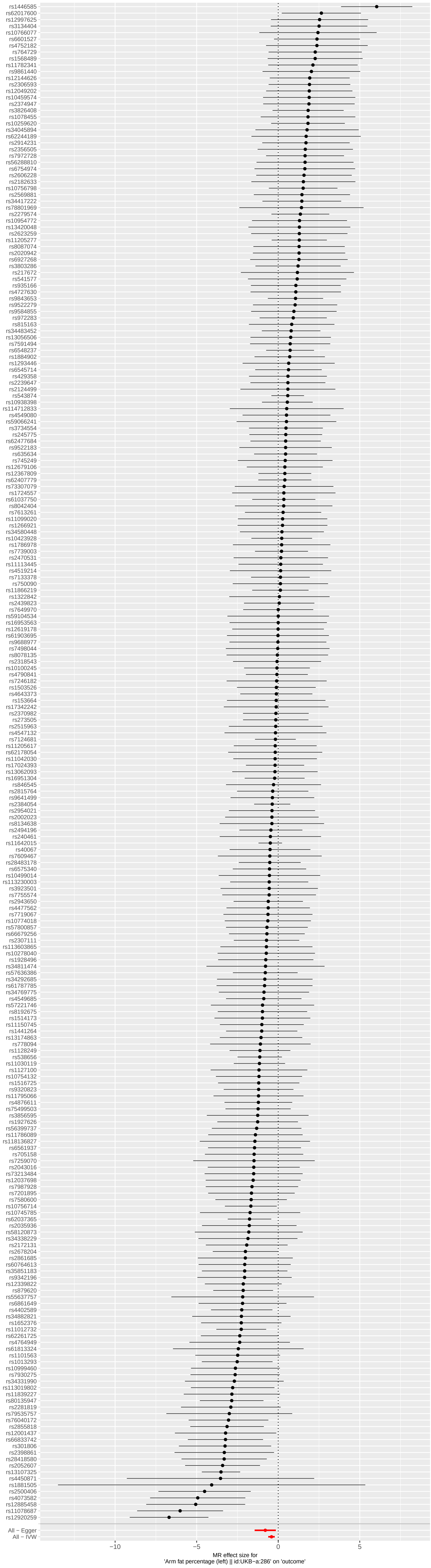
rs1446585
rs62017600
rs12997625
rs3134404
rs10766077
rs6601527
rs4752182
rs764729
rs1568489
rs11782341
rs9861440
rs12144626
rs2306593
rs12049202
rs10459574
rs2374947
rs3826408
rs1078455
rs10259620
rs34045894
rs62244189
rs2914231
rs2356505
rs7972728
rs56288810
rs6754974
rs2606228
rs2182633
rs10756798
rs2569881
rs34417222
rs78801969
rs2279574
rs10954772
rs13420048
rs2623259
rs11205277
rs8087074
rs2020942
rs6927268
rs3803286
rs217672
rs541577
rs935166
rs4727630
rs9843653
rs9522279
rs9584855
rs972283
rs815163
rs34483452
rs13056506
rs7591494
rs6548237
rs1884902
rs1293446
rs6545714
rs429358
rs2239647
rs2124499
rs543874
rs10938398
rs114712833
rs4549080
rs59066241
rs3734554
rs245775
rs62477684
rs9522183
rs635634
rs745249
rs12679106
rs12367809
rs62407779
rs73307079
rs1724557
rs61037750
rs8042404
rs7613261
rs11099020
rs1266921
rs34580448
rs10423928
rs1786978
rs7739003
rs2470531
rs11113445
rs4519214
rs7133378
rs750090
rs11866219
rs1322842
rs2439823
rs7649970
rs59104534
rs16953563
rs12619178
rs61903695
rs9688977
rs7498044
rs8078135
rs2318543
rs10100245
rs4790841
rs7246182
rs1503526
rs4643373
rs153664
rs17342242
rs2370982
rs273505
rs2515963
rs4547132
rs7124681
rs11205617
rs62178054
rs11042030
rs17024393
rs13062093
rs16951304
rs846545
rs2815764
rs9641499
rs2384054
rs2954021
rs2002023
rs8134638
rs2494196
rs240461
rs11642015
rs40067
rs7609467
rs28483178
rs6575340
rs10499014
rs113230003
rs3923501
rs7755574
rs2943650
rs4477562
rs7719067
rs10774018
rs57800857
rs66679256
rs2307111
rs113603865
rs10278040
rs1928496
rs34811474
rs57636386
rs34292685
rs61787785
rs34769775
rs4549685
rs57221746
rs8192675
rs1514173
rs11150745
rs1441264
rs13174863
rs778094
rs1128249
rs538656
rs11030119
rs1127100
rs10754132
rs1516725
rs9320823
rs11795066
rs4876611
rs75499503
rs3856595
rs1927626
rs56399737
rs11786089
rs118136827
rs6561937
rs705158
rs7259070
rs2043016
rs73213484
rs12037698
rs7987928
rs7201895
rs7580600
rs10756714
rs10745785
rs62037365
rs2035936
rs58120873
rs34338229
rs2172131
rs2678204
rs2861685
rs60764613
rs35851183
rs9342196
rs12339822
rs879620
rs55637757
rs6861649
rs4402589
rs34882821
rs1652376
rs11012732
rs62261725
rs4764949
rs61813324
rs11101563
rs1013293
rs10999460
rs7930275
rs34331990
rs113019802
rs11839227
rs80135947
rs2281819
rs79535757
rs76040172
rs2855818
rs12001437
rs66833742
rs301806
rs2398861
rs28418580
rs2052607
rs13107325
rs4450871
rs1881505
rs2500406
rs4073582
rs12885458
rs11078687
rs12920259
All − Egger
All − IVW
−10
−5
0
5
MR effect size for
'Arm fat percentage (left) || id:UKB−a:286' on 'outcome'

9

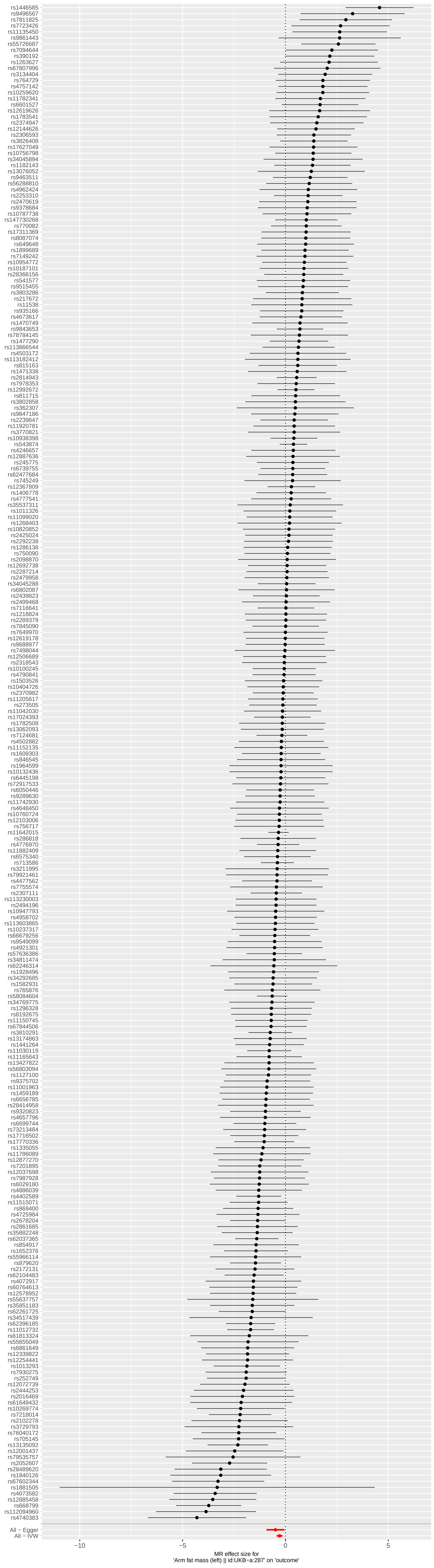

rs1446585
rs9496567
rs7811825
rs7723426
rs11135450
rs9861443
rs55726687
rs7094644
rs390192
rs1263627
rs67807996
rs3134404
rs764729
rs4757142
rs10259620
rs11782341
rs6601527
rs12619626
rs1783541
rs2374947
rs12144626
rs2306593
rs3826408
rs17627049
rs10756798
rs34045894
rs1182143
rs13076052
rs9463511
rs56288810
rs4962424
rs2253310
rs2470619
rs9378684
rs10787738
rs147730268
rs770082
rs17311369
rs8087074
rs649648
rs1899689
rs7149242
rs10954772
rs10187101
rs28366156
rs541577
rs9515455
rs3803286
rs217672
rs11538
rs935166
rs4673617
rs1470749
rs9843653
rs78784145
rs1477290
rs113866544
rs4503172
rs113182412
rs815163
rs1471338
rs2814943
rs7978353
rs12992672
rs811715
rs3802858
rs362307
rs9847186
rs2239647
rs11920781
rs3770821
rs10938398
rs543874
rs4246657
rs12887636
rs245775
rs6739755
rs62477684
rs745249
rs12367809
rs1406778
rs4777541
rs35537311
rs1011326
rs11099020
rs1268403
rs10820852
rs2425024
rs2292238
rs1286138
rs750090
rs2098870
rs12692738
rs2287214
rs2479958
rs34045288
rs6802087
rs2439823
rs2499468
rs7116641
rs1218824
rs2289379
rs7845090
rs7649970
rs12619178
rs9688977
rs7498044
rs12506689
rs2318543
rs10100245
rs4790841
rs1503526
rs10404726
rs2370982
rs11205617
rs273505
rs11042030
rs17024393
rs1782508
rs13062093
rs7124681
rs4502882
rs11152135
rs1609303
rs846545
rs1964599
rs10132436
rs6445198
rs72917533
rs6050446
rs9289630
rs11742930
rs4648450
rs10760724
rs12103006
rs756717
rs11642015
rs286818
rs4776970
rs11882409
rs6575340
rs713586
rs3211995
rs79921461
rs4477562
rs7755574
rs2307111
rs113230003
rs2494196
rs10947793
rs4958702
rs113603865
rs10237317
rs66679256
rs9549099
rs4921301
rs57636386
rs34811474
rs62246314
rs1928496
rs34292685
rs1582931
rs765876
rs58084604
rs34769775
rs1296328
rs8192675
rs11150745
rs67844506
rs3810291
rs13174863
rs1441264
rs11030119
rs11165643
rs13427822
rs56803094
rs1127100
rs9375702
rs11001963
rs1459189
rs6656785
rs28414958
rs9320823
rs4657796
rs6699744
rs73213484
rs17716502
rs17770336
rs1335055
rs11786089
rs12877270
rs7201895
rs12037698
rs7987928
rs6029180
rs4886039
rs4402589
rs11515071
rs869400
rs4725984
rs2678204
rs2861685
rs35882248
rs62037365
rs854917
rs1652376
rs55966114
rs879620
rs2172131
rs62104483
rs4072917
rs60764613
rs12578952
rs55637757
rs35851183
rs62261725
rs34517439
rs62396185
rs11012732
rs61813324
rs55655049
rs6861649
rs12339822
rs12254441
rs1013293
rs7930275
rs252749
rs12072739
rs2444253
rs2016469
rs61649432
rs10269774
rs7218014
rs2102278
rs3729793
rs76040172
rs705145
rs13135092
rs12001437
rs79535757
rs2052607
rs28489620
rs1840126
rs67602344
rs1881505
rs4073582
rs12885458
rs668799
rs112094960
rs4740383
All − Egger
All − IVW
−10
−5
0
5
MR effect size for
'Arm fat mass (left) || id:UKB−a:287' on 'outcome'

10

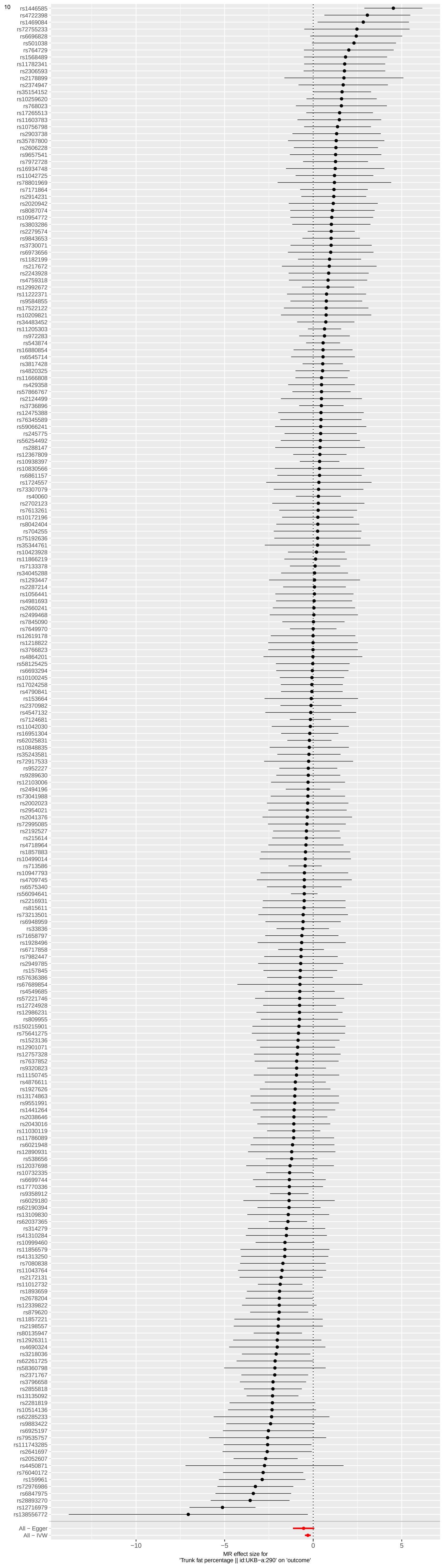
rs1446585
rs4722398
rs1469084
rs72755233
rs6696828
rs501038
rs764729
rs1568489
rs11782341
rs2306593
rs2178899
rs2374947
rs35154152
rs10259620
rs768023
rs17265513
rs11603783
rs10756798
rs2903738
rs35787800
rs2606228
rs9657541
rs7972728
rs16934748
rs11042725
rs78801969
rs7171864
rs2914231
rs2020942
rs8087074
rs10954772
rs3803286
rs2279574
rs9843653
rs3730071
rs6973656
rs1182199
rs217672
rs2243928
rs4759318
rs12992672
rs11222371
rs9584855
rs17522122
rs10209821
rs34483452
rs11205303
rs972283
rs543874
rs16880854
rs6545714
rs3817428
rs4820325
rs11666808
rs429358
rs57866767
rs2124499
rs3736896
rs12475388
rs76345589
rs59066241
rs245775
rs56254492
rs288147
rs12367809
rs10938397
rs10830566
rs6861157
rs1724557
rs73307079
rs40060
rs2702123
rs7613261
rs10172196
rs8042404
rs704255
rs75192636
rs35344761
rs10423928
rs11866219
rs7133378
rs34045288
rs1293447
rs2287214
rs1056441
rs4981693
rs2660241
rs2499468
rs7845090
rs7649970
rs12619178
rs1218822
rs3766823
rs4864201
rs58125425
rs6693294
rs10100245
rs17024258
rs4790841
rs153664
rs2370982
rs4547132
rs7124681
rs11042030
rs16951304
rs62025831
rs10848835
rs35243581
rs72917533
rs952227
rs9289630
rs12103006
rs2494196
rs73041988
rs2002023
rs2954021
rs2041376
rs72995085
rs2192527
rs215614
rs4718964
rs1857883
rs10499014
rs713586
rs10947793
rs4709745
rs6575340
rs56094641
rs2216931
rs815611
rs73213501
rs6948959
rs33836
rs71658797
rs1928496
rs6717858
rs7982447
rs2949785
rs157845
rs57636386
rs67689854
rs4549685
rs57221746
rs12724928
rs12986231
rs809955
rs150215901
rs75641275
rs1523136
rs12901071
rs12757328
rs7637852
rs9320823
rs11150745
rs4876611
rs1927626
rs13174863
rs9551991
rs1441264
rs2038646
rs2043016
rs11030119
rs11786089
rs6021948
rs12890931
rs538656
rs12037698
rs10732335
rs6699744
rs17770336
rs9358912
rs6029180
rs62190394
rs13109830
rs62037365
rs314279
rs41310284
rs10999460
rs11856579
rs41313250
rs7080838
rs11043764
rs2172131
rs11012732
rs1893659
rs2678204
rs12339822
rs879620
rs11857221
rs2198557
rs80135947
rs12926311
rs4690324
rs3218036
rs62261725
rs58360798
rs2371767
rs3796658
rs2855818
rs13135092
rs2281819
rs10514136
rs62285233
rs9883422
rs6925197
rs79535757
rs111743285
rs2641697
rs2052607
rs4450871
rs76040172
rs159961
rs72976986
rs6847975
rs28893270
rs12716979
rs138556772
All − Egger
All − IVW
−10
−5
0
5
MR effect size for
'Trunk fat percentage || id:UKB−a:290' on 'outcome'

11

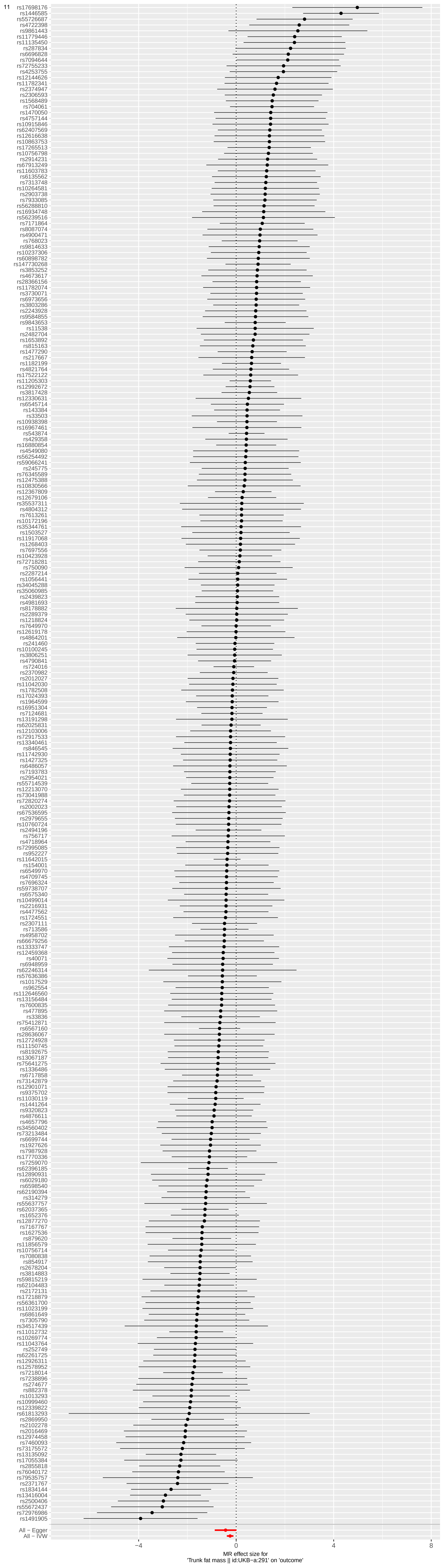
rs17698176
rs1446585
rs55726687
rs4722398
rs9861443
rs11779446
rs11135450
rs287834
rs6696828
rs7094644
rs72755233
rs4253755
rs12144626
rs11782341
rs2374947
rs2306593
rs1568489
rs704061
rs1470050
rs4757144
rs10915846
rs62407569
rs12616638
rs10863753
rs17265513
rs10756798
rs2914231
rs67913249
rs11603783
rs6135562
rs7313748
rs10264581
rs2903738
rs7933085
rs56288810
rs16934748
rs56239516
rs7171864
rs8087074
rs4900471
rs768023
rs9814633
rs10237306
rs60898782
rs147730268
rs3853252
rs4673617
rs28366156
rs11782074
rs3730071
rs6973656
rs3803286
rs2243928
rs9584855
rs9843653
rs11538
rs2482704
rs1653892
rs815163
rs1477290
rs217667
rs1182199
rs4821764
rs17522122
rs11205303
rs12992672
rs3817428
rs12330631
rs6545714
rs143384
rs33503
rs10938398
rs16967461
rs543874
rs429358
rs16880854
rs4549080
rs56254492
rs59066241
rs245775
rs76345589
rs12475388
rs10830566
rs12367809
rs12679106
rs35537311
rs4804312
rs7613261
rs10172196
rs35344761
rs1503527
rs11917068
rs1268403
rs7697556
rs10423928
rs72718281
rs750090
rs2287214
rs1056441
rs34045288
rs35060985
rs2439823
rs4981693
rs8178882
rs2289379
rs1218824
rs7649970
rs12619178
rs4864201
rs241460
rs10100245
rs3806251
rs4790841
rs724016
rs2370982
rs2012027
rs11042030
rs1782508
rs17024393
rs1964599
rs16951304
rs7124681
rs13191298
rs62025831
rs12103006
rs72917533
rs13340461
rs846545
rs11742930
rs1427325
rs6486057
rs7193783
rs2954021
rs55714539
rs12213070
rs73041988
rs72820274
rs2002023
rs67536595
rs2979655
rs10760724
rs2494196
rs756717
rs4718964
rs72995085
rs952227
rs11642015
rs154001
rs6549970
rs4709745
rs7696324
rs59738707
rs6575340
rs10499014
rs2216931
rs4477562
rs1724551
rs2307111
rs713586
rs4958702
rs66679256
rs13333747
rs12459368
rs40071
rs6948959
rs62246314
rs57636386
rs1017529
rs962554
rs112646560
rs13156484
rs7600835
rs477895
rs33836
rs75412871
rs6567160
rs28636067
rs12724928
rs11150745
rs8192675
rs13067187
rs75641275
rs1336486
rs6717858
rs73142879
rs12901071
rs9375702
rs11030119
rs1441264
rs9320823
rs4876611
rs4657796
rs34560402
rs73213484
rs6699744
rs1927626
rs7987928
rs17770336
rs7259070
rs62396185
rs12890931
rs6029180
rs6598540
rs62190394
rs314279
rs55637757
rs62037365
rs1652376
rs12877270
rs7167767
rs1627536
rs879620
rs11856579
rs10756714
rs7080838
rs854917
rs2678204
rs3814883
rs59815219
rs62104483
rs2172131
rs17218879
rs56361700
rs11023199
rs6861649
rs7305790
rs34517439
rs11012732
rs10269774
rs11043764
rs252749
rs62261725
rs12926311
rs12578952
rs7218014
rs7238896
rs274677
rs882378
rs1013293
rs10999460
rs12339822
rs61813293
rs2869950
rs2102278
rs2016469
rs12974458
rs7460093
rs73175572
rs13135092
rs17055384
rs2855818
rs76040172
rs79535757
rs2371767
rs1834144
rs13416004
rs2500406
rs55672437
rs72976986
rs1491905
All − Egger
All − IVW
−4
0
4
8
MR effect size for
'Trunk fat mass || id:UKB−a:291' on 'outcome'

12

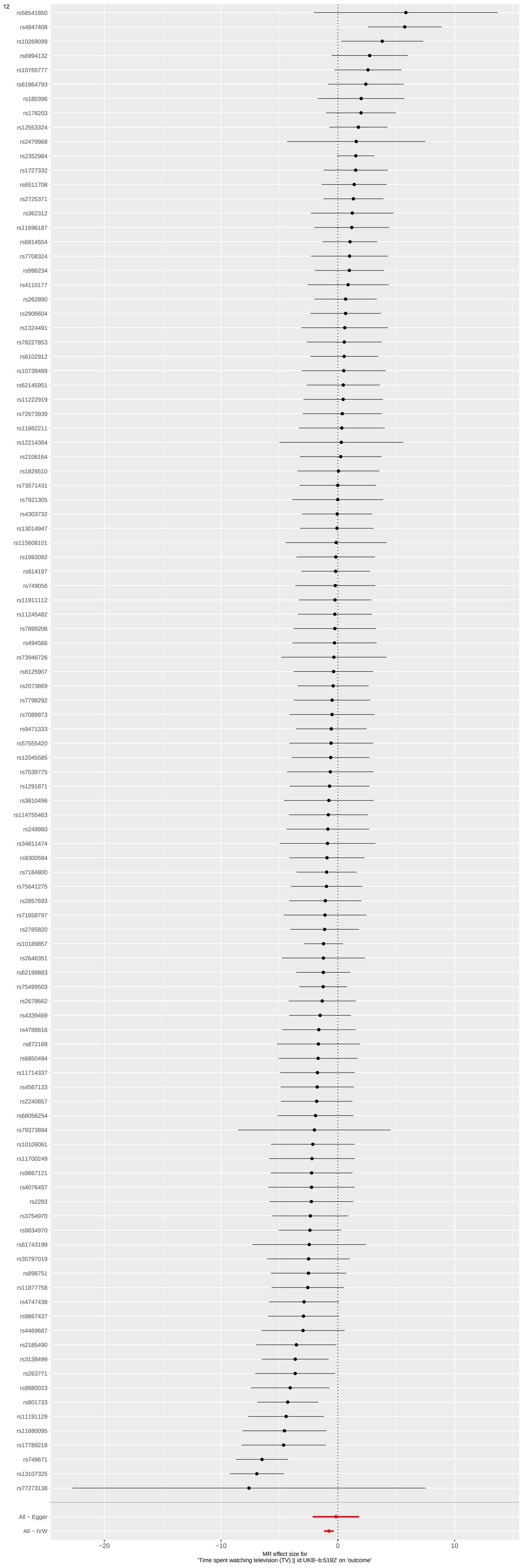
rs58541850
rs4847408
rs10269099
rs6994132
rs10765777
rs61864793
rs180396
rs178203
rs12553324
rs2479968
rs2352984
rs1727332
rs6511708
rs2725371
rs362312
rs11696187
rs6814554
rs7708324
rs996234
rs4110177
rs262890
rs2906604
rs1324491
rs78227853
rs6102912
rs10739499
rs62145951
rs11222919
rs72673939
rs11662211
rs12214364
rs2106164
rs1826510
rs73571431
rs7921305
rs4303732
rs13014947
rs115608101
rs1993092
rs814197
rs749056
rs11911112
rs11245482
rs7899206
rs494566
rs73946726
rs6125907
rs2073869
rs7798292
rs7089973
rs9471333
rs57555420
rs12045585
rs7539775
rs1291871
rs3810496
rs114755463
rs249960
rs34811474
rs9300594
rs7184800
rs75641275
rs2857693
rs71658797
rs2785820
rs10189857
rs2646351
rs62199883
rs75499503
rs2678662
rs4339469
rs4788616
rs872169
rs6850494
rs11714337
rs4567133
rs2240857
rs68056254
rs79373894
rs10109061
rs11700249
rs9867121
rs4076457
rs2283
rs3754970
rs9834970
rs61743199
rs35797019
rs898751
rs11877758
rs4747438
rs9867437
rs4469687
rs2185490
rs3138499
rs263771
rs9880023
rs801733
rs11191129
rs11680095
rs17789218
rs749671
rs13107325
rs77273138
All − Egger
All − IVW
−20
−10
0
10
MR effect size for
'Time spent watching television (TV) || id:UKB-b:5192' on 'outcome'

13

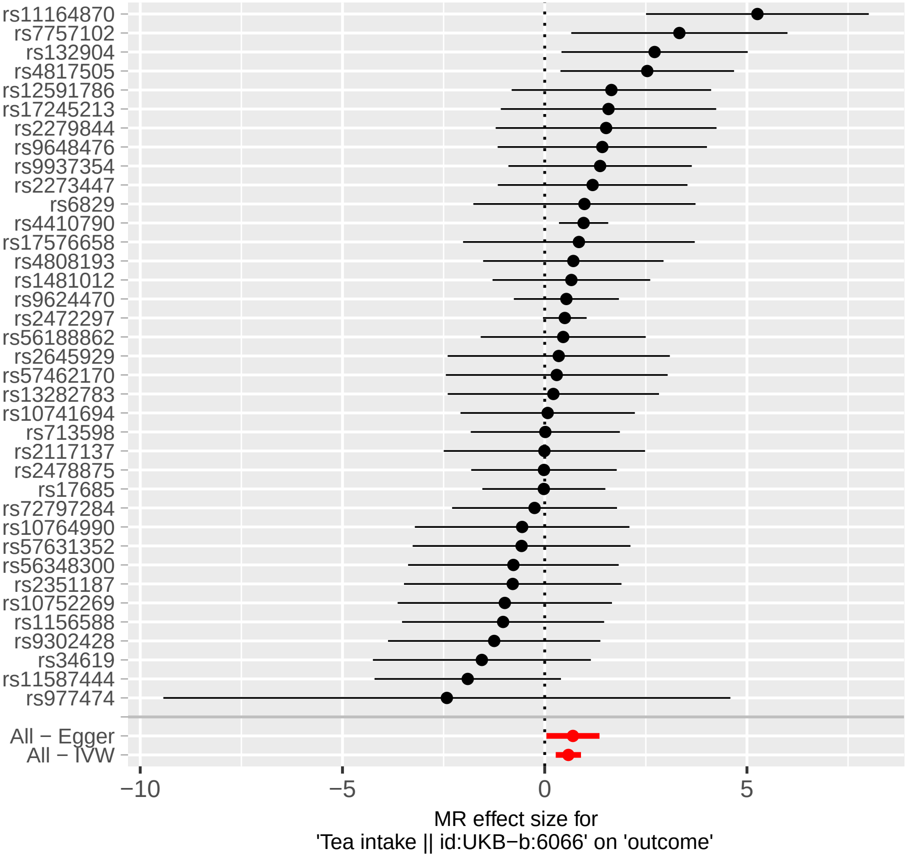
rs11164870
rs7757102
rs132904
rs4817505
rs12591786
rs17245213
rs2279844
rs9648476
rs9937354
rs2273447
rs6829
rs4410790
rs17576658
rs4808193
rs1481012
rs9624470
rs2472297
rs56188862
rs2645929
rs57462170
rs13282783
rs10741694
rs713598
rs2117137
rs2478875
rs17685
rs72797284
rs10764990
rs57631352
rs56348300
rs2351187
rs10752269
rs1156588
rs9302428
rs34619
rs11587444
rs977474
All − Egger
All − IVW
−10
−5
0
5
MR effect size for
'Tea intake || id:UKB−b:6066' on 'outcome'

### Figure S2. Funnel plots showing point estimates of the exposures of interest

(The trait appears at the bottom of each panel)

Each SNP's MR estimate is plotted against its minor-allele frequency corrected association with the exposure of interest. A minor allele frequency (MAF) correction proportional to the SNP-exposure standard error is used since a low-frequency allele is likely to be measured with low precision. Funnel plots can be used for visual inspection of symmetry, where any deviation can be suggestive of pleiotropy.

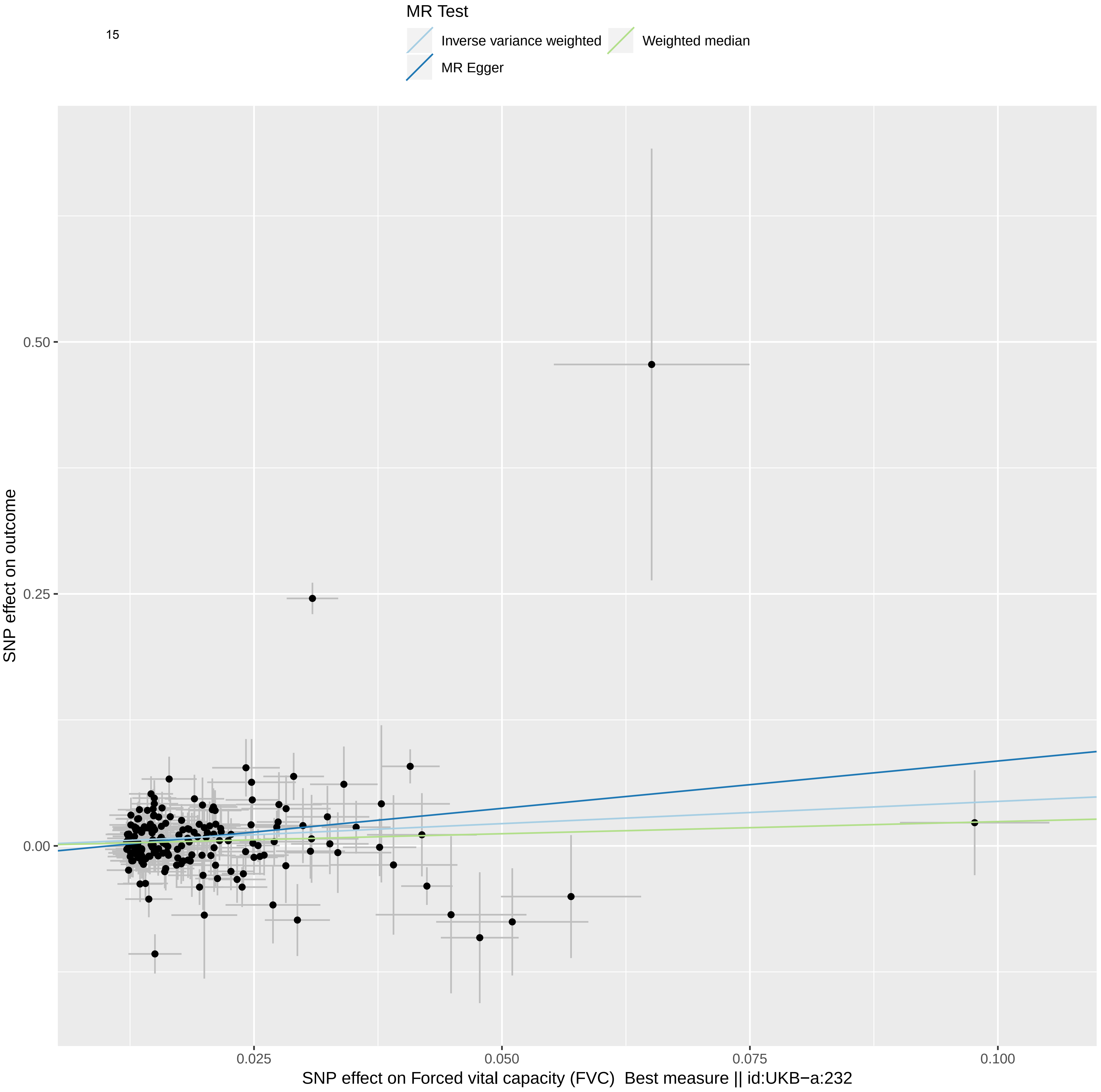

MR Test
Inverse variance weighted
Weighted median
MR Egger
SNP effect on outcome
0.50
0.25
0.00
0.025
0.050
0.075
0.100
SNP effect on Forced vital capacity (FVC) Best measure || id:UKB−a:232

16

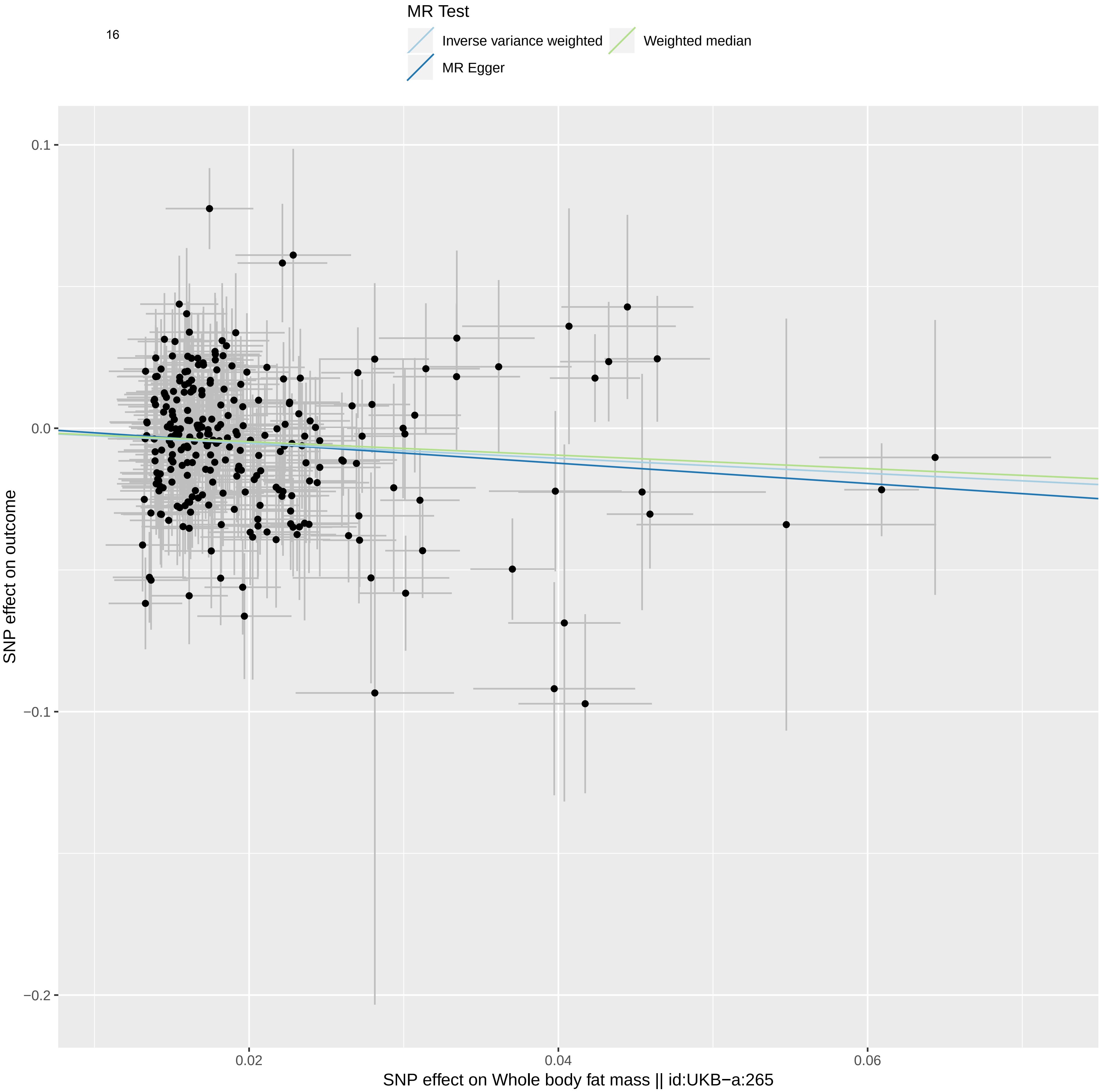

MR Test
Inverse variance weighted
Weighted median
MR Egger
0.1
0.0
−0.1
−0.2
0.02
0.04
0.06
SNP effect on outcome
SNP effect on Whole body fat mass || id:UKB−a:265

17

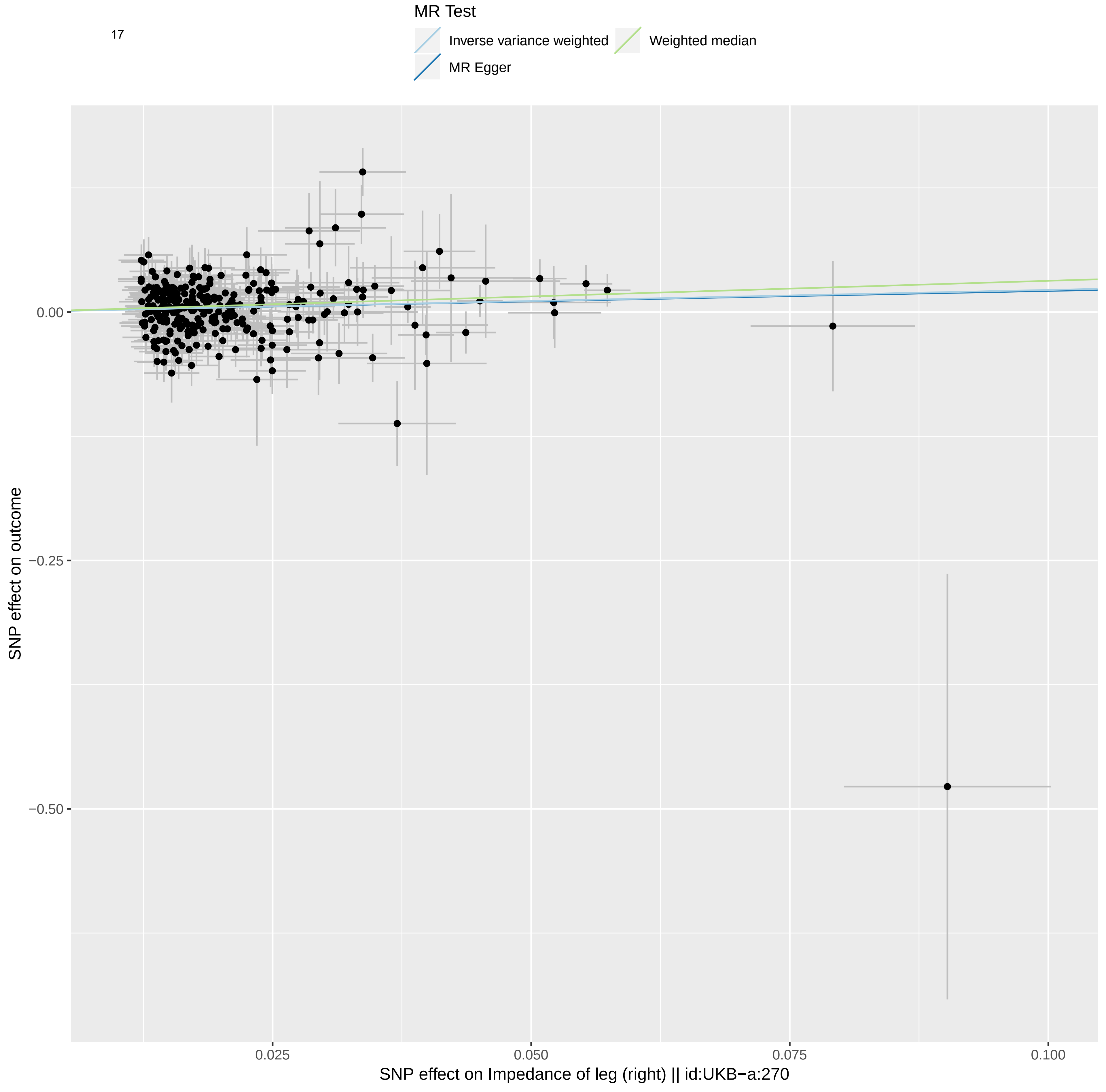
MR Test
Inverse variance weighted
Weighted median
MR Egger
0.00
−0.25
−0.50
SNP effect on outcome
0.025
0.050
0.075
0.100
SNP effect on Impedance of leg (right) || id:UKB−a:270

18

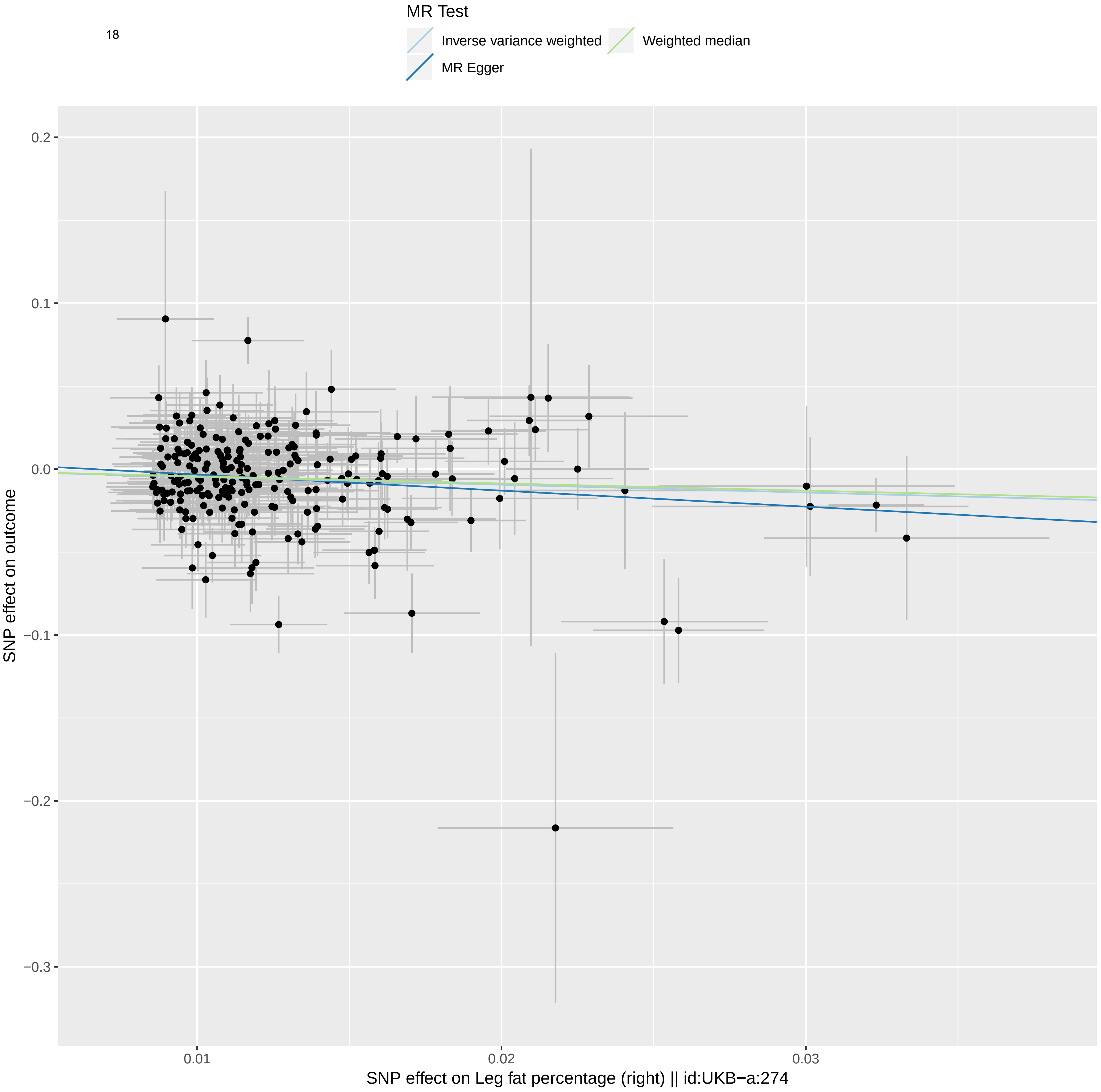

MR Test
Inverse variance weighted
Weighted median
MR Egger
0.2
0.1
0.0
−0.1
−0.2
−0.3
SNP effect on outcome
0.01
0.02
0.03
SNP effect on Leg fat percentage (right) || id:UKB−a:274

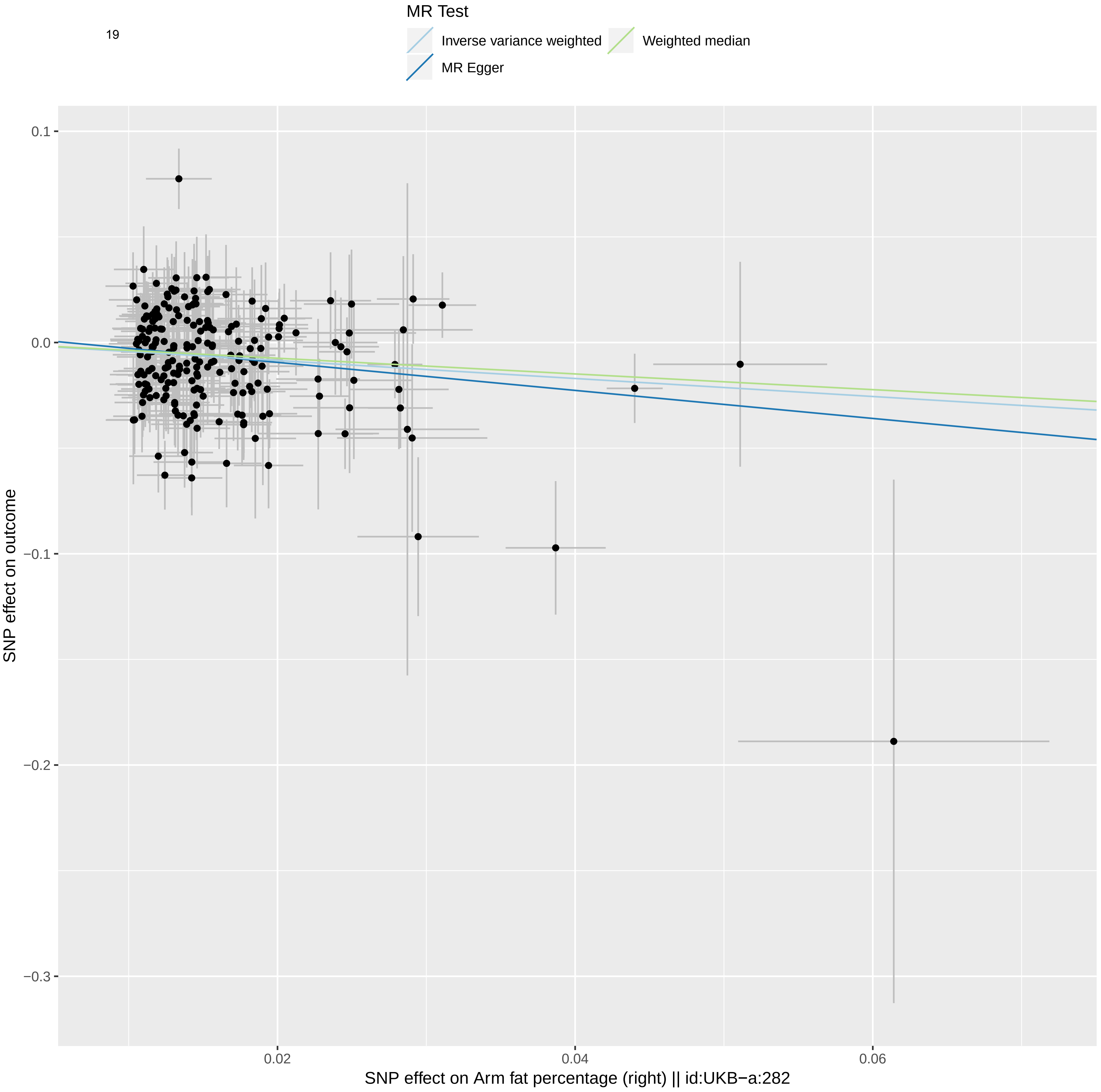
MR Test
Inverse variance weighted
Weighted median
MR Egger
0.1
0.0
−0.1
−0.2
−0.3
0.02
0.04
0.06
SNP effect on outcome
SNP effect on Arm fat percentage (right) || id:UKB−a:282

20

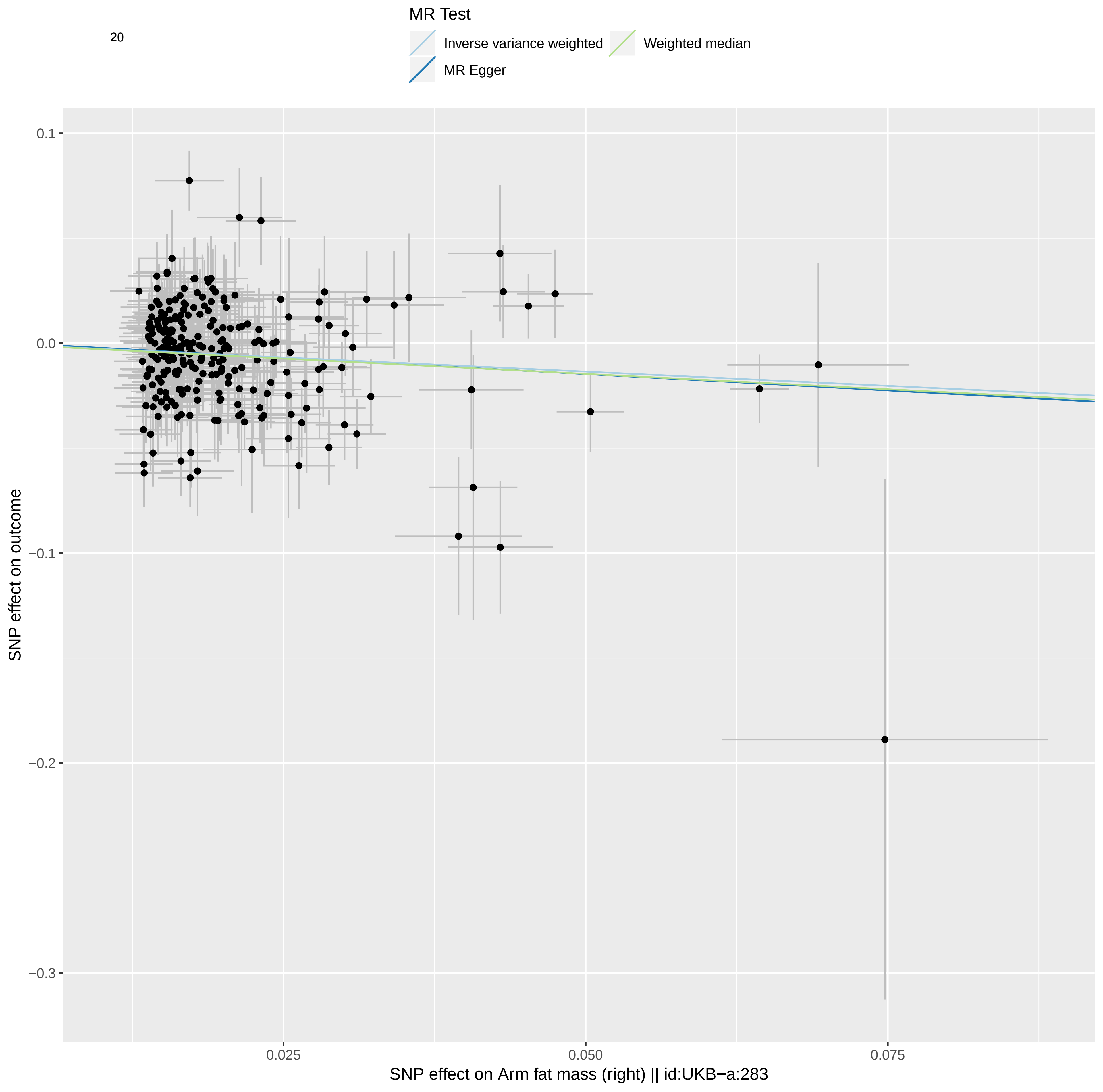

MR Test
Inverse variance weighted
Weighted median
MR Egger
0.1
0.0
−0.1
−0.2
−0.3
SNP effect on outcome
0.025
0.050
0.075
SNP effect on Arm fat mass (right) || id:UKB−a:283

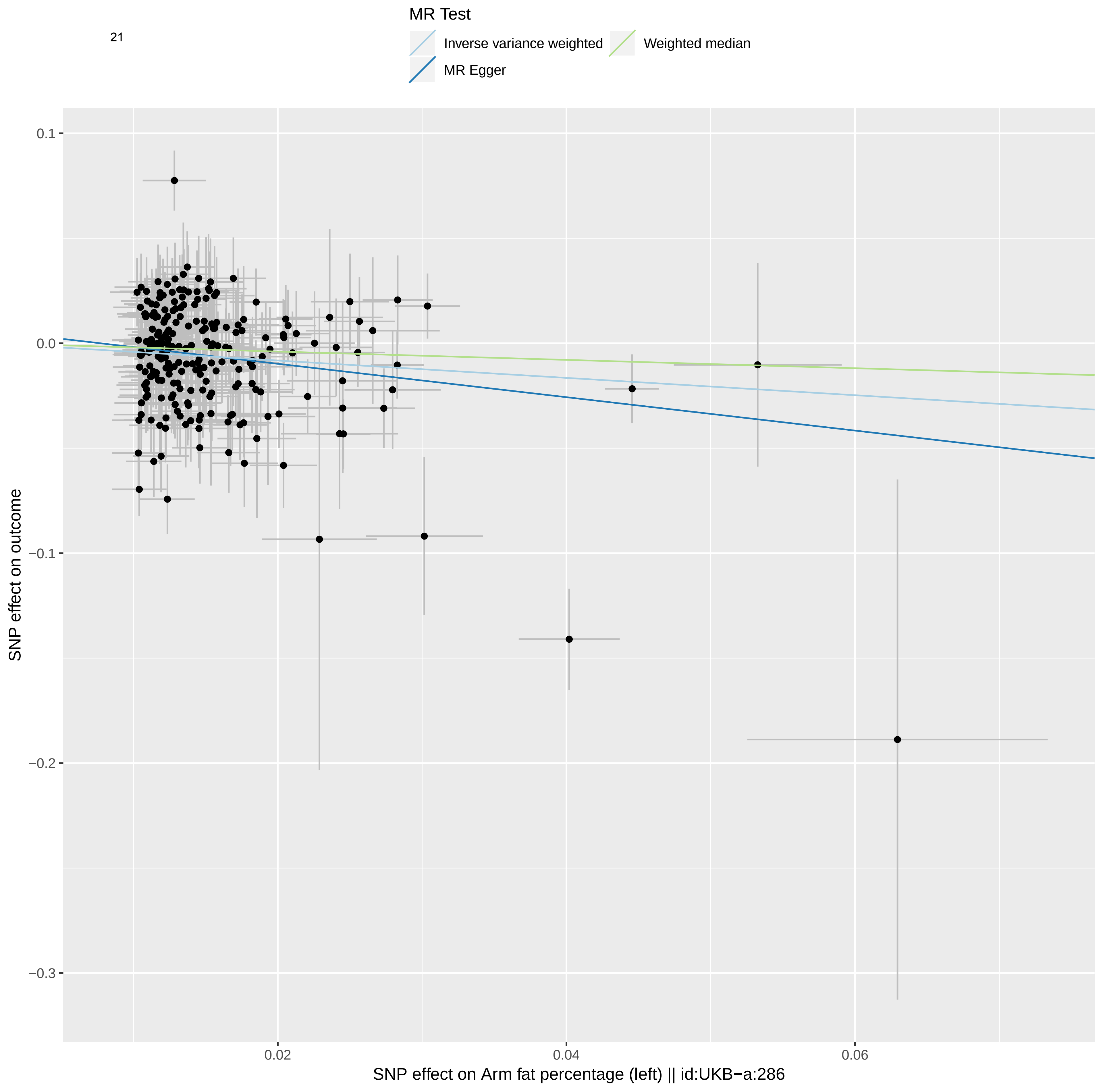

21
MR Test
Inverse variance weighted
Weighted median
MR Egger
0.1
0.0
−0.1
−0.2
−0.3
SNP effect on outcome
0.02
0.04
0.06
SNP effect on Arm fat percentage (left) || id:UKB−a:286

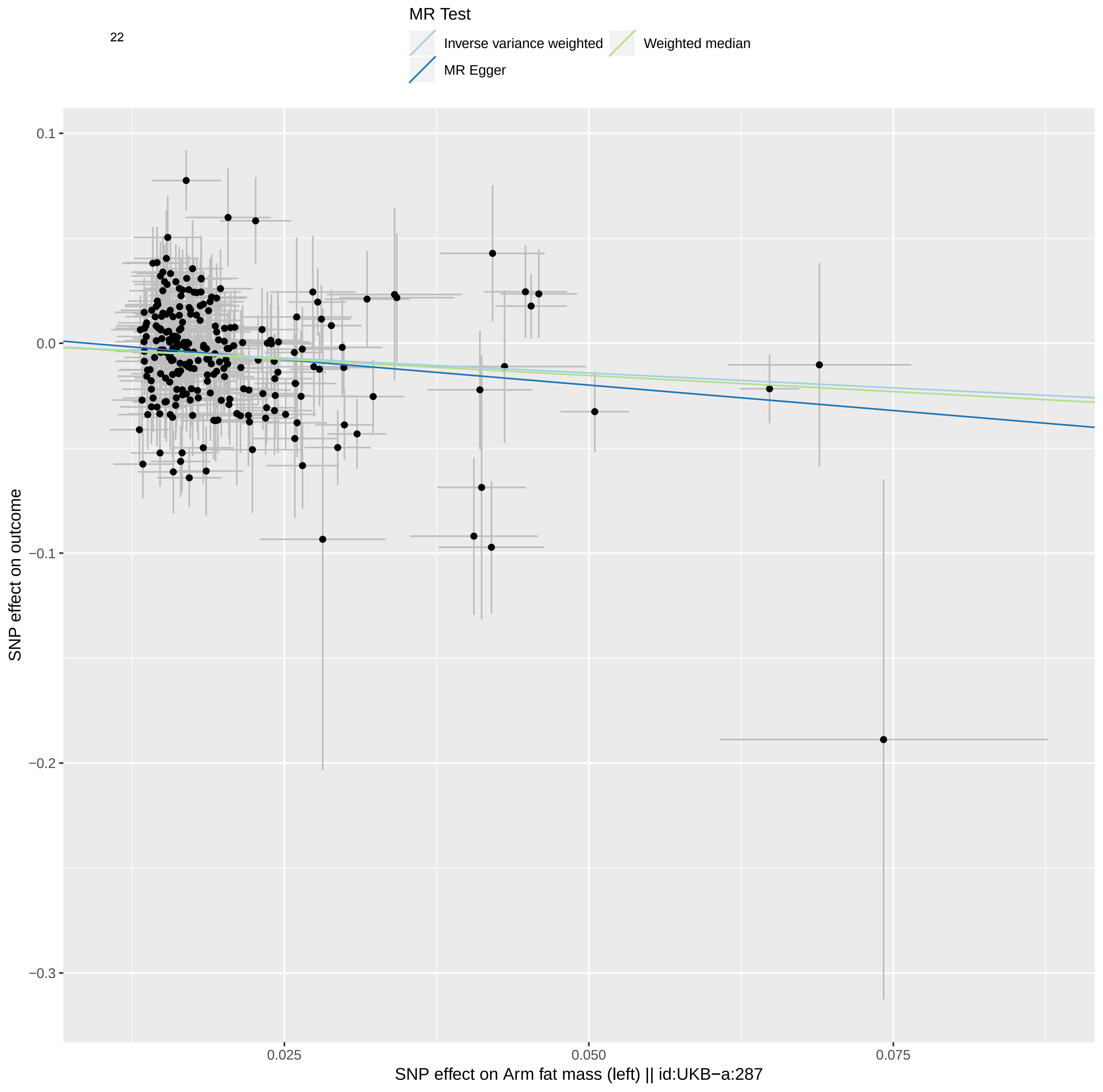
MR Test
Inverse variance weighted
Weighted median
MR Egger
0.1
0.0
−0.1
−0.2
−0.3
SNP effect on outcome
0.025
0.050
0.075
SNP effect on Arm fat mass (left) || id:UKB−a:287

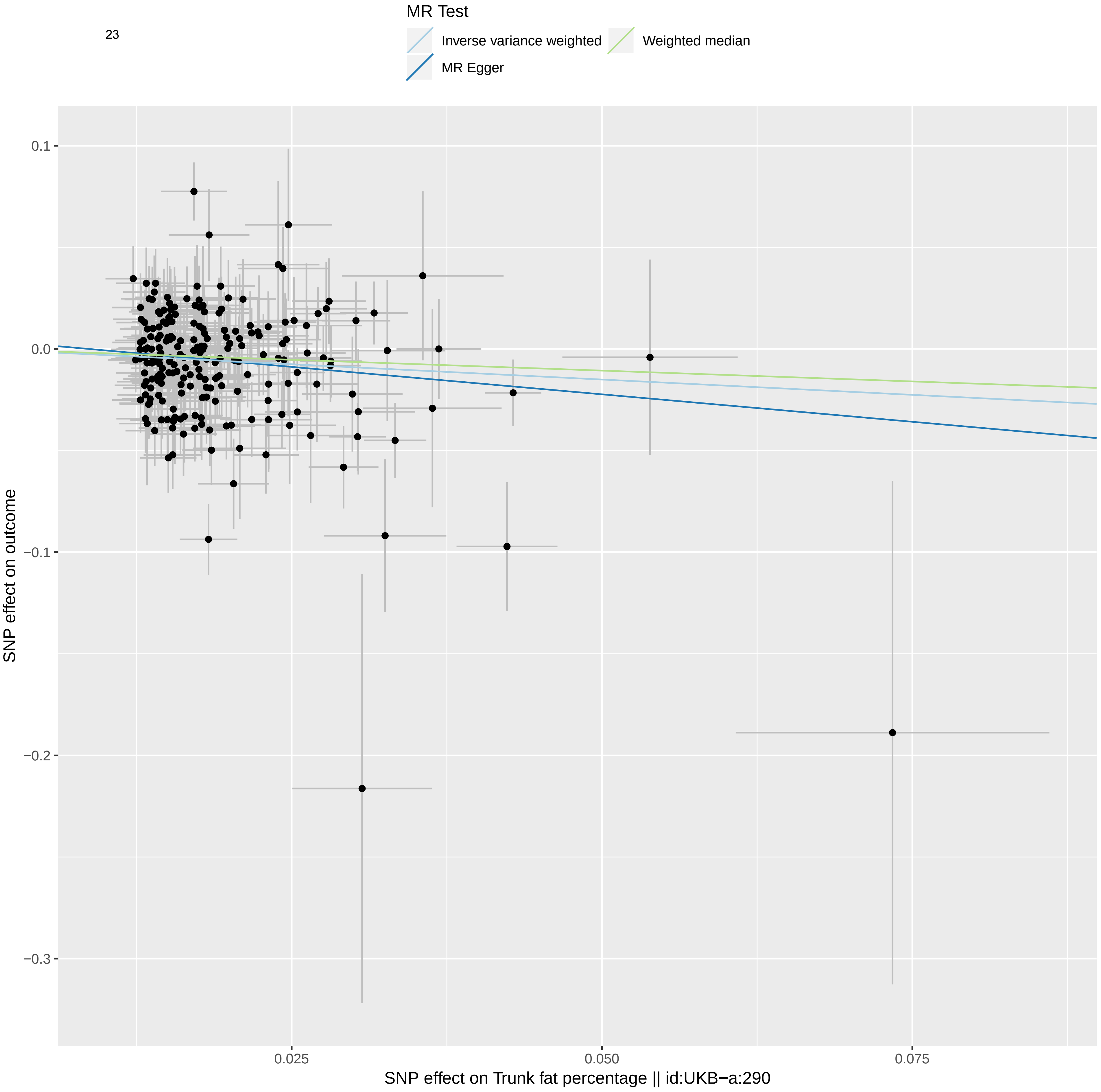
MR Test
Inverse variance weighted
Weighted median
MR Egger
0.1
0.0
−0.1
−0.2
−0.3
SNP effect on outcome
0.025
0.050
0.075
SNP effect on Trunk fat percentage || id:UKB−a:290

24

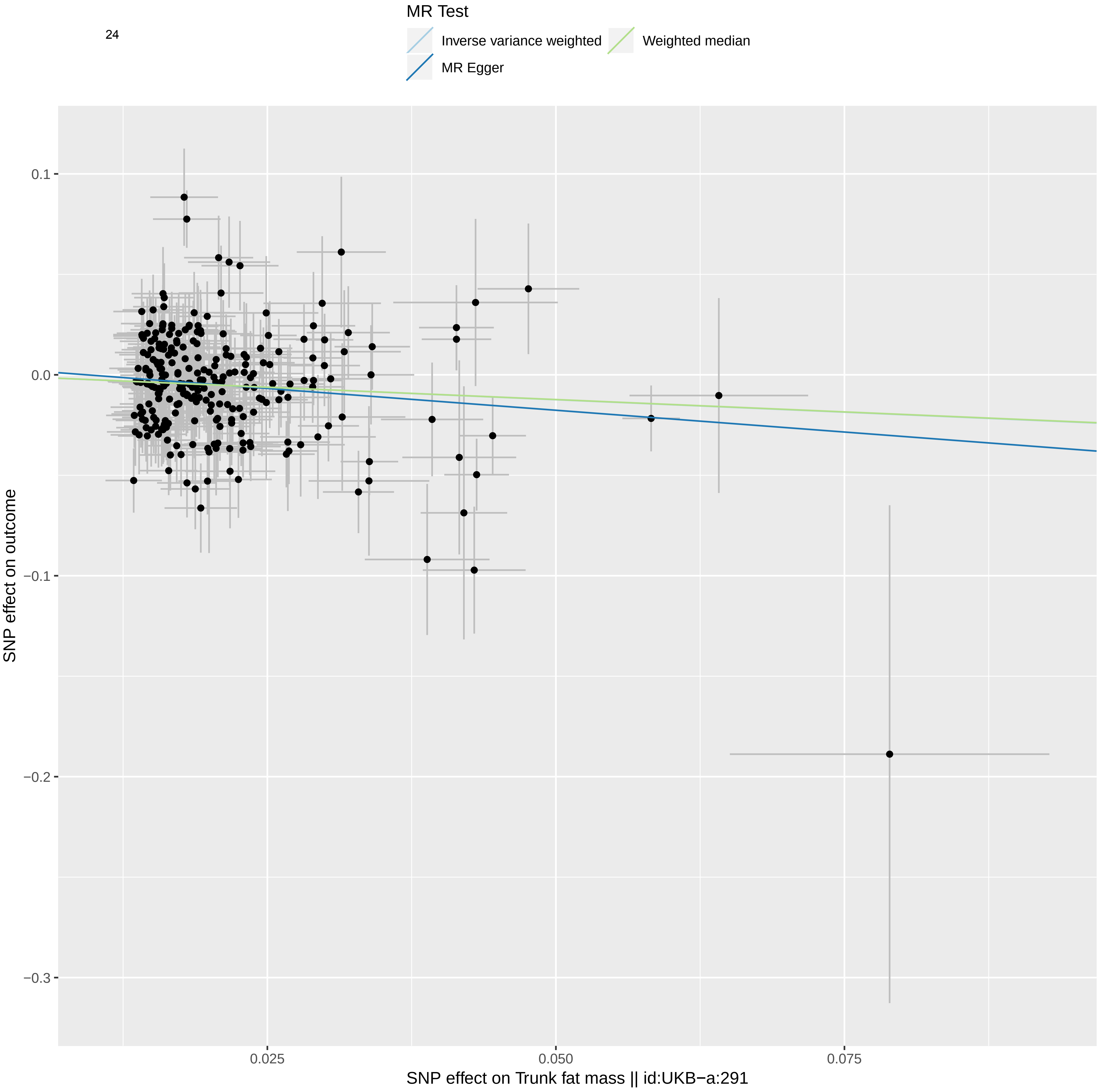
MR Test
Inverse variance weighted
Weighted median
MR Egger
0.1
0.0
−0.1
−0.2
−0.3
SNP effect on outcome
0.025
0.050
0.075
SNP effect on Trunk fat mass || id:UKB−a:291

25

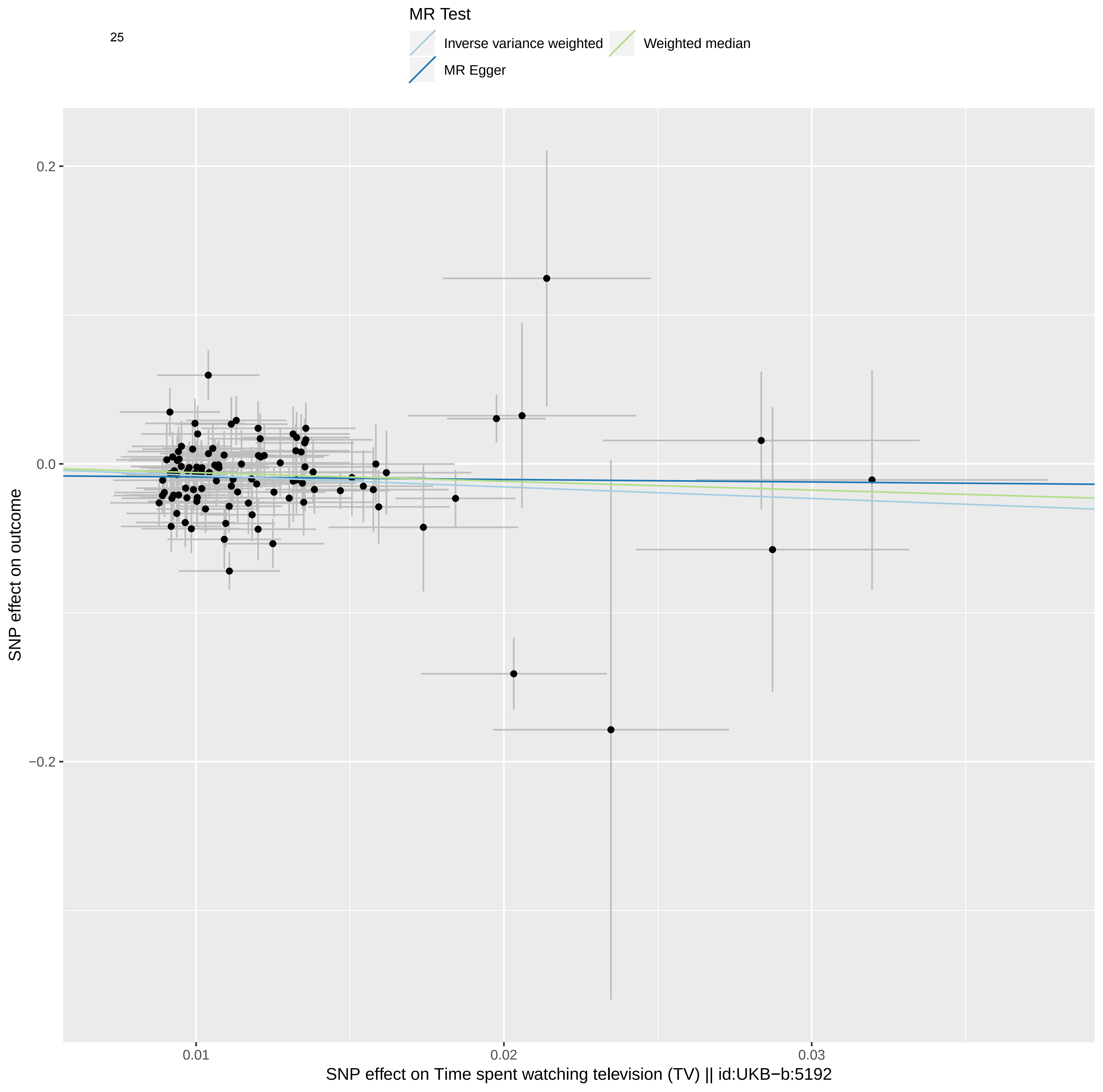
MR Test
Inverse variance weighted
Weighted median
MR Egger
0.2
0.0
−0.2
0.01
0.02
0.03
SNP effect on outcome
SNP effect on Time spent watching television (TV) || id:UKB−b:5192

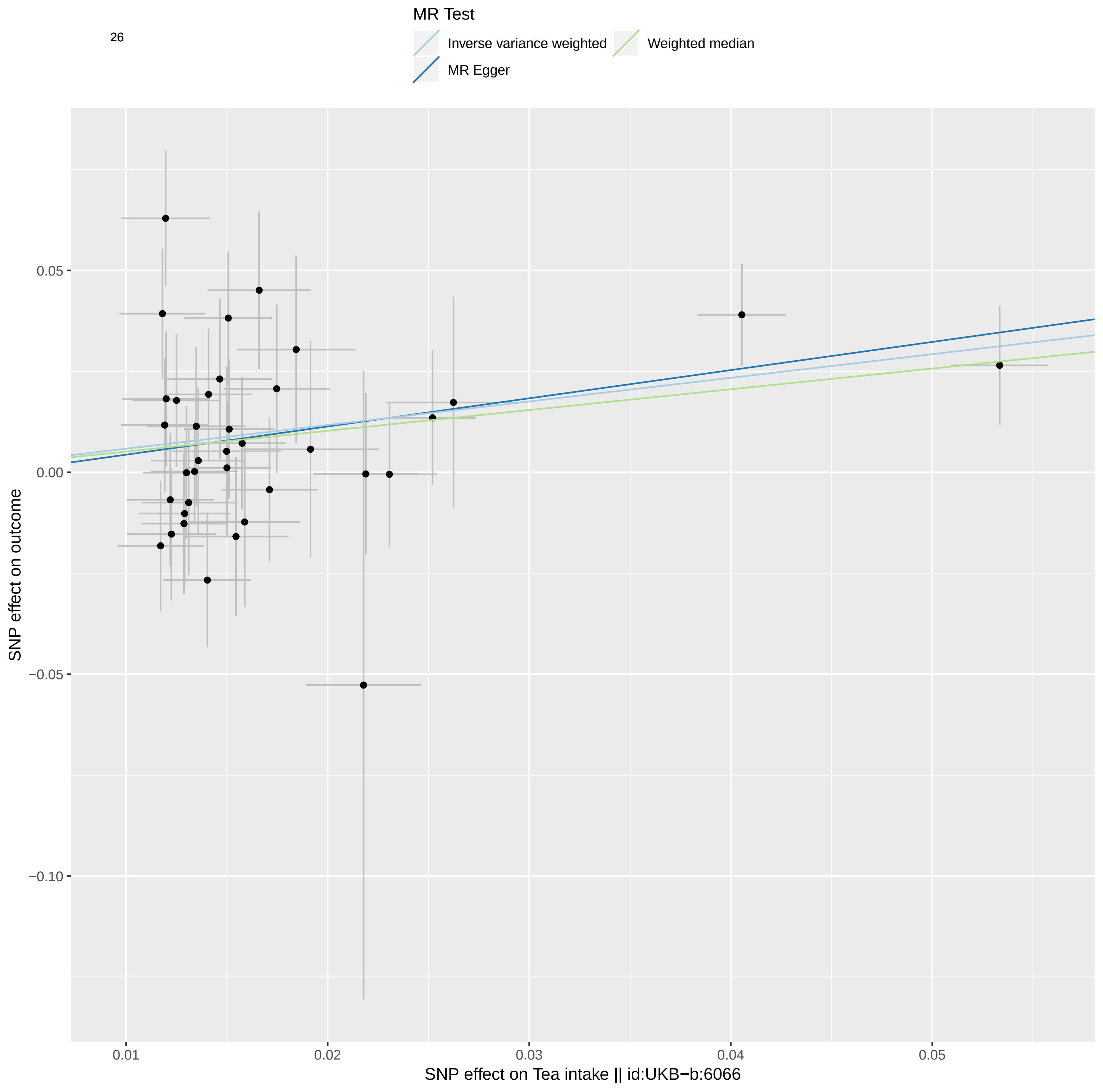
26
MR Test
Inverse variance weighted
Weighted median
MR Egger
SNP effect on outcome
0.05
0.00
−0.05
−0.10
0.01
0.02
0.03
0.04
0.05
SNP effect on Tea intake || id:UKB−b:6066

**Table S1.** Genetic correlations between fat related traits and impedance related traits

| Trait 1 | Trait 2 | rg | se | z | p | h2_obs | h2_obs_se | h2_int | h2_int_se | gcov_int | gcov_int_se |
|---|---|---|---|---|---|---|---|---|---|---|---|
| Arm fat percentage (left) \|\| id:UKB-a:286 | Impedance of arm (left) \|\| id:UKB-a:273 | -0.2947 | 0.0221 | -13.3194 | 1.78E-40 | 0.2301 | 0.008 | 1.0981 | 0.0205 | -0.184 | 0.0104 |

| Trait 1 | Trait 2 | rg | se | z | p | h2_obs | h2_obs_se | h2_int | h2_int_se | gcov_int | gcov_int_se |
|---|---|---|---|---|---|---|---|---|---|---|---|
| Arm fat percentage (right) \|\| id:UKB-a:282 | Impedance of arm (right) \|\| id:UKB-a:272 | -0.284 | 0.0222 | -12.8019 | 1.60E-37 | 0.2332 | 0.0079 | 1.085 | 0.021 | -0.1709 | 0.0102 |

| Trait 1 | Trait 2 | rg | se | z | p | h2_obs | h2_obs_se | h2_int | h2_int_se | gcov_int | gcov_int_se |
|---|---|---|---|---|---|---|---|---|---|---|---|
| Leg fat percentage (left) \|\| id:UKB-a:278 | Impedance of leg (left) \|\| id:UKB-a:271 | -0.2447 | 0.0239 | -10.2428 | 1.28E-24 | 0.2386 | 0.0091 | 1.0792 | 0.0217 | -0.1592 | 0.0106 |

| Trait 1 | Trait 2 | rg | se | z | p | h2_obs | h2_obs_se | h2_int | h2_int_se | gcov_int | gcov_int_se |
|---|---|---|---|---|---|---|---|---|---|---|---|
| Leg fat percentage (right) \|\| id:UKB-a:274 | Impedance of leg (right) \|\| id:UKB-a:270 | -0.2144 | 0.0243 | -8.8327 | 1.02E-18 | 0.2323 | 0.0089 | 1.0915 | 0.022 | -0.1254 | 0.0109 |

rg, genetic correlation; se, standard error; $h^2$ obs, narrow-sense heritability observed; $h^2$ int, narrow-sense heritability intercept, gcov_int, cross-trait intercept.

**Table S2.** Leave-one-out analyses for exposures of interest

**Impedance of leg (right) || id:UKB-a:270**

| SNP | b | se | p |
|---|---|---|---|
| rs10019888 | 0.225 | 0.066 | 6.55E-04 |
| rs1015032 | 0.223 | 0.066 | 7.29E-04 |
| rs10203320 | 0.230 | 0.065 | 4.32E-04 |
| rs10236214 | 0.219 | 0.066 | 9.27E-04 |
| rs1047891 | 0.222 | 0.066 | 7.98E-04 |
| rs1048932 | 0.225 | 0.066 | 6.59E-04 |
| rs10491967 | 0.223 | 0.066 | 7.31E-04 |
| rs1064213 | 0.217 | 0.066 | 1.01E-03 |
| rs10765992 | 0.223 | 0.066 | 7.09E-04 |
| rs10777454 | 0.221 | 0.066 | 8.09E-04 |
| rs1077750 | 0.224 | 0.066 | 7.00E-04 |
| rs10777859 | 0.228 | 0.066 | 5.15E-04 |
| rs10819047 | 0.221 | 0.066 | 8.20E-04 |
| rs10824307 | 0.222 | 0.066 | 7.75E-04 |
| rs10863578 | 0.221 | 0.066 | 8.15E-04 |
| rs1089946 | 0.223 | 0.066 | 7.15E-04 |
| rs10903340 | 0.232 | 0.066 | 4.12E-04 |
| rs10919482 | 0.221 | 0.066 | 7.98E-04 |
| rs10923724 | 0.222 | 0.066 | 7.76E-04 |
| rs10962552 | 0.225 | 0.066 | 6.30E-04 |
| rs11042717 | 0.220 | 0.066 | 8.38E-04 |
| rs11065979 | 0.225 | 0.066 | 6.52E-04 |
| rs11098676 | 0.218 | 0.066 | 8.99E-04 |
| rs11148421 | 0.221 | 0.066 | 8.05E-04 |
| rs11150745 | 0.221 | 0.066 | 7.83E-04 |
| rs11158820 | 0.222 | 0.066 | 7.51E-04 |
| rs11187838 | 0.224 | 0.066 | 7.36E-04 |
| rs11227313 | 0.224 | 0.066 | 7.05E-04 |
| rs112875651 | 0.220 | 0.066 | 8.50E-04 |
| rs11525873 | 0.222 | 0.066 | 7.81E-04 |
| rs11540358 | 0.226 | 0.066 | 6.07E-04 |
| rs11545482 | 0.222 | 0.066 | 7.41E-04 |
| rs115796963 | 0.217 | 0.066 | 9.76E-04 |
| rs11590254 | 0.223 | 0.066 | 7.33E-04 |
| rs115912456 | 0.223 | 0.066 | 7.38E-04 |
| rs11622473 | 0.223 | 0.066 | 7.28E-04 |
| rs11640504 | 0.232 | 0.066 | 4.05E-04 |
| rs11652817 | 0.222 | 0.066 | 7.48E-04 |
| rs11664242 | 0.226 | 0.066 | 6.16E-04 |
| rs11679731 | 0.224 | 0.066 | 7.01E-04 |

| | | | |
|---|---|---|---|
| rs11679846 | 0.221 | 0.066 | 8.15E-04 |
| rs116948941 | 0.226 | 0.066 | 5.60E-04 |
| rs11705686 | 0.222 | 0.066 | 7.75E-04 |
| rs117068593 | 0.222 | 0.066 | 8.01E-04 |
| rs11712037 | 0.222 | 0.066 | 7.74E-04 |
| rs11722554 | 0.224 | 0.066 | 7.08E-04 |
| rs11723923 | 0.226 | 0.066 | 6.24E-04 |
| rs11909249 | 0.223 | 0.066 | 7.35E-04 |
| rs1192923 | 0.217 | 0.066 | 9.68E-04 |
| rs12042908 | 0.219 | 0.066 | 8.95E-04 |
| rs12141801 | 0.228 | 0.066 | 5.05E-04 |
| rs12145281 | 0.225 | 0.066 | 6.32E-04 |
| rs12340775 | 0.221 | 0.066 | 7.95E-04 |
| rs12355141 | 0.220 | 0.066 | 8.46E-04 |
| rs1243473 | 0.222 | 0.066 | 7.73E-04 |
| rs12461902 | 0.218 | 0.066 | 9.08E-04 |
| rs12514357 | 0.222 | 0.066 | 7.44E-04 |
| rs1260326 | 0.228 | 0.066 | 5.64E-04 |
| rs12641981 | 0.227 | 0.066 | 5.52E-04 |
| rs12698917 | 0.220 | 0.066 | 8.32E-04 |
| rs12716975 | 0.214 | 0.066 | 1.15E-03 |
| rs12724928 | 0.225 | 0.066 | 6.36E-04 |
| rs12788682 | 0.217 | 0.066 | 9.30E-04 |
| rs12801226 | 0.224 | 0.066 | 6.63E-04 |
| rs12879626 | 0.225 | 0.066 | 6.28E-04 |
| rs12893681 | 0.227 | 0.066 | 5.64E-04 |
| rs12909300 | 0.225 | 0.066 | 6.35E-04 |
| rs12925700 | 0.221 | 0.066 | 7.99E-04 |
| rs13061214 | 0.222 | 0.066 | 7.59E-04 |
| rs13107325 | 0.200 | 0.064 | 1.72E-03 |
| rs13174863 | 0.221 | 0.066 | 7.94E-04 |
| rs13250787 | 0.229 | 0.066 | 4.66E-04 |
| rs1367228 | 0.223 | 0.066 | 7.15E-04 |
| rs1374657 | 0.225 | 0.066 | 6.60E-04 |
| rs1375566 | 0.224 | 0.066 | 6.88E-04 |
| rs140493137 | 0.225 | 0.066 | 6.31E-04 |
| rs147233090 | 0.222 | 0.066 | 7.64E-04 |
| rs1480474 | 0.223 | 0.066 | 7.16E-04 |
| rs1488373 | 0.224 | 0.066 | 6.73E-04 |
| rs150621 | 0.222 | 0.066 | 7.80E-04 |
| rs1517037 | 0.225 | 0.066 | 6.47E-04 |
| rs1593288 | 0.221 | 0.066 | 8.06E-04 |
| rs1607227 | 0.222 | 0.066 | 7.87E-04 |
| rs1614087 | 0.221 | 0.066 | 7.99E-04 |
| rs1635852 | 0.222 | 0.066 | 8.03E-04 |

| | | | |
|---|---|---|---|
| rs17110277 | 0.228 | 0.066 | 5.10E-04 |
| rs17115213 | 0.212 | 0.065 | 1.14E-03 |
| rs17159461 | 0.221 | 0.066 | 8.00E-04 |
| rs17265513 | 0.218 | 0.066 | 9.16E-04 |
| rs17373837 | 0.225 | 0.066 | 6.60E-04 |
| rs17419220 | 0.218 | 0.066 | 8.93E-04 |
| rs17466257 | 0.225 | 0.066 | 6.37E-04 |
| rs1748399 | 0.227 | 0.066 | 5.38E-04 |
| rs17776482 | 0.219 | 0.066 | 8.95E-04 |
| rs1831498 | 0.220 | 0.066 | 8.38E-04 |
| rs1880842 | 0.221 | 0.066 | 7.92E-04 |
| rs1951455 | 0.221 | 0.066 | 7.91E-04 |
| rs2114210 | 0.217 | 0.066 | 9.37E-04 |
| rs2118806 | 0.223 | 0.066 | 7.18E-04 |
| rs2203039 | 0.228 | 0.066 | 5.49E-04 |
| rs2206925 | 0.227 | 0.066 | 5.90E-04 |
| rs2208649 | 0.223 | 0.066 | 7.39E-04 |
| rs221308 | 0.223 | 0.066 | 7.46E-04 |
| rs2229840 | 0.228 | 0.066 | 5.51E-04 |
| rs2238435 | 0.217 | 0.066 | 9.85E-04 |
| rs2305756 | 0.220 | 0.066 | 8.38E-04 |
| rs2307111 | 0.221 | 0.066 | 7.94E-04 |
| rs2318761 | 0.225 | 0.066 | 6.37E-04 |
| rs2358483 | 0.221 | 0.066 | 8.21E-04 |
| rs2399213 | 0.225 | 0.066 | 6.57E-04 |
| rs2455561 | 0.225 | 0.066 | 6.39E-04 |
| rs245774 | 0.224 | 0.066 | 6.73E-04 |
| rs2517673 | 0.225 | 0.066 | 6.44E-04 |
| rs252924 | 0.223 | 0.066 | 7.44E-04 |
| rs252937 | 0.220 | 0.066 | 8.24E-04 |
| rs2530223 | 0.219 | 0.066 | 8.71E-04 |
| rs2611742 | 0.222 | 0.066 | 7.58E-04 |
| rs2621208 | 0.228 | 0.066 | 5.05E-04 |
| rs2624839 | 0.228 | 0.066 | 5.34E-04 |
| rs267803 | 0.223 | 0.066 | 7.28E-04 |
| rs2695165 | 0.220 | 0.066 | 8.45E-04 |
| rs2820295 | 0.217 | 0.066 | 9.47E-04 |
| rs2830585 | 0.222 | 0.066 | 7.91E-04 |
| rs28377268 | 0.224 | 0.066 | 6.87E-04 |
| rs2854152 | 0.227 | 0.066 | 5.78E-04 |
| rs2856448 | 0.228 | 0.066 | 5.33E-04 |
| rs2886373 | 0.220 | 0.066 | 8.19E-04 |
| rs2897968 | 0.222 | 0.066 | 7.57E-04 |
| rs2905424 | 0.221 | 0.066 | 8.10E-04 |
| rs29942 | 0.224 | 0.066 | 6.70E-04 |

| | | | |
|---|---|---|---|
| rs3118903 | 0.222 | 0.066 | 7.81E-04 |
| rs3170740 | 0.220 | 0.066 | 8.35E-04 |
| rs3205718 | 0.226 | 0.066 | 6.34E-04 |
| rs34517439 | 0.222 | 0.066 | 7.70E-04 |
| rs34571768 | 0.228 | 0.066 | 5.42E-04 |
| rs34811474 | 0.222 | 0.066 | 7.51E-04 |
| rs34838 | 0.219 | 0.066 | 8.24E-04 |
| rs35181861 | 0.224 | 0.066 | 6.62E-04 |
| rs351855 | 0.230 | 0.066 | 5.25E-04 |
| rs35320790 | 0.229 | 0.065 | 4.44E-04 |
| rs35597368 | 0.220 | 0.066 | 8.54E-04 |
| rs36083532 | 0.229 | 0.066 | 5.28E-04 |
| rs3732359 | 0.227 | 0.066 | 5.35E-04 |
| rs3751837 | 0.223 | 0.066 | 7.35E-04 |
| rs3760709 | 0.222 | 0.066 | 7.67E-04 |
| rs3764002 | 0.226 | 0.066 | 6.27E-04 |
| rs3792751 | 0.223 | 0.066 | 7.36E-04 |
| rs3798519 | 0.225 | 0.066 | 6.72E-04 |
| rs3810291 | 0.217 | 0.066 | 1.05E-03 |
| rs3815212 | 0.219 | 0.066 | 8.70E-04 |
| rs3816048 | 0.224 | 0.066 | 7.05E-04 |
| rs3818416 | 0.219 | 0.066 | 9.00E-04 |
| rs3897844 | 0.227 | 0.066 | 5.50E-04 |
| rs390444 | 0.218 | 0.066 | 8.95E-04 |
| rs3934135 | 0.222 | 0.067 | 8.62E-04 |
| rs395980 | 0.227 | 0.066 | 5.87E-04 |
| rs3966775 | 0.225 | 0.066 | 6.52E-04 |
| rs3971300 | 0.224 | 0.066 | 7.01E-04 |
| rs4277045 | 0.223 | 0.066 | 7.23E-04 |
| rs4350272 | 0.223 | 0.066 | 7.34E-04 |
| rs4373336 | 0.223 | 0.066 | 7.05E-04 |
| rs4472800 | 0.224 | 0.066 | 7.02E-04 |
| rs4515601 | 0.222 | 0.066 | 7.68E-04 |
| rs4672897 | 0.223 | 0.066 | 7.47E-04 |
| rs4719869 | 0.222 | 0.066 | 7.78E-04 |
| rs4721359 | 0.227 | 0.066 | 5.40E-04 |
| rs4776970 | 0.221 | 0.066 | 8.07E-04 |
| rs4819021 | 0.219 | 0.066 | 8.80E-04 |
| rs482190 | 0.225 | 0.066 | 6.46E-04 |
| rs4854909 | 0.222 | 0.066 | 7.57E-04 |
| rs4856876 | 0.226 | 0.066 | 5.82E-04 |
| rs4858696 | 0.223 | 0.066 | 7.46E-04 |
| rs490634 | 0.223 | 0.066 | 7.48E-04 |
| rs4965593 | 0.227 | 0.066 | 5.40E-04 |
| rs4976281 | 0.222 | 0.066 | 7.48E-04 |

| | | | |
|---|---|---|---|
| rs524344 | 0.223 | 0.066 | 7.40E-04 |
| rs539515 | 0.232 | 0.066 | 4.43E-04 |
| rs55695276 | 0.224 | 0.066 | 6.92E-04 |
| rs55983152 | 0.226 | 0.066 | 5.77E-04 |
| rs56088284 | 0.224 | 0.066 | 6.65E-04 |
| rs56172794 | 0.225 | 0.066 | 6.70E-04 |
| rs567056 | 0.221 | 0.066 | 8.16E-04 |
| rs572015 | 0.219 | 0.066 | 8.59E-04 |
| rs57307148 | 0.230 | 0.065 | 4.38E-04 |
| rs57636386 | 0.221 | 0.066 | 8.16E-04 |
| rs58939796 | 0.221 | 0.066 | 8.19E-04 |
| rs5995843 | 0.223 | 0.066 | 7.15E-04 |
| rs6014523 | 0.217 | 0.066 | 9.39E-04 |
| rs60609985 | 0.222 | 0.066 | 7.76E-04 |
| rs6063776 | 0.222 | 0.066 | 7.61E-04 |
| rs610476 | 0.221 | 0.066 | 8.09E-04 |
| rs6141551 | 0.226 | 0.066 | 6.01E-04 |
| rs61580113 | 0.224 | 0.066 | 6.87E-04 |
| rs61613873 | 0.222 | 0.066 | 7.49E-04 |
| rs61897977 | 0.223 | 0.066 | 7.00E-04 |
| rs61961465 | 0.222 | 0.066 | 7.46E-04 |
| rs62020775 | 0.221 | 0.066 | 8.00E-04 |
| rs62048402 | 0.217 | 0.067 | 1.24E-03 |
| rs62064607 | 0.223 | 0.066 | 6.97E-04 |
| rs62076439 | 0.229 | 0.066 | 5.06E-04 |
| rs62275493 | 0.223 | 0.066 | 7.14E-04 |
| rs62400317 | 0.222 | 0.066 | 7.71E-04 |
| rs62574545 | 0.223 | 0.066 | 7.26E-04 |
| rs62621812 | 0.225 | 0.066 | 6.69E-04 |
| rs6265 | 0.216 | 0.066 | 1.09E-03 |
| rs6445198 | 0.223 | 0.066 | 7.31E-04 |
| rs6452461 | 0.229 | 0.066 | 5.09E-04 |
| rs6480714 | 0.220 | 0.066 | 8.25E-04 |
| rs6485758 | 0.222 | 0.066 | 7.62E-04 |
| rs6494623 | 0.220 | 0.066 | 8.26E-04 |
| rs6503666 | 0.227 | 0.066 | 5.33E-04 |
| rs6548237 | 0.232 | 0.066 | 4.59E-04 |
| rs6596870 | 0.221 | 0.066 | 8.19E-04 |
| rs6607331 | 0.223 | 0.066 | 7.32E-04 |
| rs6657024 | 0.224 | 0.066 | 7.08E-04 |
| rs6664236 | 0.223 | 0.066 | 7.52E-04 |
| rs66768395 | 0.219 | 0.066 | 8.99E-04 |
| rs66824127 | 0.230 | 0.065 | 4.35E-04 |
| rs66848709 | 0.224 | 0.066 | 7.09E-04 |
| rs66922415 | 0.214 | 0.066 | 1.31E-03 |

| | | | |
|---|---|---|---|
| rs6711710 | 0.220 | 0.066 | 8.65E-04 |
| rs68049170 | 0.232 | 0.066 | 4.12E-04 |
| rs68083605 | 0.223 | 0.066 | 7.30E-04 |
| rs6822283 | 0.223 | 0.066 | 7.19E-04 |
| rs6852276 | 0.224 | 0.066 | 6.87E-04 |
| rs6860245 | 0.215 | 0.067 | 1.26E-03 |
| rs6869051 | 0.223 | 0.066 | 7.31E-04 |
| rs6871876 | 0.223 | 0.066 | 7.32E-04 |
| rs6916983 | 0.222 | 0.066 | 7.59E-04 |
| rs6950388 | 0.222 | 0.066 | 7.56E-04 |
| rs6962980 | 0.225 | 0.066 | 6.43E-04 |
| rs700910 | 0.227 | 0.066 | 5.72E-04 |
| rs7021479 | 0.224 | 0.066 | 6.62E-04 |
| rs705150 | 0.220 | 0.066 | 8.54E-04 |
| rs7102273 | 0.230 | 0.066 | 4.79E-04 |
| rs7126859 | 0.227 | 0.066 | 5.60E-04 |
| rs713586 | 0.220 | 0.066 | 8.47E-04 |
| rs7173751 | 0.218 | 0.066 | 9.10E-04 |
| rs720856 | 0.226 | 0.066 | 5.98E-04 |
| rs72721979 | 0.220 | 0.066 | 8.28E-04 |
| rs72755233 | 0.218 | 0.066 | 9.20E-04 |
| rs72796402 | 0.232 | 0.065 | 4.04E-04 |
| rs72853854 | 0.220 | 0.066 | 8.25E-04 |
| rs72885917 | 0.226 | 0.066 | 6.15E-04 |
| rs72916126 | 0.223 | 0.066 | 7.13E-04 |
| rs72948836 | 0.223 | 0.066 | 7.36E-04 |
| rs73052033 | 0.219 | 0.066 | 9.51E-04 |
| rs73052131 | 0.222 | 0.066 | 7.62E-04 |
| rs73230339 | 0.218 | 0.066 | 8.97E-04 |
| rs73245728 | 0.221 | 0.066 | 7.92E-04 |
| rs7328187 | 0.224 | 0.066 | 6.79E-04 |
| rs736839 | 0.223 | 0.066 | 7.09E-04 |
| rs738084 | 0.220 | 0.066 | 8.24E-04 |
| rs74501871 | 0.224 | 0.066 | 7.00E-04 |
| rs748457 | 0.219 | 0.066 | 8.77E-04 |
| rs752015 | 0.216 | 0.065 | 9.23E-04 |
| rs75336464 | 0.220 | 0.066 | 8.21E-04 |
| rs7561002 | 0.225 | 0.066 | 6.28E-04 |
| rs75625682 | 0.221 | 0.066 | 8.14E-04 |
| rs7592852 | 0.223 | 0.066 | 7.25E-04 |
| rs7596038 | 0.225 | 0.066 | 6.48E-04 |
| rs7649884 | 0.220 | 0.066 | 8.43E-04 |
| rs7701134 | 0.229 | 0.066 | 5.02E-04 |
| rs7732234 | 0.224 | 0.066 | 6.80E-04 |
| rs77364196 | 0.218 | 0.066 | 8.80E-04 |

| | | | |
|---|---|---|---|
| rs77974561 | 0.225 | 0.066 | 6.27E-04 |
| rs78135519 | 0.229 | 0.066 | 4.82E-04 |
| rs7821392 | 0.218 | 0.066 | 8.99E-04 |
| rs78378222 | 0.226 | 0.066 | 5.67E-04 |
| rs7839319 | 0.222 | 0.066 | 7.56E-04 |
| rs7861842 | 0.220 | 0.066 | 8.62E-04 |
| rs7868984 | 0.220 | 0.066 | 8.17E-04 |
| rs7902581 | 0.221 | 0.066 | 7.90E-04 |
| rs79616223 | 0.230 | 0.065 | 4.27E-04 |
| rs7978659 | 0.226 | 0.066 | 5.90E-04 |
| rs7994814 | 0.225 | 0.066 | 6.44E-04 |
| rs8051363 | 0.224 | 0.066 | 6.77E-04 |
| rs8071080 | 0.220 | 0.066 | 8.34E-04 |
| rs8104651 | 0.222 | 0.066 | 7.53E-04 |
| rs8176671 | 0.228 | 0.066 | 5.41E-04 |
| rs823096 | 0.216 | 0.065 | 9.40E-04 |
| rs905938 | 0.221 | 0.066 | 8.67E-04 |
| rs9288528 | 0.224 | 0.066 | 6.93E-04 |
| rs9289413 | 0.223 | 0.066 | 7.21E-04 |
| rs9375702 | 0.221 | 0.066 | 7.91E-04 |
| rs9391253 | 0.226 | 0.066 | 5.92E-04 |
| rs9469780 | 0.218 | 0.066 | 9.46E-04 |
| rs9496567 | 0.229 | 0.065 | 4.69E-04 |
| rs958225 | 0.226 | 0.066 | 5.88E-04 |
| rs960923 | 0.223 | 0.066 | 7.19E-04 |
| rs9638713 | 0.222 | 0.066 | 7.75E-04 |
| rs9647379 | 0.226 | 0.066 | 5.96E-04 |
| rs982004 | 0.227 | 0.066 | 5.62E-04 |
| rs9828976 | 0.223 | 0.066 | 7.06E-04 |
| rs9896535 | 0.217 | 0.066 | 9.93E-04 |
| rs9922288 | 0.222 | 0.066 | 7.80E-04 |
| rs9983616 | 0.221 | 0.066 | 8.15E-04 |
| All | 0.223 | 0.066 | 7.05E-04 |

**Trunk fat percentage || id:UKB-a:290**

| SNP | b | se | p |
|---|---|---|---|
| rs10100245 | -0.302 | 0.085 | 4.00E-04 |
| rs10172196 | -0.303 | 0.085 | 3.76E-04 |
| rs10209821 | -0.304 | 0.085 | 3.64E-04 |
| rs10259620 | -0.309 | 0.085 | 2.71E-04 |
| rs10423928 | -0.304 | 0.085 | 3.67E-04 |
| rs10499014 | -0.301 | 0.085 | 4.22E-04 |
| rs10514136 | -0.295 | 0.085 | 5.06E-04 |
| rs1056441 | -0.302 | 0.085 | 3.93E-04 |
| rs10732335 | -0.291 | 0.085 | 6.43E-04 |
| rs10756798 | -0.309 | 0.085 | 2.77E-04 |
| rs10830566 | -0.303 | 0.085 | 3.82E-04 |
| rs10848835 | -0.301 | 0.085 | 4.12E-04 |
| rs10938397 | -0.310 | 0.086 | 2.90E-04 |
| rs10947793 | -0.300 | 0.085 | 4.25E-04 |
| rs10954772 | -0.305 | 0.085 | 3.37E-04 |
| rs10999460 | -0.293 | 0.085 | 5.78E-04 |
| rs11012732 | -0.284 | 0.085 | 8.17E-04 |
| rs11030119 | -0.295 | 0.085 | 5.47E-04 |
| rs11042030 | -0.302 | 0.085 | 4.09E-04 |
| rs11042725 | -0.306 | 0.085 | 3.18E-04 |
| rs11043764 | -0.297 | 0.085 | 4.83E-04 |
| rs11150745 | -0.299 | 0.085 | 4.49E-04 |
| rs111743285 | -0.295 | 0.085 | 5.16E-04 |
| rs11205303 | -0.319 | 0.086 | 1.87E-04 |
| rs11222371 | -0.305 | 0.085 | 3.50E-04 |
| rs11603783 | -0.306 | 0.085 | 3.14E-04 |
| rs11666808 | -0.307 | 0.085 | 3.22E-04 |
| rs11782341 | -0.308 | 0.085 | 2.90E-04 |
| rs11786089 | -0.298 | 0.085 | 4.63E-04 |
| rs1182199 | -0.308 | 0.085 | 3.04E-04 |
| rs11856579 | -0.298 | 0.085 | 4.75E-04 |
| rs11857221 | -0.297 | 0.085 | 4.91E-04 |
| rs11866219 | -0.303 | 0.085 | 3.79E-04 |
| rs12037698 | -0.298 | 0.085 | 4.64E-04 |
| rs12103006 | -0.301 | 0.085 | 4.18E-04 |
| rs1218822 | -0.302 | 0.085 | 4.01E-04 |
| rs12339822 | -0.295 | 0.085 | 5.26E-04 |
| rs12367809 | -0.306 | 0.085 | 3.37E-04 |
| rs12475388 | -0.303 | 0.085 | 3.75E-04 |
| rs12619178 | -0.302 | 0.085 | 4.00E-04 |
| rs12716979 | -0.277 | 0.082 | 7.36E-04 |
| rs12724928 | -0.299 | 0.085 | 4.53E-04 |

| | | | |
|---|---|---|---|
| rs12757328 | -0.299 | 0.085 | 4.46E-04 |
| rs12890931 | -0.298 | 0.085 | 4.60E-04 |
| rs12901071 | -0.299 | 0.085 | 4.58E-04 |
| rs12926311 | -0.296 | 0.085 | 4.94E-04 |
| rs1293447 | -0.302 | 0.085 | 3.97E-04 |
| rs12986231 | -0.300 | 0.085 | 4.40E-04 |
| rs12992672 | -0.310 | 0.085 | 2.76E-04 |
| rs13109830 | -0.298 | 0.085 | 4.78E-04 |
| rs13135092 | -0.285 | 0.085 | 7.52E-04 |
| rs13174863 | -0.299 | 0.085 | 4.51E-04 |
| rs138556772 | -0.299 | 0.085 | 4.25E-04 |
| rs1441264 | -0.299 | 0.085 | 4.60E-04 |
| rs1446585 | -0.332 | 0.081 | 4.50E-05 |
| rs1469084 | -0.309 | 0.085 | 2.59E-04 |
| rs150215901 | -0.300 | 0.085 | 4.37E-04 |
| rs1523136 | -0.299 | 0.085 | 4.47E-04 |
| rs153664 | -0.301 | 0.085 | 4.06E-04 |
| rs1568489 | -0.308 | 0.085 | 2.92E-04 |
| rs157845 | -0.299 | 0.085 | 4.49E-04 |
| rs159961 | -0.294 | 0.085 | 5.30E-04 |
| rs16880854 | -0.306 | 0.085 | 3.30E-04 |
| rs16934748 | -0.304 | 0.085 | 3.48E-04 |
| rs16951304 | -0.302 | 0.085 | 4.13E-04 |
| rs17024258 | -0.302 | 0.085 | 4.00E-04 |
| rs1724557 | -0.302 | 0.085 | 3.91E-04 |
| rs17265513 | -0.310 | 0.085 | 2.67E-04 |
| rs17522122 | -0.304 | 0.085 | 3.58E-04 |
| rs17770336 | -0.296 | 0.085 | 5.11E-04 |
| rs1857883 | -0.301 | 0.085 | 4.22E-04 |
| rs1893659 | -0.293 | 0.085 | 5.67E-04 |
| rs1927626 | -0.298 | 0.085 | 4.76E-04 |
| rs1928496 | -0.300 | 0.085 | 4.33E-04 |
| rs2002023 | -0.301 | 0.085 | 4.17E-04 |
| rs2020942 | -0.305 | 0.085 | 3.41E-04 |
| rs2038646 | -0.297 | 0.085 | 4.91E-04 |
| rs2041376 | -0.301 | 0.085 | 4.17E-04 |
| rs2043016 | -0.298 | 0.085 | 4.77E-04 |
| rs2052607 | -0.289 | 0.085 | 6.42E-04 |
| rs2124499 | -0.304 | 0.085 | 3.70E-04 |
| rs215614 | -0.301 | 0.085 | 4.27E-04 |
| rs2172131 | -0.296 | 0.085 | 4.96E-04 |
| rs217672 | -0.304 | 0.085 | 3.59E-04 |
| rs2178899 | -0.304 | 0.085 | 3.50E-04 |
| rs2192527 | -0.301 | 0.085 | 4.28E-04 |
| rs2198557 | -0.297 | 0.085 | 4.90E-04 |

| | | | |
|---|---|---|---|
| rs2216931 | -0.300 | 0.085 | 4.28E-04 |
| rs2243928 | -0.305 | 0.085 | 3.44E-04 |
| rs2279574 | -0.314 | 0.085 | 2.27E-04 |
| rs2281819 | -0.295 | 0.085 | 5.11E-04 |
| rs2287214 | -0.303 | 0.085 | 3.85E-04 |
| rs2306593 | -0.308 | 0.085 | 2.92E-04 |
| rs2370982 | -0.302 | 0.085 | 4.05E-04 |
| rs2371767 | -0.292 | 0.085 | 5.81E-04 |
| rs2374947 | -0.306 | 0.085 | 3.13E-04 |
| rs245775 | -0.304 | 0.085 | 3.64E-04 |
| rs2494196 | -0.301 | 0.086 | 4.36E-04 |
| rs2499468 | -0.302 | 0.085 | 3.98E-04 |
| rs2606228 | -0.306 | 0.085 | 3.26E-04 |
| rs2641697 | -0.295 | 0.085 | 5.12E-04 |
| rs2660241 | -0.302 | 0.085 | 3.97E-04 |
| rs2678204 | -0.293 | 0.085 | 5.57E-04 |
| rs2702123 | -0.302 | 0.085 | 3.87E-04 |
| rs2855818 | -0.288 | 0.085 | 6.65E-04 |
| rs288147 | -0.303 | 0.085 | 3.81E-04 |
| rs28893270 | -0.290 | 0.084 | 5.85E-04 |
| rs2903738 | -0.305 | 0.085 | 3.30E-04 |
| rs2914231 | -0.308 | 0.085 | 2.89E-04 |
| rs2949785 | -0.300 | 0.085 | 4.37E-04 |
| rs2954021 | -0.301 | 0.085 | 4.19E-04 |
| rs314279 | -0.297 | 0.085 | 4.93E-04 |
| rs3218036 | -0.293 | 0.085 | 5.65E-04 |
| rs33836 | -0.299 | 0.085 | 4.72E-04 |
| rs34045288 | -0.303 | 0.085 | 3.87E-04 |
| rs34483452 | -0.308 | 0.085 | 3.08E-04 |
| rs35154152 | -0.313 | 0.085 | 2.20E-04 |
| rs35243581 | -0.301 | 0.085 | 4.17E-04 |
| rs35344761 | -0.302 | 0.085 | 3.93E-04 |
| rs35787800 | -0.305 | 0.085 | 3.43E-04 |
| rs3730071 | -0.305 | 0.085 | 3.37E-04 |
| rs3736896 | -0.309 | 0.085 | 2.95E-04 |
| rs3766823 | -0.302 | 0.085 | 4.01E-04 |
| rs3796658 | -0.291 | 0.085 | 5.92E-04 |
| rs3803286 | -0.306 | 0.085 | 3.31E-04 |
| rs3817428 | -0.312 | 0.085 | 2.60E-04 |
| rs40060 | -0.307 | 0.086 | 3.26E-04 |
| rs41310284 | -0.297 | 0.085 | 4.86E-04 |
| rs41313250 | -0.297 | 0.085 | 4.78E-04 |
| rs429358 | -0.305 | 0.085 | 3.53E-04 |
| rs4450871 | -0.299 | 0.085 | 4.40E-04 |
| rs4547132 | -0.301 | 0.085 | 4.07E-04 |

| | | | |
|---|---|---|---|
| rs4549685 | -0.299 | 0.085 | 4.57E-04 |
| rs4690324 | -0.297 | 0.085 | 4.79E-04 |
| rs4709745 | -0.301 | 0.085 | 4.23E-04 |
| rs4718964 | -0.301 | 0.085 | 4.26E-04 |
| rs4722398 | -0.311 | 0.084 | 2.32E-04 |
| rs4759318 | -0.305 | 0.085 | 3.43E-04 |
| rs4790841 | -0.302 | 0.085 | 4.00E-04 |
| rs4820325 | -0.307 | 0.085 | 3.20E-04 |
| rs4864201 | -0.302 | 0.085 | 4.02E-04 |
| rs4876611 | -0.297 | 0.085 | 5.01E-04 |
| rs4981693 | -0.302 | 0.085 | 3.94E-04 |
| rs501038 | -0.309 | 0.085 | 2.67E-04 |
| rs538656 | -0.294 | 0.085 | 5.75E-04 |
| rs543874 | -0.317 | 0.086 | 2.11E-04 |
| rs56094641 | -0.295 | 0.086 | 6.49E-04 |
| rs56254492 | -0.303 | 0.085 | 3.72E-04 |
| rs57221746 | -0.300 | 0.085 | 4.38E-04 |
| rs57636386 | -0.299 | 0.085 | 4.62E-04 |
| rs57866767 | -0.306 | 0.085 | 3.37E-04 |
| rs58125425 | -0.302 | 0.085 | 3.99E-04 |
| rs58360798 | -0.297 | 0.085 | 4.75E-04 |
| rs59066241 | -0.303 | 0.085 | 3.80E-04 |
| rs6021948 | -0.298 | 0.085 | 4.63E-04 |
| rs6029180 | -0.298 | 0.085 | 4.62E-04 |
| rs62025831 | -0.302 | 0.086 | 4.20E-04 |
| rs62037365 | -0.285 | 0.085 | 8.48E-04 |
| rs62190394 | -0.295 | 0.085 | 5.29E-04 |
| rs62261725 | -0.294 | 0.085 | 5.31E-04 |
| rs62285233 | -0.298 | 0.085 | 4.63E-04 |
| rs6545714 | -0.306 | 0.085 | 3.40E-04 |
| rs6575340 | -0.300 | 0.085 | 4.33E-04 |
| rs6693294 | -0.302 | 0.085 | 4.00E-04 |
| rs6696828 | -0.308 | 0.085 | 2.80E-04 |
| rs6699744 | -0.297 | 0.085 | 4.95E-04 |
| rs6717858 | -0.297 | 0.086 | 5.15E-04 |
| rs67689854 | -0.300 | 0.085 | 4.22E-04 |
| rs6847975 | -0.290 | 0.084 | 5.96E-04 |
| rs6861157 | -0.303 | 0.085 | 3.79E-04 |
| rs6925197 | -0.295 | 0.085 | 5.06E-04 |
| rs6948959 | -0.300 | 0.085 | 4.37E-04 |
| rs6973656 | -0.305 | 0.085 | 3.45E-04 |
| rs704255 | -0.303 | 0.085 | 3.87E-04 |
| rs7080838 | -0.297 | 0.085 | 4.86E-04 |
| rs7124681 | -0.303 | 0.086 | 4.09E-04 |
| rs7133378 | -0.304 | 0.085 | 3.68E-04 |

| | | | |
|---|---|---|---|
| rs713586 | -0.298 | 0.086 | 5.28E-04 |
| rs71658797 | -0.300 | 0.085 | 4.44E-04 |
| rs7171864 | -0.308 | 0.085 | 2.96E-04 |
| rs72755233 | -0.306 | 0.085 | 3.05E-04 |
| rs72917533 | -0.301 | 0.085 | 4.13E-04 |
| rs72976986 | -0.290 | 0.084 | 5.90E-04 |
| rs72995085 | -0.301 | 0.085 | 4.21E-04 |
| rs73041988 | -0.301 | 0.085 | 4.18E-04 |
| rs73213501 | -0.300 | 0.085 | 4.28E-04 |
| rs73307079 | -0.303 | 0.085 | 3.85E-04 |
| rs75192636 | -0.303 | 0.085 | 3.86E-04 |
| rs75641275 | -0.300 | 0.085 | 4.38E-04 |
| rs76040172 | -0.293 | 0.085 | 5.51E-04 |
| rs7613261 | -0.303 | 0.085 | 3.79E-04 |
| rs76345589 | -0.303 | 0.085 | 3.72E-04 |
| rs7637852 | -0.299 | 0.085 | 4.51E-04 |
| rs764729 | -0.307 | 0.085 | 2.97E-04 |
| rs7649970 | -0.304 | 0.086 | 3.80E-04 |
| rs768023 | -0.306 | 0.085 | 3.21E-04 |
| rs7845090 | -0.303 | 0.085 | 3.91E-04 |
| rs78801969 | -0.303 | 0.085 | 3.64E-04 |
| rs79535757 | -0.298 | 0.085 | 4.65E-04 |
| rs7972728 | -0.309 | 0.085 | 2.80E-04 |
| rs7982447 | -0.299 | 0.085 | 4.47E-04 |
| rs80135947 | -0.285 | 0.085 | 7.63E-04 |
| rs8042404 | -0.303 | 0.085 | 3.84E-04 |
| rs8087074 | -0.305 | 0.085 | 3.37E-04 |
| rs809955 | -0.299 | 0.085 | 4.49E-04 |
| rs815611 | -0.300 | 0.085 | 4.28E-04 |
| rs879620 | -0.291 | 0.085 | 6.21E-04 |
| rs9289630 | -0.301 | 0.085 | 4.20E-04 |
| rs9320823 | -0.297 | 0.085 | 4.99E-04 |
| rs9358912 | -0.286 | 0.085 | 8.09E-04 |
| rs952227 | -0.301 | 0.085 | 4.22E-04 |
| rs9551991 | -0.299 | 0.085 | 4.51E-04 |
| rs9584855 | -0.305 | 0.085 | 3.39E-04 |
| rs9657541 | -0.305 | 0.085 | 3.38E-04 |
| rs972283 | -0.309 | 0.085 | 2.94E-04 |
| rs9843653 | -0.310 | 0.085 | 2.77E-04 |
| rs9883422 | -0.295 | 0.085 | 5.08E-04 |
| All | -0.301 | 0.085 | 3.95E-04 |

**Forced vital capacity (FVC) Best measure || id:UKB-a:232**

| SNP | b | se | p |
|---|---|---|---|
| rs10171807 | 0.443 | 0.129 | 5.86E-04 |
| rs10202701 | 0.433 | 0.129 | 7.61E-04 |
| rs10283100 | 0.442 | 0.129 | 6.02E-04 |
| rs10410606 | 0.452 | 0.128 | 4.28E-04 |
| rs1046934 | 0.442 | 0.129 | 6.04E-04 |
| rs10477538 | 0.444 | 0.129 | 5.65E-04 |
| rs10487888 | 0.444 | 0.129 | 5.62E-04 |
| rs10489880 | 0.445 | 0.129 | 5.46E-04 |
| rs10753570 | 0.437 | 0.129 | 6.82E-04 |
| rs10797116 | 0.434 | 0.129 | 7.51E-04 |
| rs10821713 | 0.440 | 0.129 | 6.33E-04 |
| rs10852940 | 0.438 | 0.129 | 6.72E-04 |
| rs10916137 | 0.445 | 0.129 | 5.40E-04 |
| rs11083092 | 0.439 | 0.129 | 6.52E-04 |
| rs11098727 | 0.447 | 0.129 | 5.06E-04 |
| rs111251222 | 0.440 | 0.129 | 6.47E-04 |
| rs113454775 | 0.430 | 0.128 | 8.02E-04 |
| rs1136287 | 0.447 | 0.129 | 5.24E-04 |
| rs11645286 | 0.443 | 0.129 | 5.76E-04 |
| rs117372259 | 0.443 | 0.129 | 5.77E-04 |
| rs11867847 | 0.447 | 0.129 | 5.13E-04 |
| rs11967850 | 0.436 | 0.129 | 6.99E-04 |
| rs12144044 | 0.439 | 0.129 | 6.54E-04 |
| rs1232604 | 0.438 | 0.129 | 6.66E-04 |
| rs12368149 | 0.446 | 0.129 | 5.35E-04 |
| rs12452590 | 0.444 | 0.129 | 5.64E-04 |
| rs12509305 | 0.439 | 0.129 | 6.60E-04 |
| rs12571363 | 0.446 | 0.129 | 5.50E-04 |
| rs12620308 | 0.444 | 0.129 | 5.74E-04 |
| rs12713685 | 0.443 | 0.129 | 5.89E-04 |
| rs12798028 | 0.442 | 0.129 | 6.08E-04 |
| rs12894067 | 0.439 | 0.129 | 6.63E-04 |
| rs12917449 | 0.441 | 0.129 | 6.22E-04 |
| rs13146142 | 0.389 | 0.129 | 2.59E-03 |
| rs13156484 | 0.447 | 0.129 | 5.19E-04 |
| rs13229167 | 0.437 | 0.129 | 7.14E-04 |
| rs13433809 | 0.446 | 0.129 | 5.38E-04 |
| rs1366594 | 0.438 | 0.129 | 6.79E-04 |
| rs140988776 | 0.438 | 0.129 | 6.78E-04 |
| rs1415701 | 0.449 | 0.129 | 5.04E-04 |
| rs1417488 | 0.448 | 0.129 | 5.17E-04 |
| rs143384 | 0.440 | 0.129 | 6.49E-04 |

| | | | |
|---|---|---|---|
| rs1438909 | 0.446 | 0.129 | 5.26E-04 |
| rs1460039 | 0.444 | 0.129 | 5.68E-04 |
| rs147110934 | 0.448 | 0.129 | 5.02E-04 |
| rs1473527 | 0.459 | 0.128 | 3.35E-04 |
| rs1490384 | 0.428 | 0.130 | 1.02E-03 |
| rs149542 | 0.438 | 0.129 | 6.76E-04 |
| rs1509310 | 0.432 | 0.128 | 7.67E-04 |
| rs1516890 | 0.439 | 0.129 | 6.56E-04 |
| rs1553521 | 0.443 | 0.129 | 5.91E-04 |
| rs17647719 | 0.444 | 0.129 | 5.71E-04 |
| rs17855988 | 0.438 | 0.129 | 6.67E-04 |
| rs1991556 | 0.266 | 0.097 | 5.86E-03 |
| rs2093210 | 0.429 | 0.128 | 8.12E-04 |
| rs2131354 | 0.461 | 0.129 | 3.38E-04 |
| rs2131371 | 0.445 | 0.129 | 5.49E-04 |
| rs2167734 | 0.431 | 0.128 | 7.77E-04 |
| rs2186288 | 0.444 | 0.129 | 5.62E-04 |
| rs2194411 | 0.453 | 0.128 | 4.07E-04 |
| rs2286425 | 0.430 | 0.129 | 8.11E-04 |
| rs2294214 | 0.449 | 0.129 | 4.93E-04 |
| rs2311313 | 0.440 | 0.129 | 6.38E-04 |
| rs2364232 | 0.451 | 0.129 | 4.78E-04 |
| rs240483 | 0.443 | 0.129 | 5.78E-04 |
| rs254577 | 0.452 | 0.128 | 4.17E-04 |
| rs2579762 | 0.437 | 0.129 | 6.89E-04 |
| rs2610986 | 0.448 | 0.129 | 5.00E-04 |
| rs2631676 | 0.442 | 0.129 | 6.00E-04 |
| rs2678408 | 0.434 | 0.129 | 7.53E-04 |
| rs2717329 | 0.446 | 0.129 | 5.40E-04 |
| rs2812208 | 0.446 | 0.130 | 6.05E-04 |
| rs2833 | 0.446 | 0.129 | 5.37E-04 |
| rs28391281 | 0.441 | 0.129 | 6.06E-04 |
| rs28427480 | 0.446 | 0.129 | 5.55E-04 |
| rs28446901 | 0.442 | 0.129 | 5.94E-04 |
| rs2854152 | 0.437 | 0.129 | 7.22E-04 |
| rs2885697 | 0.445 | 0.129 | 5.76E-04 |
| rs28925904 | 0.440 | 0.129 | 6.22E-04 |
| rs28929474 | 0.452 | 0.129 | 4.42E-04 |
| rs3019020 | 0.433 | 0.128 | 7.44E-04 |
| rs309137 | 0.467 | 0.124 | 1.71E-04 |
| rs3116605 | 0.444 | 0.129 | 5.66E-04 |
| rs314276 | 0.448 | 0.129 | 4.93E-04 |
| rs3213183 | 0.441 | 0.129 | 6.15E-04 |
| rs3253 | 0.447 | 0.129 | 5.22E-04 |
| rs34104813 | 0.460 | 0.128 | 3.40E-04 |

| | | | |
|---|---|---|---|
| rs34517439 | 0.444 | 0.128 | 5.55E-04 |
| rs34810766 | 0.444 | 0.129 | 5.70E-04 |
| rs35344761 | 0.442 | 0.129 | 6.02E-04 |
| rs35506 | 0.438 | 0.129 | 6.67E-04 |
| rs36226649 | 0.445 | 0.129 | 5.45E-04 |
| rs3774219 | 0.447 | 0.129 | 5.10E-04 |
| rs3790085 | 0.441 | 0.129 | 6.61E-04 |
| rs3924298 | 0.440 | 0.129 | 6.43E-04 |
| rs4073717 | 0.446 | 0.129 | 5.36E-04 |
| rs4128846 | 0.441 | 0.129 | 6.16E-04 |
| rs42031 | 0.447 | 0.129 | 5.12E-04 |
| rs4320932 | 0.438 | 0.129 | 6.63E-04 |
| rs4413296 | 0.436 | 0.129 | 7.02E-04 |
| rs4659427 | 0.438 | 0.129 | 6.77E-04 |
| rs4670549 | 0.443 | 0.129 | 5.91E-04 |
| rs4804519 | 0.439 | 0.129 | 6.58E-04 |
| rs4809327 | 0.444 | 0.129 | 5.77E-04 |
| rs4852252 | 0.443 | 0.129 | 5.86E-04 |
| rs4902684 | 0.439 | 0.129 | 6.53E-04 |
| rs4909296 | 0.443 | 0.129 | 5.98E-04 |
| rs4938337 | 0.442 | 0.129 | 5.86E-04 |
| rs4951407 | 0.439 | 0.129 | 6.65E-04 |
| rs4984992 | 0.436 | 0.129 | 7.08E-04 |
| rs550245 | 0.444 | 0.129 | 5.62E-04 |
| rs555554 | 0.438 | 0.129 | 6.75E-04 |
| rs55654976 | 0.434 | 0.129 | 7.37E-04 |
| rs55681913 | 0.445 | 0.129 | 5.39E-04 |
| rs55811836 | 0.440 | 0.129 | 6.59E-04 |
| rs56318008 | 0.440 | 0.129 | 6.39E-04 |
| rs575837 | 0.443 | 0.129 | 5.93E-04 |
| rs59985551 | 0.484 | 0.129 | 1.81E-04 |
| rs6066104 | 0.441 | 0.129 | 6.23E-04 |
| rs62115563 | 0.442 | 0.129 | 5.92E-04 |
| rs62276616 | 0.440 | 0.129 | 6.31E-04 |
| rs62437411 | 0.443 | 0.129 | 5.81E-04 |
| rs6709833 | 0.441 | 0.129 | 6.20E-04 |
| rs6729308 | 0.444 | 0.129 | 5.65E-04 |
| rs6742854 | 0.435 | 0.129 | 7.18E-04 |
| rs6762578 | 0.447 | 0.129 | 5.17E-04 |
| rs67672535 | 0.438 | 0.129 | 6.82E-04 |
| rs6820254 | 0.448 | 0.129 | 5.04E-04 |
| rs6902771 | 0.442 | 0.129 | 6.17E-04 |
| rs6907284 | 0.443 | 0.129 | 5.78E-04 |
| rs6937121 | 0.452 | 0.129 | 4.66E-04 |
| rs6977081 | 0.446 | 0.129 | 5.30E-04 |

| | | | |
|---|---|---|---|
| rs7029491 | 0.435 | 0.129 | 7.22E-04 |
| rs7143806 | 0.449 | 0.129 | 4.83E-04 |
| rs7176223 | 0.454 | 0.128 | 3.95E-04 |
| rs7210583 | 0.446 | 0.129 | 5.41E-04 |
| rs723588 | 0.439 | 0.129 | 6.56E-04 |
| rs72721175 | 0.452 | 0.129 | 4.33E-04 |
| rs72755233 | 0.435 | 0.129 | 7.32E-04 |
| rs72776502 | 0.460 | 0.128 | 3.40E-04 |
| rs72964584 | 0.449 | 0.128 | 4.71E-04 |
| rs72976986 | 0.430 | 0.128 | 7.79E-04 |
| rs735830 | 0.443 | 0.129 | 6.04E-04 |
| rs7443323 | 0.449 | 0.129 | 4.91E-04 |
| rs75390002 | 0.437 | 0.129 | 6.72E-04 |
| rs7594073 | 0.439 | 0.129 | 6.56E-04 |
| rs7680140 | 0.442 | 0.129 | 6.10E-04 |
| rs77328429 | 0.436 | 0.129 | 6.94E-04 |
| rs773652 | 0.437 | 0.129 | 6.95E-04 |
| rs774736 | 0.440 | 0.129 | 6.40E-04 |
| rs7775683 | 0.457 | 0.127 | 3.40E-04 |
| rs7815955 | 0.429 | 0.129 | 8.72E-04 |
| rs78378222 | 0.438 | 0.128 | 6.35E-04 |
| rs78524772 | 0.445 | 0.129 | 5.37E-04 |
| rs7854962 | 0.433 | 0.129 | 7.67E-04 |
| rs79409628 | 0.444 | 0.129 | 5.82E-04 |
| rs7952436 | 0.449 | 0.128 | 4.77E-04 |
| rs7968682 | 0.447 | 0.129 | 5.49E-04 |
| rs798490 | 0.437 | 0.129 | 7.00E-04 |
| rs8002779 | 0.454 | 0.128 | 4.03E-04 |
| rs8037675 | 0.442 | 0.129 | 6.06E-04 |
| rs8041080 | 0.443 | 0.129 | 6.04E-04 |
| rs828907 | 0.441 | 0.129 | 6.25E-04 |
| rs830643 | 0.444 | 0.129 | 5.70E-04 |
| rs841217 | 0.435 | 0.129 | 7.10E-04 |
| rs871632 | 0.440 | 0.129 | 6.47E-04 |
| rs876531 | 0.441 | 0.129 | 6.33E-04 |
| rs9268118 | 0.424 | 0.128 | 9.68E-04 |
| rs9321170 | 0.446 | 0.129 | 5.37E-04 |
| rs9379084 | 0.443 | 0.129 | 5.92E-04 |
| rs9435733 | 0.437 | 0.130 | 7.43E-04 |
| rs9442580 | 0.439 | 0.129 | 6.49E-04 |
| rs9462085 | 0.440 | 0.129 | 6.50E-04 |
| rs9542595 | 0.445 | 0.129 | 5.51E-04 |
| rs9696116 | 0.455 | 0.129 | 4.11E-04 |
| rs9809116 | 0.430 | 0.128 | 7.98E-04 |
| rs9865460 | 0.438 | 0.129 | 6.98E-04 |

| | | | |
|---|---|---|---|
| rs9912553 | 0.431 | 0.129 | 8.22E-04 |
| rs9988963 | 0.435 | 0.129 | 7.15E-04 |
| All | 0.441 | 0.128 | 5.80E-04 |

**Trunk fat percentage || id:UKB-a:290**

| SNP | b | se | p |
|---|---|---|---|
| rs10100245 | -0.302 | 0.085 | 4.00E-04 |
| rs10172196 | -0.303 | 0.085 | 3.76E-04 |
| rs10209821 | -0.304 | 0.085 | 3.64E-04 |
| rs10259620 | -0.309 | 0.085 | 2.71E-04 |
| rs10423928 | -0.304 | 0.085 | 3.67E-04 |
| rs10499014 | -0.301 | 0.085 | 4.22E-04 |
| rs10514136 | -0.295 | 0.085 | 5.06E-04 |
| rs1056441 | -0.302 | 0.085 | 3.93E-04 |
| rs10732335 | -0.291 | 0.085 | 6.43E-04 |
| rs10756798 | -0.309 | 0.085 | 2.77E-04 |
| rs10830566 | -0.303 | 0.085 | 3.82E-04 |
| rs10848835 | -0.301 | 0.085 | 4.12E-04 |
| rs10938397 | -0.310 | 0.086 | 2.90E-04 |
| rs10947793 | -0.300 | 0.085 | 4.25E-04 |
| rs10954772 | -0.305 | 0.085 | 3.37E-04 |
| rs10999460 | -0.293 | 0.085 | 5.78E-04 |
| rs11012732 | -0.284 | 0.085 | 8.17E-04 |
| rs11030119 | -0.295 | 0.085 | 5.47E-04 |
| rs11042030 | -0.302 | 0.085 | 4.09E-04 |
| rs11042725 | -0.306 | 0.085 | 3.18E-04 |
| rs11043764 | -0.297 | 0.085 | 4.83E-04 |
| rs11150745 | -0.299 | 0.085 | 4.49E-04 |
| rs111743285 | -0.295 | 0.085 | 5.16E-04 |
| rs11205303 | -0.319 | 0.086 | 1.87E-04 |
| rs11222371 | -0.305 | 0.085 | 3.50E-04 |
| rs11603783 | -0.306 | 0.085 | 3.14E-04 |
| rs11666808 | -0.307 | 0.085 | 3.22E-04 |
| rs11782341 | -0.308 | 0.085 | 2.90E-04 |
| rs11786089 | -0.298 | 0.085 | 4.63E-04 |
| rs1182199 | -0.308 | 0.085 | 3.04E-04 |
| rs11856579 | -0.298 | 0.085 | 4.75E-04 |
| rs11857221 | -0.297 | 0.085 | 4.91E-04 |
| rs11866219 | -0.303 | 0.085 | 3.79E-04 |
| rs12037698 | -0.298 | 0.085 | 4.64E-04 |
| rs12103006 | -0.301 | 0.085 | 4.18E-04 |
| rs1218822 | -0.302 | 0.085 | 4.01E-04 |
| rs12339822 | -0.295 | 0.085 | 5.26E-04 |
| rs12367809 | -0.306 | 0.085 | 3.37E-04 |
| rs12475388 | -0.303 | 0.085 | 3.75E-04 |
| rs12619178 | -0.302 | 0.085 | 4.00E-04 |
| rs12716979 | -0.277 | 0.082 | 7.36E-04 |
| rs12724928 | -0.299 | 0.085 | 4.53E-04 |

| | | | |
|---|---|---|---|
| rs12757328 | -0.299 | 0.085 | 4.46E-04 |
| rs12890931 | -0.298 | 0.085 | 4.60E-04 |
| rs12901071 | -0.299 | 0.085 | 4.58E-04 |
| rs12926311 | -0.296 | 0.085 | 4.94E-04 |
| rs1293447 | -0.302 | 0.085 | 3.97E-04 |
| rs12986231 | -0.300 | 0.085 | 4.40E-04 |
| rs12992672 | -0.310 | 0.085 | 2.76E-04 |
| rs13109830 | -0.298 | 0.085 | 4.78E-04 |
| rs13135092 | -0.285 | 0.085 | 7.52E-04 |
| rs13174863 | -0.299 | 0.085 | 4.51E-04 |
| rs138556772 | -0.299 | 0.085 | 4.25E-04 |
| rs1441264 | -0.299 | 0.085 | 4.60E-04 |
| rs1446585 | -0.332 | 0.081 | 4.50E-05 |
| rs1469084 | -0.309 | 0.085 | 2.59E-04 |
| rs150215901 | -0.300 | 0.085 | 4.37E-04 |
| rs1523136 | -0.299 | 0.085 | 4.47E-04 |
| rs153664 | -0.301 | 0.085 | 4.06E-04 |
| rs1568489 | -0.308 | 0.085 | 2.92E-04 |
| rs157845 | -0.299 | 0.085 | 4.49E-04 |
| rs159961 | -0.294 | 0.085 | 5.30E-04 |
| rs16880854 | -0.306 | 0.085 | 3.30E-04 |
| rs16934748 | -0.304 | 0.085 | 3.48E-04 |
| rs16951304 | -0.302 | 0.085 | 4.13E-04 |
| rs17024258 | -0.302 | 0.085 | 4.00E-04 |
| rs1724557 | -0.302 | 0.085 | 3.91E-04 |
| rs17265513 | -0.310 | 0.085 | 2.67E-04 |
| rs17522122 | -0.304 | 0.085 | 3.58E-04 |
| rs17770336 | -0.296 | 0.085 | 5.11E-04 |
| rs1857883 | -0.301 | 0.085 | 4.22E-04 |
| rs1893659 | -0.293 | 0.085 | 5.67E-04 |
| rs1927626 | -0.298 | 0.085 | 4.76E-04 |
| rs1928496 | -0.300 | 0.085 | 4.33E-04 |
| rs2002023 | -0.301 | 0.085 | 4.17E-04 |
| rs2020942 | -0.305 | 0.085 | 3.41E-04 |
| rs2038646 | -0.297 | 0.085 | 4.91E-04 |
| rs2041376 | -0.301 | 0.085 | 4.17E-04 |
| rs2043016 | -0.298 | 0.085 | 4.77E-04 |
| rs2052607 | -0.289 | 0.085 | 6.42E-04 |
| rs2124499 | -0.304 | 0.085 | 3.70E-04 |
| rs215614 | -0.301 | 0.085 | 4.27E-04 |
| rs2172131 | -0.296 | 0.085 | 4.96E-04 |
| rs217672 | -0.304 | 0.085 | 3.59E-04 |
| rs2178899 | -0.304 | 0.085 | 3.50E-04 |
| rs2192527 | -0.301 | 0.085 | 4.28E-04 |
| rs2198557 | -0.297 | 0.085 | 4.90E-04 |

| | | | |
|---|---|---|---|
| rs2216931 | -0.300 | 0.085 | 4.28E-04 |
| rs2243928 | -0.305 | 0.085 | 3.44E-04 |
| rs2279574 | -0.314 | 0.085 | 2.27E-04 |
| rs2281819 | -0.295 | 0.085 | 5.11E-04 |
| rs2287214 | -0.303 | 0.085 | 3.85E-04 |
| rs2306593 | -0.308 | 0.085 | 2.92E-04 |
| rs2370982 | -0.302 | 0.085 | 4.05E-04 |
| rs2371767 | -0.292 | 0.085 | 5.81E-04 |
| rs2374947 | -0.306 | 0.085 | 3.13E-04 |
| rs245775 | -0.304 | 0.085 | 3.64E-04 |
| rs2494196 | -0.301 | 0.086 | 4.36E-04 |
| rs2499468 | -0.302 | 0.085 | 3.98E-04 |
| rs2606228 | -0.306 | 0.085 | 3.26E-04 |
| rs2641697 | -0.295 | 0.085 | 5.12E-04 |
| rs2660241 | -0.302 | 0.085 | 3.97E-04 |
| rs2678204 | -0.293 | 0.085 | 5.57E-04 |
| rs2702123 | -0.302 | 0.085 | 3.87E-04 |
| rs2855818 | -0.288 | 0.085 | 6.65E-04 |
| rs288147 | -0.303 | 0.085 | 3.81E-04 |
| rs28893270 | -0.290 | 0.084 | 5.85E-04 |
| rs2903738 | -0.305 | 0.085 | 3.30E-04 |
| rs2914231 | -0.308 | 0.085 | 2.89E-04 |
| rs2949785 | -0.300 | 0.085 | 4.37E-04 |
| rs2954021 | -0.301 | 0.085 | 4.19E-04 |
| rs314279 | -0.297 | 0.085 | 4.93E-04 |
| rs3218036 | -0.293 | 0.085 | 5.65E-04 |
| rs33836 | -0.299 | 0.085 | 4.72E-04 |
| rs34045288 | -0.303 | 0.085 | 3.87E-04 |
| rs34483452 | -0.308 | 0.085 | 3.08E-04 |
| rs35154152 | -0.313 | 0.085 | 2.20E-04 |
| rs35243581 | -0.301 | 0.085 | 4.17E-04 |
| rs35344761 | -0.302 | 0.085 | 3.93E-04 |
| rs35787800 | -0.305 | 0.085 | 3.43E-04 |
| rs3730071 | -0.305 | 0.085 | 3.37E-04 |
| rs3736896 | -0.309 | 0.085 | 2.95E-04 |
| rs3766823 | -0.302 | 0.085 | 4.01E-04 |
| rs3796658 | -0.291 | 0.085 | 5.92E-04 |
| rs3803286 | -0.306 | 0.085 | 3.31E-04 |
| rs3817428 | -0.312 | 0.085 | 2.60E-04 |
| rs40060 | -0.307 | 0.086 | 3.26E-04 |
| rs41310284 | -0.297 | 0.085 | 4.86E-04 |
| rs41313250 | -0.297 | 0.085 | 4.78E-04 |
| rs429358 | -0.305 | 0.085 | 3.53E-04 |
| rs4450871 | -0.299 | 0.085 | 4.40E-04 |
| rs4547132 | -0.301 | 0.085 | 4.07E-04 |

| | | | |
|---|---|---|---|
| rs4549685 | -0.299 | 0.085 | 4.57E-04 |
| rs4690324 | -0.297 | 0.085 | 4.79E-04 |
| rs4709745 | -0.301 | 0.085 | 4.23E-04 |
| rs4718964 | -0.301 | 0.085 | 4.26E-04 |
| rs4722398 | -0.311 | 0.084 | 2.32E-04 |
| rs4759318 | -0.305 | 0.085 | 3.43E-04 |
| rs4790841 | -0.302 | 0.085 | 4.00E-04 |
| rs4820325 | -0.307 | 0.085 | 3.20E-04 |
| rs4864201 | -0.302 | 0.085 | 4.02E-04 |
| rs4876611 | -0.297 | 0.085 | 5.01E-04 |
| rs4981693 | -0.302 | 0.085 | 3.94E-04 |
| rs501038 | -0.309 | 0.085 | 2.67E-04 |
| rs538656 | -0.294 | 0.085 | 5.75E-04 |
| rs543874 | -0.317 | 0.086 | 2.11E-04 |
| rs56094641 | -0.295 | 0.086 | 6.49E-04 |
| rs56254492 | -0.303 | 0.085 | 3.72E-04 |
| rs57221746 | -0.300 | 0.085 | 4.38E-04 |
| rs57636386 | -0.299 | 0.085 | 4.62E-04 |
| rs57866767 | -0.306 | 0.085 | 3.37E-04 |
| rs58125425 | -0.302 | 0.085 | 3.99E-04 |
| rs58360798 | -0.297 | 0.085 | 4.75E-04 |
| rs59066241 | -0.303 | 0.085 | 3.80E-04 |
| rs6021948 | -0.298 | 0.085 | 4.63E-04 |
| rs6029180 | -0.298 | 0.085 | 4.62E-04 |
| rs62025831 | -0.302 | 0.086 | 4.20E-04 |
| rs62037365 | -0.285 | 0.085 | 8.48E-04 |
| rs62190394 | -0.295 | 0.085 | 5.29E-04 |
| rs62261725 | -0.294 | 0.085 | 5.31E-04 |
| rs62285233 | -0.298 | 0.085 | 4.63E-04 |
| rs6545714 | -0.306 | 0.085 | 3.40E-04 |
| rs6575340 | -0.300 | 0.085 | 4.33E-04 |
| rs6693294 | -0.302 | 0.085 | 4.00E-04 |
| rs6696828 | -0.308 | 0.085 | 2.80E-04 |
| rs6699744 | -0.297 | 0.085 | 4.95E-04 |
| rs6717858 | -0.297 | 0.086 | 5.15E-04 |
| rs67689854 | -0.300 | 0.085 | 4.22E-04 |
| rs6847975 | -0.290 | 0.084 | 5.96E-04 |
| rs6861157 | -0.303 | 0.085 | 3.79E-04 |
| rs6925197 | -0.295 | 0.085 | 5.06E-04 |
| rs6948959 | -0.300 | 0.085 | 4.37E-04 |
| rs6973656 | -0.305 | 0.085 | 3.45E-04 |
| rs704255 | -0.303 | 0.085 | 3.87E-04 |
| rs7080838 | -0.297 | 0.085 | 4.86E-04 |
| rs7124681 | -0.303 | 0.086 | 4.09E-04 |
| rs7133378 | -0.304 | 0.085 | 3.68E-04 |

| | | | |
|---|---|---|---|
| rs713586 | -0.298 | 0.086 | 5.28E-04 |
| rs71658797 | -0.300 | 0.085 | 4.44E-04 |
| rs7171864 | -0.308 | 0.085 | 2.96E-04 |
| rs72755233 | -0.306 | 0.085 | 3.05E-04 |
| rs72917533 | -0.301 | 0.085 | 4.13E-04 |
| rs72976986 | -0.290 | 0.084 | 5.90E-04 |
| rs72995085 | -0.301 | 0.085 | 4.21E-04 |
| rs73041988 | -0.301 | 0.085 | 4.18E-04 |
| rs73213501 | -0.300 | 0.085 | 4.28E-04 |
| rs73307079 | -0.303 | 0.085 | 3.85E-04 |
| rs75192636 | -0.303 | 0.085 | 3.86E-04 |
| rs75641275 | -0.300 | 0.085 | 4.38E-04 |
| rs76040172 | -0.293 | 0.085 | 5.51E-04 |
| rs7613261 | -0.303 | 0.085 | 3.79E-04 |
| rs76345589 | -0.303 | 0.085 | 3.72E-04 |
| rs7637852 | -0.299 | 0.085 | 4.51E-04 |
| rs764729 | -0.307 | 0.085 | 2.97E-04 |
| rs7649970 | -0.304 | 0.086 | 3.80E-04 |
| rs768023 | -0.306 | 0.085 | 3.21E-04 |
| rs7845090 | -0.303 | 0.085 | 3.91E-04 |
| rs78801969 | -0.303 | 0.085 | 3.64E-04 |
| rs79535757 | -0.298 | 0.085 | 4.65E-04 |
| rs7972728 | -0.309 | 0.085 | 2.80E-04 |
| rs7982447 | -0.299 | 0.085 | 4.47E-04 |
| rs80135947 | -0.285 | 0.085 | 7.63E-04 |
| rs8042404 | -0.303 | 0.085 | 3.84E-04 |
| rs8087074 | -0.305 | 0.085 | 3.37E-04 |
| rs809955 | -0.299 | 0.085 | 4.49E-04 |
| rs815611 | -0.300 | 0.085 | 4.28E-04 |
| rs879620 | -0.291 | 0.085 | 6.21E-04 |
| rs9289630 | -0.301 | 0.085 | 4.20E-04 |
| rs9320823 | -0.297 | 0.085 | 4.99E-04 |
| rs9358912 | -0.286 | 0.085 | 8.09E-04 |
| rs952227 | -0.301 | 0.085 | 4.22E-04 |
| rs9551991 | -0.299 | 0.085 | 4.51E-04 |
| rs9584855 | -0.305 | 0.085 | 3.39E-04 |
| rs9657541 | -0.305 | 0.085 | 3.38E-04 |
| rs972283 | -0.309 | 0.085 | 2.94E-04 |
| rs9843653 | -0.310 | 0.085 | 2.77E-04 |
| rs9883422 | -0.295 | 0.085 | 5.08E-04 |
| All | -0.301 | 0.085 | 3.95E-04 |

**Arm fat percentage (right) || id:UKB-a:282**

| SNP | b | se | p |
|---|---|---|---|
| rs10064669 | -0.429 | 0.098 | 1.12E-05 |
| rs10100245 | -0.428 | 0.098 | 1.23E-05 |
| rs10245306 | -0.433 | 0.097 | 8.42E-06 |
| rs10278040 | -0.425 | 0.098 | 1.37E-05 |
| rs10423928 | -0.431 | 0.098 | 1.09E-05 |
| rs10459574 | -0.434 | 0.097 | 8.49E-06 |
| rs10745787 | -0.422 | 0.098 | 1.52E-05 |
| rs10754132 | -0.423 | 0.098 | 1.50E-05 |
| rs10756714 | -0.414 | 0.098 | 2.28E-05 |
| rs10756798 | -0.437 | 0.097 | 7.33E-06 |
| rs10774018 | -0.425 | 0.098 | 1.38E-05 |
| rs1078455 | -0.433 | 0.097 | 8.73E-06 |
| rs10938398 | -0.436 | 0.098 | 8.38E-06 |
| rs10954772 | -0.431 | 0.098 | 1.00E-05 |
| rs10999460 | -0.418 | 0.097 | 1.75E-05 |
| rs11012732 | -0.407 | 0.097 | 2.89E-05 |
| rs1101563 | -0.418 | 0.097 | 1.76E-05 |
| rs11030119 | -0.418 | 0.098 | 1.96E-05 |
| rs11042030 | -0.427 | 0.098 | 1.27E-05 |
| rs11150745 | -0.424 | 0.098 | 1.47E-05 |
| rs111625081 | -0.423 | 0.098 | 1.50E-05 |
| rs11165643 | -0.422 | 0.098 | 1.57E-05 |
| rs11205277 | -0.441 | 0.097 | 5.96E-06 |
| rs1128249 | -0.421 | 0.098 | 1.70E-05 |
| rs113019802 | -0.417 | 0.097 | 1.83E-05 |
| rs113230003 | -0.425 | 0.098 | 1.37E-05 |
| rs11609659 | -0.435 | 0.097 | 7.57E-06 |
| rs11642015 | -0.423 | 0.100 | 2.36E-05 |
| rs11668971 | -0.427 | 0.098 | 1.28E-05 |
| rs11782341 | -0.435 | 0.097 | 7.80E-06 |
| rs11786089 | -0.423 | 0.098 | 1.50E-05 |
| rs11824092 | -0.435 | 0.097 | 7.98E-06 |
| rs11839227 | -0.419 | 0.097 | 1.69E-05 |
| rs11857221 | -0.420 | 0.098 | 1.66E-05 |
| rs11866219 | -0.430 | 0.098 | 1.11E-05 |
| rs12037698 | -0.422 | 0.098 | 1.54E-05 |
| rs12072739 | -0.419 | 0.097 | 1.68E-05 |
| rs12140153 | -0.433 | 0.098 | 9.40E-06 |
| rs12144626 | -0.435 | 0.097 | 7.57E-06 |
| rs12339822 | -0.418 | 0.098 | 1.84E-05 |
| rs12367809 | -0.434 | 0.098 | 9.50E-06 |
| rs12578952 | -0.421 | 0.098 | 1.60E-05 |

| | | | |
|---|---|---|---|
| rs12614861 | -0.419 | 0.097 | 1.68E-05 |
| rs12619178 | -0.427 | 0.098 | 1.24E-05 |
| rs12679106 | -0.430 | 0.098 | 1.09E-05 |
| rs1293446 | -0.430 | 0.098 | 1.10E-05 |
| rs1296328 | -0.424 | 0.098 | 1.43E-05 |
| rs12997625 | -0.434 | 0.097 | 7.93E-06 |
| rs13053342 | -0.426 | 0.098 | 1.30E-05 |
| rs13056506 | -0.431 | 0.098 | 1.03E-05 |
| rs13114014 | -0.426 | 0.098 | 1.31E-05 |
| rs13135092 | -0.406 | 0.097 | 2.94E-05 |
| rs13174863 | -0.423 | 0.098 | 1.49E-05 |
| rs1321455 | -0.432 | 0.098 | 9.84E-06 |
| rs1322842 | -0.427 | 0.098 | 1.24E-05 |
| rs1357079 | -0.432 | 0.097 | 9.03E-06 |
| rs1376286 | -0.430 | 0.098 | 1.05E-05 |
| rs1441264 | -0.423 | 0.098 | 1.54E-05 |
| rs1446585 | -0.461 | 0.092 | 6.09E-07 |
| rs1451533 | -0.427 | 0.098 | 1.23E-05 |
| rs1503526 | -0.427 | 0.098 | 1.26E-05 |
| rs1513475 | -0.420 | 0.098 | 1.66E-05 |
| rs1516725 | -0.423 | 0.098 | 1.53E-05 |
| rs153664 | -0.427 | 0.098 | 1.27E-05 |
| rs1652376 | -0.418 | 0.097 | 1.78E-05 |
| rs17024393 | -0.428 | 0.098 | 1.27E-05 |
| rs17342242 | -0.427 | 0.098 | 1.27E-05 |
| rs17835600 | -0.427 | 0.098 | 1.24E-05 |
| rs1927626 | -0.423 | 0.098 | 1.55E-05 |
| rs1928496 | -0.425 | 0.098 | 1.38E-05 |
| rs2002023 | -0.426 | 0.098 | 1.31E-05 |
| rs2020942 | -0.431 | 0.098 | 9.76E-06 |
| rs2035936 | -0.422 | 0.098 | 1.53E-05 |
| rs2043016 | -0.422 | 0.098 | 1.53E-05 |
| rs2124499 | -0.429 | 0.098 | 1.14E-05 |
| rs2161796 | -0.411 | 0.097 | 2.11E-05 |
| rs217672 | -0.429 | 0.098 | 1.09E-05 |
| rs2239647 | -0.431 | 0.098 | 1.04E-05 |
| rs2279574 | -0.442 | 0.097 | 5.87E-06 |
| rs2306593 | -0.436 | 0.097 | 7.58E-06 |
| rs2307111 | -0.424 | 0.098 | 1.48E-05 |
| rs2318543 | -0.427 | 0.098 | 1.25E-05 |
| rs2370982 | -0.428 | 0.098 | 1.26E-05 |
| rs2384054 | -0.427 | 0.099 | 1.48E-05 |
| rs2439823 | -0.428 | 0.098 | 1.19E-05 |
| rs245775 | -0.431 | 0.098 | 1.07E-05 |
| rs2470619 | -0.432 | 0.098 | 9.69E-06 |

| | | | |
|---|---|---|---|
| rs2494196 | -0.426 | 0.098 | 1.38E-05 |
| rs2499468 | -0.428 | 0.098 | 1.23E-05 |
| rs2500406 | -0.413 | 0.097 | 1.87E-05 |
| rs2569881 | -0.432 | 0.098 | 9.58E-06 |
| rs2606228 | -0.432 | 0.097 | 9.26E-06 |
| rs2678204 | -0.416 | 0.098 | 2.03E-05 |
| rs28483178 | -0.425 | 0.098 | 1.42E-05 |
| rs2857082 | -0.419 | 0.097 | 1.62E-05 |
| rs2861685 | -0.421 | 0.098 | 1.60E-05 |
| rs2943650 | -0.425 | 0.098 | 1.42E-05 |
| rs2954021 | -0.426 | 0.098 | 1.32E-05 |
| rs301806 | -0.418 | 0.097 | 1.71E-05 |
| rs34045894 | -0.432 | 0.097 | 9.47E-06 |
| rs34331990 | -0.420 | 0.097 | 1.66E-05 |
| rs34338229 | -0.422 | 0.098 | 1.55E-05 |
| rs34417222 | -0.434 | 0.097 | 8.50E-06 |
| rs34483452 | -0.435 | 0.098 | 8.62E-06 |
| rs34580448 | -0.429 | 0.098 | 1.17E-05 |
| rs34769775 | -0.424 | 0.098 | 1.42E-05 |
| rs34811474 | -0.425 | 0.098 | 1.35E-05 |
| rs34882821 | -0.421 | 0.098 | 1.59E-05 |
| rs3734554 | -0.430 | 0.098 | 1.07E-05 |
| rs3770821 | -0.429 | 0.098 | 1.15E-05 |
| rs3771653 | -0.430 | 0.098 | 1.06E-05 |
| rs3784699 | -0.419 | 0.098 | 1.86E-05 |
| rs3803286 | -0.432 | 0.098 | 9.44E-06 |
| rs3856595 | -0.424 | 0.098 | 1.44E-05 |
| rs3923501 | -0.426 | 0.098 | 1.34E-05 |
| rs40067 | -0.426 | 0.098 | 1.35E-05 |
| rs4123668 | -0.429 | 0.098 | 1.12E-05 |
| rs4246657 | -0.429 | 0.098 | 1.13E-05 |
| rs429358 | -0.430 | 0.098 | 1.06E-05 |
| rs4402589 | -0.413 | 0.097 | 2.25E-05 |
| rs4450871 | -0.424 | 0.097 | 1.39E-05 |
| rs4477562 | -0.425 | 0.098 | 1.38E-05 |
| rs4519214 | -0.427 | 0.098 | 1.22E-05 |
| rs4547132 | -0.427 | 0.098 | 1.28E-05 |
| rs4549685 | -0.424 | 0.098 | 1.47E-05 |
| rs4727630 | -0.431 | 0.098 | 1.02E-05 |
| rs4745359 | -0.428 | 0.098 | 1.21E-05 |
| rs4777541 | -0.429 | 0.098 | 1.14E-05 |
| rs477895 | -0.425 | 0.098 | 1.39E-05 |
| rs4790841 | -0.428 | 0.098 | 1.22E-05 |
| rs4864201 | -0.427 | 0.098 | 1.25E-05 |
| rs4876611 | -0.421 | 0.098 | 1.67E-05 |

| | | | |
|---|---|---|---|
| rs538656 | -0.416 | 0.098 | 2.24E-05 |
| rs543874 | -0.452 | 0.098 | 4.14E-06 |
| rs550057 | -0.429 | 0.098 | 1.15E-05 |
| rs55714539 | -0.426 | 0.098 | 1.33E-05 |
| rs56169792 | -0.426 | 0.098 | 1.31E-05 |
| rs56375099 | -0.424 | 0.098 | 1.46E-05 |
| rs56399737 | -0.423 | 0.098 | 1.49E-05 |
| rs56803094 | -0.424 | 0.098 | 1.41E-05 |
| rs57636386 | -0.424 | 0.098 | 1.52E-05 |
| rs58120873 | -0.423 | 0.098 | 1.48E-05 |
| rs58360798 | -0.421 | 0.098 | 1.58E-05 |
| rs58824810 | -0.426 | 0.098 | 1.30E-05 |
| rs59066241 | -0.428 | 0.098 | 1.16E-05 |
| rs60764613 | -0.421 | 0.098 | 1.63E-05 |
| rs61813324 | -0.423 | 0.098 | 1.46E-05 |
| rs62037365 | -0.407 | 0.098 | 3.09E-05 |
| rs62178054 | -0.427 | 0.098 | 1.27E-05 |
| rs62246314 | -0.425 | 0.098 | 1.33E-05 |
| rs62261725 | -0.418 | 0.097 | 1.82E-05 |
| rs62477684 | -0.431 | 0.098 | 1.06E-05 |
| rs649648 | -0.431 | 0.098 | 1.02E-05 |
| rs6534624 | -0.408 | 0.096 | 2.03E-05 |
| rs6545714 | -0.432 | 0.098 | 9.99E-06 |
| rs6548237 | -0.440 | 0.098 | 6.92E-06 |
| rs6561937 | -0.423 | 0.098 | 1.51E-05 |
| rs6575340 | -0.425 | 0.098 | 1.38E-05 |
| rs6601527 | -0.436 | 0.097 | 7.05E-06 |
| rs6658368 | -0.427 | 0.098 | 1.26E-05 |
| rs66679256 | -0.425 | 0.098 | 1.42E-05 |
| rs66833742 | -0.413 | 0.097 | 1.99E-05 |
| rs668799 | -0.398 | 0.095 | 2.89E-05 |
| rs6699744 | -0.421 | 0.098 | 1.66E-05 |
| rs6861649 | -0.420 | 0.098 | 1.65E-05 |
| rs6887247 | -0.425 | 0.098 | 1.37E-05 |
| rs6898357 | -0.423 | 0.098 | 1.50E-05 |
| rs6927268 | -0.431 | 0.098 | 1.02E-05 |
| rs7116641 | -0.429 | 0.098 | 1.18E-05 |
| rs7124681 | -0.430 | 0.098 | 1.25E-05 |
| rs7133378 | -0.430 | 0.098 | 1.11E-05 |
| rs71658797 | -0.425 | 0.098 | 1.42E-05 |
| rs7172082 | -0.431 | 0.097 | 9.66E-06 |
| rs7201895 | -0.422 | 0.098 | 1.56E-05 |
| rs7259070 | -0.424 | 0.098 | 1.41E-05 |
| rs72817602 | -0.424 | 0.098 | 1.42E-05 |
| rs73213484 | -0.423 | 0.098 | 1.49E-05 |

| | | | |
|---|---|---|---|
| rs750090 | -0.428 | 0.098 | 1.22E-05 |
| rs7516554 | -0.428 | 0.098 | 1.19E-05 |
| rs7539903 | -0.431 | 0.097 | 9.65E-06 |
| rs75499503 | -0.421 | 0.098 | 1.68E-05 |
| rs7580600 | -0.420 | 0.098 | 1.74E-05 |
| rs7591494 | -0.431 | 0.098 | 1.05E-05 |
| rs76040172 | -0.415 | 0.097 | 1.91E-05 |
| rs7613261 | -0.429 | 0.098 | 1.14E-05 |
| rs764729 | -0.434 | 0.097 | 8.15E-06 |
| rs7649970 | -0.429 | 0.098 | 1.20E-05 |
| rs76824303 | -0.427 | 0.098 | 1.27E-05 |
| rs7755574 | -0.425 | 0.098 | 1.34E-05 |
| rs7758082 | -0.433 | 0.098 | 9.94E-06 |
| rs7774 | -0.424 | 0.098 | 1.41E-05 |
| rs7915723 | -0.421 | 0.098 | 1.57E-05 |
| rs7925100 | -0.423 | 0.098 | 1.47E-05 |
| rs7930275 | -0.418 | 0.097 | 1.73E-05 |
| rs79535757 | -0.422 | 0.097 | 1.50E-05 |
| rs7972728 | -0.435 | 0.097 | 7.91E-06 |
| rs7987928 | -0.422 | 0.098 | 1.54E-05 |
| rs80135947 | -0.411 | 0.097 | 2.28E-05 |
| rs8078135 | -0.427 | 0.098 | 1.26E-05 |
| rs8087074 | -0.432 | 0.098 | 9.65E-06 |
| rs815163 | -0.430 | 0.098 | 1.05E-05 |
| rs8192675 | -0.424 | 0.098 | 1.43E-05 |
| rs845084 | -0.424 | 0.098 | 1.47E-05 |
| rs879620 | -0.413 | 0.097 | 2.25E-05 |
| rs9320823 | -0.422 | 0.098 | 1.60E-05 |
| rs9342196 | -0.421 | 0.098 | 1.58E-05 |
| rs935166 | -0.431 | 0.098 | 1.02E-05 |
| rs945211 | -0.424 | 0.098 | 1.43E-05 |
| rs9522183 | -0.429 | 0.098 | 1.16E-05 |
| rs9522279 | -0.431 | 0.098 | 9.99E-06 |
| rs9584855 | -0.431 | 0.098 | 1.01E-05 |
| rs9688977 | -0.427 | 0.098 | 1.25E-05 |
| rs972283 | -0.433 | 0.098 | 9.00E-06 |
| rs9770544 | -0.432 | 0.098 | 9.60E-06 |
| rs9843653 | -0.439 | 0.098 | 6.93E-06 |
| rs9906944 | -0.423 | 0.098 | 1.52E-05 |
| All | -0.426 | 0.097 | 1.22E-05 |

**Time spent watching television (TV) || id:UKB-b:5192**

| SNP | b | se | p |
|---|---|---|---|
| rs10109061 | -0.765 | 0.212 | 3.20E-04 |
| rs10189857 | -0.759 | 0.216 | 4.33E-04 |
| rs10269099 | -0.810 | 0.209 | 1.07E-04 |
| rs10739499 | -0.784 | 0.212 | 2.24E-04 |
| rs10765777 | -0.814 | 0.210 | 1.08E-04 |
| rs11191129 | -0.742 | 0.210 | 4.14E-04 |
| rs11222919 | -0.785 | 0.213 | 2.22E-04 |
| rs11245482 | -0.780 | 0.213 | 2.51E-04 |
| rs114755463 | -0.774 | 0.213 | 2.75E-04 |
| rs115608101 | -0.778 | 0.212 | 2.51E-04 |
| rs11662211 | -0.783 | 0.213 | 2.31E-04 |
| rs11680095 | -0.747 | 0.210 | 3.83E-04 |
| rs11696187 | -0.793 | 0.212 | 1.86E-04 |
| rs11700249 | -0.764 | 0.212 | 3.19E-04 |
| rs11714337 | -0.766 | 0.213 | 3.20E-04 |
| rs11877758 | -0.757 | 0.212 | 3.64E-04 |
| rs11911112 | -0.780 | 0.213 | 2.50E-04 |
| rs12045585 | -0.776 | 0.213 | 2.67E-04 |
| rs12214364 | -0.778 | 0.212 | 2.46E-04 |
| rs12553324 | -0.814 | 0.211 | 1.17E-04 |
| rs1291871 | -0.775 | 0.213 | 2.70E-04 |
| rs13014947 | -0.781 | 0.213 | 2.42E-04 |
| rs13107325 | -0.665 | 0.198 | 7.57E-04 |
| rs1324491 | -0.784 | 0.212 | 2.23E-04 |
| rs1727332 | -0.804 | 0.212 | 1.47E-04 |
| rs17789218 | -0.746 | 0.210 | 3.84E-04 |
| rs178203 | -0.804 | 0.211 | 1.41E-04 |
| rs180396 | -0.794 | 0.211 | 1.74E-04 |
| rs1826510 | -0.781 | 0.213 | 2.40E-04 |
| rs1993092 | -0.780 | 0.213 | 2.48E-04 |
| rs2073869 | -0.779 | 0.213 | 2.58E-04 |
| rs2106164 | -0.783 | 0.213 | 2.33E-04 |
| rs2185490 | -0.752 | 0.211 | 3.73E-04 |
| rs2240857 | -0.764 | 0.213 | 3.30E-04 |
| rs2283 | -0.764 | 0.212 | 3.23E-04 |
| rs2352984 | -0.864 | 0.211 | 4.31E-05 |
| rs2479968 | -0.781 | 0.212 | 2.28E-04 |
| rs249960 | -0.774 | 0.213 | 2.75E-04 |
| rs262890 | -0.794 | 0.213 | 1.90E-04 |
| rs263771 | -0.751 | 0.211 | 3.76E-04 |
| rs2646351 | -0.771 | 0.213 | 2.89E-04 |
| rs2678662 | -0.768 | 0.213 | 3.13E-04 |

| | | | |
|---|---|---|---|
| rs2725371 | -0.805 | 0.212 | 1.46E-04 |
| rs2785820 | -0.771 | 0.213 | 2.98E-04 |
| rs2857693 | -0.772 | 0.213 | 2.91E-04 |
| rs2906604 | -0.790 | 0.213 | 2.04E-04 |
| rs3138499 | -0.741 | 0.211 | 4.43E-04 |
| rs34811474 | -0.774 | 0.213 | 2.71E-04 |
| rs35797019 | -0.762 | 0.212 | 3.33E-04 |
| rs362312 | -0.790 | 0.212 | 1.95E-04 |
| rs3754970 | -0.761 | 0.212 | 3.43E-04 |
| rs3810496 | -0.775 | 0.213 | 2.70E-04 |
| rs4076457 | -0.764 | 0.212 | 3.19E-04 |
| rs4110177 | -0.788 | 0.212 | 2.07E-04 |
| rs4303732 | -0.782 | 0.213 | 2.40E-04 |
| rs4339469 | -0.765 | 0.213 | 3.37E-04 |
| rs4469687 | -0.758 | 0.212 | 3.47E-04 |
| rs4567133 | -0.765 | 0.213 | 3.24E-04 |
| rs4747438 | -0.752 | 0.212 | 3.88E-04 |
| rs4788616 | -0.766 | 0.213 | 3.17E-04 |
| rs4847408 | -0.836 | 0.203 | 3.93E-05 |
| rs494566 | -0.778 | 0.213 | 2.54E-04 |
| rs57555420 | -0.776 | 0.213 | 2.64E-04 |
| rs58541850 | -0.785 | 0.211 | 1.94E-04 |
| rs6102912 | -0.790 | 0.213 | 2.06E-04 |
| rs6125907 | -0.778 | 0.213 | 2.56E-04 |
| rs61743199 | -0.768 | 0.212 | 2.95E-04 |
| rs61864793 | -0.804 | 0.211 | 1.38E-04 |
| rs62145951 | -0.787 | 0.213 | 2.16E-04 |
| rs62199883 | -0.766 | 0.214 | 3.38E-04 |
| rs6511708 | -0.801 | 0.212 | 1.56E-04 |
| rs68056254 | -0.765 | 0.213 | 3.25E-04 |
| rs6814554 | -0.807 | 0.212 | 1.47E-04 |
| rs6850494 | -0.767 | 0.213 | 3.10E-04 |
| rs6994132 | -0.806 | 0.210 | 1.28E-04 |
| rs7089973 | -0.777 | 0.213 | 2.61E-04 |
| rs71658797 | -0.772 | 0.213 | 2.84E-04 |
| rs7184800 | -0.772 | 0.214 | 2.99E-04 |
| rs72673939 | -0.784 | 0.213 | 2.25E-04 |
| rs73571431 | -0.782 | 0.213 | 2.40E-04 |
| rs73946726 | -0.777 | 0.212 | 2.56E-04 |
| rs749056 | -0.779 | 0.213 | 2.50E-04 |
| rs749671 | -0.664 | 0.199 | 8.33E-04 |
| rs7539775 | -0.776 | 0.213 | 2.66E-04 |
| rs75499503 | -0.764 | 0.214 | 3.67E-04 |
| rs75641275 | -0.773 | 0.213 | 2.87E-04 |
| rs7708324 | -0.790 | 0.212 | 1.97E-04 |

| | | | |
|---|---|---|---|
| rs77273138 | -0.772 | 0.212 | 2.64E-04 |
| rs7798292 | -0.777 | 0.213 | 2.62E-04 |
| rs78227853 | -0.787 | 0.213 | 2.14E-04 |
| rs7899206 | -0.779 | 0.213 | 2.52E-04 |
| rs7921305 | -0.780 | 0.213 | 2.45E-04 |
| rs79373894 | -0.772 | 0.212 | 2.74E-04 |
| rs801733 | -0.725 | 0.209 | 5.34E-04 |
| rs814197 | -0.781 | 0.213 | 2.46E-04 |
| rs872169 | -0.768 | 0.213 | 3.05E-04 |
| rs898751 | -0.759 | 0.212 | 3.52E-04 |
| rs9300594 | -0.773 | 0.213 | 2.82E-04 |
| rs9471333 | -0.777 | 0.213 | 2.66E-04 |
| rs9834970 | -0.753 | 0.213 | 3.95E-04 |
| rs9867121 | -0.763 | 0.212 | 3.27E-04 |
| rs9867437 | -0.752 | 0.212 | 3.85E-04 |
| rs9880023 | -0.747 | 0.211 | 3.95E-04 |
| rs996234 | -0.794 | 0.212 | 1.86E-04 |
| All | -0.775 | 0.211 | 2.40E-04 |

**Trunk fat mass || id:UKB-a:291**

| SNP | b | se | p |
|---|---|---|---|
| rs10100245 | -0.2485 | 0.0699 | 3.79E-04 |
| rs1013293 | -0.2403 | 0.0696 | 5.54E-04 |
| rs10172196 | -0.2495 | 0.0699 | 3.55E-04 |
| rs1017529 | -0.2470 | 0.0698 | 4.04E-04 |
| rs10237306 | -0.2511 | 0.0697 | 3.17E-04 |
| rs10264581 | -0.2510 | 0.0697 | 3.16E-04 |
| rs10269774 | -0.2413 | 0.0697 | 5.31E-04 |
| rs10423928 | -0.2503 | 0.0699 | 3.47E-04 |
| rs10499014 | -0.2473 | 0.0698 | 3.98E-04 |
| rs1056441 | -0.2485 | 0.0698 | 3.74E-04 |
| rs10756714 | -0.2401 | 0.0697 | 5.72E-04 |
| rs10756798 | -0.2531 | 0.0696 | 2.78E-04 |
| rs10760724 | -0.2474 | 0.0698 | 3.95E-04 |
| rs10830566 | -0.2488 | 0.0698 | 3.64E-04 |
| rs10863753 | -0.2512 | 0.0697 | 3.12E-04 |
| rs10915846 | -0.2510 | 0.0697 | 3.15E-04 |
| rs10938398 | -0.2529 | 0.0699 | 2.98E-04 |
| rs10999460 | -0.2426 | 0.0696 | 4.93E-04 |
| rs11012732 | -0.2343 | 0.0696 | 7.56E-04 |
| rs11023199 | -0.2446 | 0.0697 | 4.50E-04 |
| rs11030119 | -0.2423 | 0.0700 | 5.33E-04 |
| rs11042030 | -0.2480 | 0.0699 | 3.86E-04 |
| rs11043764 | -0.2447 | 0.0697 | 4.48E-04 |
| rs11135450 | -0.2546 | 0.0693 | 2.40E-04 |
| rs11150745 | -0.2460 | 0.0698 | 4.29E-04 |
| rs11205303 | -0.2610 | 0.0700 | 1.92E-04 |
| rs112646560 | -0.2467 | 0.0698 | 4.11E-04 |
| rs11538 | -0.2496 | 0.0698 | 3.45E-04 |
| rs11603783 | -0.2519 | 0.0697 | 2.99E-04 |
| rs11642015 | -0.2427 | 0.0711 | 6.41E-04 |
| rs11742930 | -0.2476 | 0.0698 | 3.93E-04 |
| rs11779446 | -0.2558 | 0.0692 | 2.21E-04 |
| rs11782074 | -0.2502 | 0.0698 | 3.35E-04 |
| rs11782341 | -0.2524 | 0.0696 | 2.86E-04 |
| rs1182199 | -0.2545 | 0.0699 | 2.69E-04 |
| rs11856579 | -0.2449 | 0.0697 | 4.46E-04 |
| rs11917068 | -0.2484 | 0.0698 | 3.72E-04 |
| rs12103006 | -0.2477 | 0.0699 | 3.95E-04 |
| rs12144626 | -0.2524 | 0.0696 | 2.86E-04 |
| rs1218824 | -0.2484 | 0.0698 | 3.76E-04 |
| rs12213070 | -0.2475 | 0.0698 | 3.94E-04 |
| rs12330631 | -0.2494 | 0.0698 | 3.52E-04 |

| | | | |
|---|---|---|---|
| rs12339822 | -0.2432 | 0.0696 | 4.78E-04 |
| rs12367809 | -0.2523 | 0.0700 | 3.12E-04 |
| rs12459368 | -0.2468 | 0.0698 | 4.08E-04 |
| rs12475388 | -0.2494 | 0.0698 | 3.54E-04 |
| rs12578952 | -0.2444 | 0.0697 | 4.54E-04 |
| rs12616638 | -0.2513 | 0.0697 | 3.10E-04 |
| rs12619178 | -0.2483 | 0.0698 | 3.78E-04 |
| rs12679106 | -0.2505 | 0.0699 | 3.40E-04 |
| rs1268403 | -0.2486 | 0.0698 | 3.70E-04 |
| rs12724928 | -0.2461 | 0.0698 | 4.26E-04 |
| rs12877270 | -0.2452 | 0.0698 | 4.39E-04 |
| rs12890931 | -0.2457 | 0.0698 | 4.30E-04 |
| rs12901071 | -0.2459 | 0.0698 | 4.29E-04 |
| rs12926311 | -0.2438 | 0.0697 | 4.69E-04 |
| rs12974458 | -0.2440 | 0.0696 | 4.58E-04 |
| rs12992672 | -0.2571 | 0.0699 | 2.36E-04 |
| rs13067187 | -0.2462 | 0.0698 | 4.21E-04 |
| rs13135092 | -0.2364 | 0.0693 | 6.49E-04 |
| rs13156484 | -0.2466 | 0.0698 | 4.13E-04 |
| rs13191298 | -0.2477 | 0.0698 | 3.88E-04 |
| rs13333747 | -0.2470 | 0.0698 | 4.03E-04 |
| rs13340461 | -0.2476 | 0.0699 | 3.93E-04 |
| rs1336486 | -0.2463 | 0.0698 | 4.19E-04 |
| rs13416004 | -0.2330 | 0.0689 | 7.13E-04 |
| rs1427325 | -0.2476 | 0.0699 | 3.94E-04 |
| rs143384 | -0.2520 | 0.0699 | 3.11E-04 |
| rs1441264 | -0.2455 | 0.0698 | 4.38E-04 |
| rs1446585 | -0.2694 | 0.0671 | 5.95E-05 |
| rs1470050 | -0.2512 | 0.0697 | 3.11E-04 |
| rs1477290 | -0.2528 | 0.0698 | 2.93E-04 |
| rs147730268 | -0.2550 | 0.0697 | 2.55E-04 |
| rs1491905 | -0.2398 | 0.0690 | 5.12E-04 |
| rs1503527 | -0.2489 | 0.0698 | 3.65E-04 |
| rs154001 | -0.2471 | 0.0699 | 4.07E-04 |
| rs1568489 | -0.2531 | 0.0696 | 2.76E-04 |
| rs1627536 | -0.2451 | 0.0697 | 4.41E-04 |
| rs1652376 | -0.2414 | 0.0698 | 5.40E-04 |
| rs1653892 | -0.2503 | 0.0698 | 3.35E-04 |
| rs16880854 | -0.2526 | 0.0699 | 3.02E-04 |
| rs16934748 | -0.2501 | 0.0697 | 3.34E-04 |
| rs16951304 | -0.2480 | 0.0699 | 3.91E-04 |
| rs16967461 | -0.2492 | 0.0698 | 3.57E-04 |
| rs17024393 | -0.2480 | 0.0699 | 3.90E-04 |
| rs17055384 | -0.2433 | 0.0696 | 4.72E-04 |
| rs17218879 | -0.2448 | 0.0697 | 4.47E-04 |

| | | | |
|---|---|---|---|
| rs1724551 | -0.2471 | 0.0698 | 4.02E-04 |
| rs17265513 | -0.2539 | 0.0696 | 2.64E-04 |
| rs17522122 | -0.2502 | 0.0698 | 3.38E-04 |
| rs17698176 | -0.2561 | 0.0686 | 1.88E-04 |
| rs17770336 | -0.2435 | 0.0698 | 4.88E-04 |
| rs1782508 | -0.2478 | 0.0698 | 3.87E-04 |
| rs1834144 | -0.2372 | 0.0692 | 6.08E-04 |
| rs1927626 | -0.2454 | 0.0698 | 4.38E-04 |
| rs1964599 | -0.2479 | 0.0699 | 3.88E-04 |
| rs2002023 | -0.2475 | 0.0698 | 3.94E-04 |
| rs2012027 | -0.2480 | 0.0699 | 3.86E-04 |
| rs2016469 | -0.2443 | 0.0696 | 4.52E-04 |
| rs2102278 | -0.2431 | 0.0696 | 4.79E-04 |
| rs2172131 | -0.2438 | 0.0697 | 4.70E-04 |
| rs217667 | -0.2497 | 0.0698 | 3.45E-04 |
| rs2216931 | -0.2471 | 0.0699 | 4.05E-04 |
| rs2243928 | -0.2504 | 0.0698 | 3.31E-04 |
| rs2287214 | -0.2490 | 0.0699 | 3.67E-04 |
| rs2289379 | -0.2483 | 0.0698 | 3.77E-04 |
| rs2306593 | -0.2527 | 0.0696 | 2.81E-04 |
| rs2307111 | -0.2461 | 0.0700 | 4.36E-04 |
| rs2370982 | -0.2485 | 0.0700 | 3.82E-04 |
| rs2371767 | -0.2419 | 0.0695 | 5.00E-04 |
| rs2374947 | -0.2514 | 0.0696 | 3.06E-04 |
| rs241460 | -0.2485 | 0.0699 | 3.78E-04 |
| rs2439823 | -0.2487 | 0.0699 | 3.71E-04 |
| rs245775 | -0.2498 | 0.0698 | 3.47E-04 |
| rs2482704 | -0.2500 | 0.0698 | 3.39E-04 |
| rs2494196 | -0.2472 | 0.0700 | 4.12E-04 |
| rs2500406 | -0.2385 | 0.0692 | 5.64E-04 |
| rs252749 | -0.2417 | 0.0697 | 5.20E-04 |
| rs2678204 | -0.2410 | 0.0697 | 5.46E-04 |
| rs274677 | -0.2441 | 0.0697 | 4.59E-04 |
| rs28366156 | -0.2514 | 0.0698 | 3.13E-04 |
| rs2855818 | -0.2389 | 0.0694 | 5.75E-04 |
| rs28636067 | -0.2466 | 0.0698 | 4.12E-04 |
| rs2869950 | -0.2385 | 0.0695 | 6.00E-04 |
| rs287834 | -0.2532 | 0.0694 | 2.66E-04 |
| rs2903738 | -0.2509 | 0.0697 | 3.18E-04 |
| rs2914231 | -0.2519 | 0.0697 | 2.99E-04 |
| rs2954021 | -0.2475 | 0.0699 | 3.96E-04 |
| rs2979655 | -0.2475 | 0.0698 | 3.94E-04 |
| rs314279 | -0.2441 | 0.0698 | 4.69E-04 |
| rs33503 | -0.2491 | 0.0698 | 3.57E-04 |
| rs33836 | -0.2458 | 0.0699 | 4.35E-04 |

| | | | |
|---|---|---|---|
| rs34045288 | -0.2492 | 0.0699 | 3.65E-04 |
| rs34517439 | -0.2457 | 0.0697 | 4.24E-04 |
| rs34560402 | -0.2459 | 0.0698 | 4.25E-04 |
| rs35060985 | -0.2492 | 0.0699 | 3.65E-04 |
| rs35344761 | -0.2485 | 0.0698 | 3.72E-04 |
| rs35537311 | -0.2484 | 0.0698 | 3.72E-04 |
| rs3730071 | -0.2511 | 0.0698 | 3.19E-04 |
| rs3803286 | -0.2516 | 0.0698 | 3.11E-04 |
| rs3806251 | -0.2482 | 0.0699 | 3.81E-04 |
| rs3814883 | -0.2378 | 0.0697 | 6.42E-04 |
| rs3817428 | -0.2546 | 0.0699 | 2.70E-04 |
| rs3853252 | -0.2508 | 0.0697 | 3.24E-04 |
| rs40071 | -0.2470 | 0.0698 | 4.05E-04 |
| rs4253755 | -0.2528 | 0.0695 | 2.76E-04 |
| rs429358 | -0.2503 | 0.0698 | 3.38E-04 |
| rs4477562 | -0.2469 | 0.0699 | 4.09E-04 |
| rs4549080 | -0.2492 | 0.0698 | 3.57E-04 |
| rs4657796 | -0.2459 | 0.0698 | 4.27E-04 |
| rs4673617 | -0.2500 | 0.0697 | 3.37E-04 |
| rs4709745 | -0.2472 | 0.0698 | 4.00E-04 |
| rs4718964 | -0.2473 | 0.0699 | 4.03E-04 |
| rs4722398 | -0.2554 | 0.0692 | 2.25E-04 |
| rs4757144 | -0.2514 | 0.0697 | 3.08E-04 |
| rs477895 | -0.2468 | 0.0698 | 4.09E-04 |
| rs4790841 | -0.2485 | 0.0699 | 3.79E-04 |
| rs4804312 | -0.2485 | 0.0698 | 3.71E-04 |
| rs4821764 | -0.2516 | 0.0698 | 3.13E-04 |
| rs4864201 | -0.2481 | 0.0698 | 3.81E-04 |
| rs4876611 | -0.2444 | 0.0699 | 4.68E-04 |
| rs4900471 | -0.2501 | 0.0697 | 3.34E-04 |
| rs4958702 | -0.2469 | 0.0698 | 4.07E-04 |
| rs4981693 | -0.2487 | 0.0699 | 3.72E-04 |
| rs543874 | -0.2624 | 0.0702 | 1.87E-04 |
| rs55637757 | -0.2457 | 0.0698 | 4.27E-04 |
| rs55672437 | -0.2403 | 0.0693 | 5.23E-04 |
| rs55714539 | -0.2475 | 0.0699 | 3.99E-04 |
| rs55726687 | -0.2567 | 0.0691 | 2.02E-04 |
| rs56239516 | -0.2494 | 0.0697 | 3.47E-04 |
| rs56254492 | -0.2492 | 0.0698 | 3.58E-04 |
| rs56288810 | -0.2513 | 0.0697 | 3.11E-04 |
| rs56361700 | -0.2443 | 0.0697 | 4.57E-04 |
| rs57636386 | -0.2457 | 0.0699 | 4.42E-04 |
| rs59066241 | -0.2490 | 0.0698 | 3.61E-04 |
| rs59738707 | -0.2472 | 0.0698 | 3.99E-04 |
| rs59815219 | -0.2450 | 0.0697 | 4.43E-04 |

| | | | |
|---|---|---|---|
| rs6029180 | -0.2454 | 0.0698 | 4.35E-04 |
| rs60898782 | -0.2506 | 0.0697 | 3.27E-04 |
| rs6135562 | -0.2510 | 0.0697 | 3.15E-04 |
| rs61813293 | -0.2468 | 0.0697 | 4.01E-04 |
| rs62025831 | -0.2479 | 0.0700 | 4.00E-04 |
| rs62037365 | -0.2347 | 0.0698 | 7.72E-04 |
| rs62104483 | -0.2404 | 0.0697 | 5.61E-04 |
| rs62190394 | -0.2432 | 0.0698 | 4.93E-04 |
| rs62246314 | -0.2472 | 0.0698 | 3.97E-04 |
| rs62261725 | -0.2419 | 0.0697 | 5.16E-04 |
| rs62396185 | -0.2315 | 0.0700 | 9.34E-04 |
| rs62407569 | -0.2517 | 0.0696 | 3.01E-04 |
| rs6486057 | -0.2476 | 0.0698 | 3.91E-04 |
| rs6545714 | -0.2512 | 0.0699 | 3.22E-04 |
| rs6549970 | -0.2472 | 0.0698 | 3.99E-04 |
| rs6567160 | -0.2405 | 0.0702 | 6.17E-04 |
| rs6575340 | -0.2470 | 0.0699 | 4.09E-04 |
| rs6598540 | -0.2446 | 0.0698 | 4.54E-04 |
| rs66679256 | -0.2465 | 0.0699 | 4.20E-04 |
| rs6696828 | -0.2529 | 0.0695 | 2.73E-04 |
| rs6699744 | -0.2440 | 0.0698 | 4.76E-04 |
| rs6717858 | -0.2447 | 0.0699 | 4.63E-04 |
| rs67536595 | -0.2475 | 0.0698 | 3.93E-04 |
| rs67913249 | -0.2504 | 0.0697 | 3.27E-04 |
| rs6861649 | -0.2436 | 0.0697 | 4.75E-04 |
| rs6948959 | -0.2468 | 0.0698 | 4.10E-04 |
| rs6973656 | -0.2507 | 0.0698 | 3.26E-04 |
| rs704061 | -0.2543 | 0.0696 | 2.57E-04 |
| rs7080838 | -0.2443 | 0.0697 | 4.59E-04 |
| rs7094644 | -0.2538 | 0.0694 | 2.57E-04 |
| rs7124681 | -0.2481 | 0.0700 | 3.93E-04 |
| rs713586 | -0.2448 | 0.0701 | 4.83E-04 |
| rs7167767 | -0.2452 | 0.0697 | 4.40E-04 |
| rs7171864 | -0.2526 | 0.0697 | 2.89E-04 |
| rs7193783 | -0.2476 | 0.0699 | 3.95E-04 |
| rs7218014 | -0.2357 | 0.0695 | 6.97E-04 |
| rs7238896 | -0.2440 | 0.0697 | 4.61E-04 |
| rs724016 | -0.2502 | 0.0703 | 3.75E-04 |
| rs7259070 | -0.2463 | 0.0698 | 4.15E-04 |
| rs72718281 | -0.2491 | 0.0699 | 3.63E-04 |
| rs72755233 | -0.2522 | 0.0695 | 2.87E-04 |
| rs72820274 | -0.2475 | 0.0698 | 3.92E-04 |
| rs72917533 | -0.2476 | 0.0698 | 3.91E-04 |
| rs72976986 | -0.2404 | 0.0692 | 5.12E-04 |
| rs72995085 | -0.2473 | 0.0698 | 3.97E-04 |

| | | | |
|---|---|---|---|
| rs73041988 | -0.2475 | 0.0699 | 3.96E-04 |
| rs7305790 | -0.2442 | 0.0697 | 4.60E-04 |
| rs7313748 | -0.2514 | 0.0697 | 3.08E-04 |
| rs73142879 | -0.2457 | 0.0698 | 4.35E-04 |
| rs73175572 | -0.2441 | 0.0696 | 4.54E-04 |
| rs73213484 | -0.2454 | 0.0698 | 4.38E-04 |
| rs7460093 | -0.2447 | 0.0696 | 4.42E-04 |
| rs750090 | -0.2484 | 0.0698 | 3.74E-04 |
| rs75412871 | -0.2467 | 0.0698 | 4.11E-04 |
| rs75641275 | -0.2466 | 0.0698 | 4.12E-04 |
| rs756717 | -0.2474 | 0.0698 | 3.95E-04 |
| rs7600835 | -0.2467 | 0.0698 | 4.09E-04 |
| rs76040172 | -0.2408 | 0.0695 | 5.27E-04 |
| rs7613261 | -0.2494 | 0.0699 | 3.57E-04 |
| rs76345589 | -0.2496 | 0.0698 | 3.52E-04 |
| rs7649970 | -0.2490 | 0.0699 | 3.70E-04 |
| rs768023 | -0.2534 | 0.0697 | 2.78E-04 |
| rs7696324 | -0.2471 | 0.0699 | 4.03E-04 |
| rs7697556 | -0.2493 | 0.0699 | 3.59E-04 |
| rs7933085 | -0.2513 | 0.0697 | 3.12E-04 |
| rs79535757 | -0.2450 | 0.0696 | 4.35E-04 |
| rs7987928 | -0.2449 | 0.0698 | 4.49E-04 |
| rs8087074 | -0.2506 | 0.0697 | 3.25E-04 |
| rs815163 | -0.2499 | 0.0698 | 3.42E-04 |
| rs8178882 | -0.2481 | 0.0698 | 3.80E-04 |
| rs8192675 | -0.2463 | 0.0698 | 4.20E-04 |
| rs846545 | -0.2476 | 0.0698 | 3.90E-04 |
| rs854917 | -0.2445 | 0.0697 | 4.54E-04 |
| rs879620 | -0.2383 | 0.0697 | 6.33E-04 |
| rs882378 | -0.2444 | 0.0697 | 4.52E-04 |
| rs9320823 | -0.2446 | 0.0699 | 4.61E-04 |
| rs9375702 | -0.2458 | 0.0698 | 4.30E-04 |
| rs952227 | -0.2473 | 0.0698 | 3.98E-04 |
| rs9584855 | -0.2502 | 0.0698 | 3.36E-04 |
| rs962554 | -0.2466 | 0.0698 | 4.16E-04 |
| rs9814633 | -0.2508 | 0.0697 | 3.22E-04 |
| rs9843653 | -0.2552 | 0.0698 | 2.55E-04 |
| rs9861443 | -0.2515 | 0.0695 | 2.95E-04 |
| All | -0.2476 | 0.0696 | 3.76E-04 |

**Arm fat mass (left) || id:UKB-a:287**

| SNP | b | se | p |
|---|---|---|---|
| rs10100245 | -0.2847 | 0.0742 | 1.26E-04 |
| rs1011326 | -0.2847 | 0.0741 | 1.23E-04 |
| rs10132436 | -0.2836 | 0.0741 | 1.30E-04 |
| rs1013293 | -0.2758 | 0.0739 | 1.90E-04 |
| rs10187101 | -0.2867 | 0.0740 | 1.08E-04 |
| rs10237317 | -0.2828 | 0.0741 | 1.36E-04 |
| rs10259620 | -0.2887 | 0.0738 | 9.24E-05 |
| rs10269774 | -0.2782 | 0.0739 | 1.66E-04 |
| rs10404726 | -0.2842 | 0.0742 | 1.28E-04 |
| rs10756798 | -0.2894 | 0.0739 | 9.02E-05 |
| rs10760724 | -0.2834 | 0.0742 | 1.32E-04 |
| rs10787738 | -0.2870 | 0.0740 | 1.05E-04 |
| rs10820852 | -0.2846 | 0.0741 | 1.23E-04 |
| rs10938398 | -0.2902 | 0.0743 | 9.33E-05 |
| rs10947793 | -0.2830 | 0.0741 | 1.34E-04 |
| rs10954772 | -0.2870 | 0.0740 | 1.06E-04 |
| rs11001963 | -0.2820 | 0.0741 | 1.42E-04 |
| rs11012732 | -0.2698 | 0.0739 | 2.60E-04 |
| rs11030119 | -0.2779 | 0.0744 | 1.86E-04 |
| rs11042030 | -0.2840 | 0.0742 | 1.29E-04 |
| rs11099020 | -0.2849 | 0.0741 | 1.22E-04 |
| rs11135450 | -0.2904 | 0.0736 | 7.93E-05 |
| rs11150745 | -0.2818 | 0.0742 | 1.45E-04 |
| rs11152135 | -0.2837 | 0.0741 | 1.30E-04 |
| rs11165643 | -0.2809 | 0.0742 | 1.53E-04 |
| rs11205617 | -0.2842 | 0.0742 | 1.28E-04 |
| rs112094960 | -0.2759 | 0.0734 | 1.70E-04 |
| rs1127100 | -0.2819 | 0.0741 | 1.43E-04 |
| rs113182412 | -0.2851 | 0.0741 | 1.18E-04 |
| rs113230003 | -0.2829 | 0.0742 | 1.36E-04 |
| rs113603865 | -0.2828 | 0.0742 | 1.38E-04 |
| rs113866544 | -0.2872 | 0.0741 | 1.06E-04 |
| rs11515071 | -0.2770 | 0.0741 | 1.86E-04 |
| rs11538 | -0.2857 | 0.0741 | 1.14E-04 |
| rs11642015 | -0.2807 | 0.0760 | 2.21E-04 |
| rs11742930 | -0.2835 | 0.0741 | 1.31E-04 |
| rs11782341 | -0.2887 | 0.0739 | 9.30E-05 |
| rs11786089 | -0.2815 | 0.0741 | 1.44E-04 |
| rs1182143 | -0.2893 | 0.0739 | 9.10E-05 |
| rs11882409 | -0.2831 | 0.0742 | 1.35E-04 |
| rs11920781 | -0.2857 | 0.0741 | 1.16E-04 |
| rs12001437 | -0.2786 | 0.0738 | 1.61E-04 |

| | | | |
|---|---|---|---|
| rs12037698 | -0.2813 | 0.0741 | 1.46E-04 |
| rs12072739 | -0.2790 | 0.0739 | 1.61E-04 |
| rs12103006 | -0.2834 | 0.0741 | 1.32E-04 |
| rs12144626 | -0.2897 | 0.0739 | 8.79E-05 |
| rs1218824 | -0.2844 | 0.0742 | 1.25E-04 |
| rs12254441 | -0.2795 | 0.0740 | 1.58E-04 |
| rs12339822 | -0.2787 | 0.0740 | 1.64E-04 |
| rs12367809 | -0.2888 | 0.0743 | 1.01E-04 |
| rs12506689 | -0.2842 | 0.0742 | 1.27E-04 |
| rs12578952 | -0.2798 | 0.0740 | 1.57E-04 |
| rs12619178 | -0.2844 | 0.0742 | 1.26E-04 |
| rs12619626 | -0.2875 | 0.0739 | 1.00E-04 |
| rs1263627 | -0.2888 | 0.0738 | 9.06E-05 |
| rs1268403 | -0.2844 | 0.0741 | 1.24E-04 |
| rs12692738 | -0.2847 | 0.0742 | 1.23E-04 |
| rs1286138 | -0.2845 | 0.0741 | 1.24E-04 |
| rs12877270 | -0.2808 | 0.0741 | 1.50E-04 |
| rs12885458 | -0.2744 | 0.0733 | 1.83E-04 |
| rs12887636 | -0.2850 | 0.0741 | 1.20E-04 |
| rs1296328 | -0.2822 | 0.0741 | 1.42E-04 |
| rs12992672 | -0.2959 | 0.0743 | 6.89E-05 |
| rs13062093 | -0.2839 | 0.0742 | 1.29E-04 |
| rs13076052 | -0.2863 | 0.0740 | 1.09E-04 |
| rs13135092 | -0.2717 | 0.0736 | 2.23E-04 |
| rs13174863 | -0.2816 | 0.0742 | 1.46E-04 |
| rs1335055 | -0.2816 | 0.0741 | 1.45E-04 |
| rs13427822 | -0.2821 | 0.0741 | 1.41E-04 |
| rs1406778 | -0.2859 | 0.0742 | 1.16E-04 |
| rs1441264 | -0.2812 | 0.0742 | 1.50E-04 |
| rs1446585 | -0.3058 | 0.0712 | 1.75E-05 |
| rs1459189 | -0.2819 | 0.0741 | 1.42E-04 |
| rs1470749 | -0.2858 | 0.0741 | 1.14E-04 |
| rs1471338 | -0.2853 | 0.0741 | 1.17E-04 |
| rs1477290 | -0.2893 | 0.0741 | 9.49E-05 |
| rs147730268 | -0.2906 | 0.0740 | 8.58E-05 |
| rs1503526 | -0.2841 | 0.0742 | 1.28E-04 |
| rs1582931 | -0.2824 | 0.0742 | 1.41E-04 |
| rs1609303 | -0.2837 | 0.0742 | 1.31E-04 |
| rs1652376 | -0.2775 | 0.0740 | 1.79E-04 |
| rs17024393 | -0.2843 | 0.0743 | 1.29E-04 |
| rs17311369 | -0.2869 | 0.0740 | 1.06E-04 |
| rs17627049 | -0.2880 | 0.0739 | 9.82E-05 |
| rs17716502 | -0.2801 | 0.0741 | 1.59E-04 |
| rs17770336 | -0.2790 | 0.0742 | 1.69E-04 |
| rs1782508 | -0.2838 | 0.0741 | 1.29E-04 |

| | | | |
|---|---|---|---|
| rs1783541 | -0.2877 | 0.0739 | 9.95E-05 |
| rs1840126 | -0.2775 | 0.0737 | 1.65E-04 |
| rs1881505 | -0.2828 | 0.0740 | 1.32E-04 |
| rs1899689 | -0.2869 | 0.0740 | 1.06E-04 |
| rs1928496 | -0.2827 | 0.0741 | 1.37E-04 |
| rs1964599 | -0.2836 | 0.0741 | 1.30E-04 |
| rs2016469 | -0.2799 | 0.0739 | 1.53E-04 |
| rs2052607 | -0.2743 | 0.0736 | 1.93E-04 |
| rs2098870 | -0.2843 | 0.0741 | 1.25E-04 |
| rs2102278 | -0.2790 | 0.0739 | 1.59E-04 |
| rs2172131 | -0.2794 | 0.0740 | 1.61E-04 |
| rs217672 | -0.2859 | 0.0741 | 1.13E-04 |
| rs2239647 | -0.2868 | 0.0742 | 1.10E-04 |
| rs2253310 | -0.2897 | 0.0740 | 8.99E-05 |
| rs2287214 | -0.2846 | 0.0742 | 1.24E-04 |
| rs2289379 | -0.2845 | 0.0742 | 1.25E-04 |
| rs2292238 | -0.2846 | 0.0741 | 1.23E-04 |
| rs2306593 | -0.2899 | 0.0739 | 8.75E-05 |
| rs2307111 | -0.2821 | 0.0743 | 1.47E-04 |
| rs2318543 | -0.2841 | 0.0741 | 1.27E-04 |
| rs2370982 | -0.2845 | 0.0743 | 1.27E-04 |
| rs2374947 | -0.2879 | 0.0739 | 9.84E-05 |
| rs2425024 | -0.2847 | 0.0741 | 1.23E-04 |
| rs2439823 | -0.2850 | 0.0742 | 1.22E-04 |
| rs2444253 | -0.2797 | 0.0739 | 1.55E-04 |
| rs245775 | -0.2861 | 0.0741 | 1.14E-04 |
| rs2470619 | -0.2865 | 0.0740 | 1.08E-04 |
| rs2479958 | -0.2845 | 0.0741 | 1.24E-04 |
| rs2494196 | -0.2829 | 0.0742 | 1.36E-04 |
| rs2499468 | -0.2843 | 0.0741 | 1.26E-04 |
| rs252749 | -0.2778 | 0.0739 | 1.71E-04 |
| rs2678204 | -0.2760 | 0.0741 | 1.95E-04 |
| rs273505 | -0.2842 | 0.0742 | 1.28E-04 |
| rs2814943 | -0.2946 | 0.0743 | 7.30E-05 |
| rs28366156 | -0.2875 | 0.0740 | 1.03E-04 |
| rs28414958 | -0.2819 | 0.0741 | 1.42E-04 |
| rs28489620 | -0.2763 | 0.0736 | 1.73E-04 |
| rs2861685 | -0.2799 | 0.0741 | 1.57E-04 |
| rs286818 | -0.2832 | 0.0742 | 1.35E-04 |
| rs3134404 | -0.2888 | 0.0738 | 9.14E-05 |
| rs3211995 | -0.2832 | 0.0741 | 1.33E-04 |
| rs34045288 | -0.2856 | 0.0743 | 1.20E-04 |
| rs34045894 | -0.2870 | 0.0740 | 1.05E-04 |
| rs34292685 | -0.2826 | 0.0741 | 1.38E-04 |
| rs34517439 | -0.2815 | 0.0740 | 1.43E-04 |

| | | | |
|---|---|---|---|
| rs34769775 | -0.2824 | 0.0741 | 1.40E-04 |
| rs34811474 | -0.2829 | 0.0741 | 1.35E-04 |
| rs35537311 | -0.2844 | 0.0741 | 1.24E-04 |
| rs35851183 | -0.2795 | 0.0740 | 1.59E-04 |
| rs35882248 | -0.2788 | 0.0741 | 1.67E-04 |
| rs362307 | -0.2846 | 0.0741 | 1.22E-04 |
| rs3729793 | -0.2799 | 0.0739 | 1.53E-04 |
| rs3770821 | -0.2852 | 0.0741 | 1.19E-04 |
| rs3802858 | -0.2851 | 0.0741 | 1.19E-04 |
| rs3803286 | -0.2879 | 0.0741 | 1.01E-04 |
| rs3810291 | -0.2783 | 0.0744 | 1.83E-04 |
| rs3826408 | -0.2912 | 0.0739 | 8.05E-05 |
| rs390192 | -0.2900 | 0.0737 | 8.35E-05 |
| rs4072917 | -0.2805 | 0.0740 | 1.51E-04 |
| rs4073582 | -0.2738 | 0.0733 | 1.88E-04 |
| rs4246657 | -0.2855 | 0.0741 | 1.17E-04 |
| rs4402589 | -0.2727 | 0.0741 | 2.35E-04 |
| rs4477562 | -0.2829 | 0.0742 | 1.37E-04 |
| rs4502882 | -0.2837 | 0.0742 | 1.30E-04 |
| rs4503172 | -0.2855 | 0.0741 | 1.16E-04 |
| rs4648450 | -0.2834 | 0.0741 | 1.31E-04 |
| rs4657796 | -0.2817 | 0.0741 | 1.44E-04 |
| rs4673617 | -0.2867 | 0.0741 | 1.08E-04 |
| rs4725984 | -0.2802 | 0.0741 | 1.55E-04 |
| rs4740383 | -0.2746 | 0.0731 | 1.74E-04 |
| rs4757142 | -0.2891 | 0.0738 | 9.00E-05 |
| rs4776970 | -0.2826 | 0.0745 | 1.48E-04 |
| rs4777541 | -0.2853 | 0.0741 | 1.19E-04 |
| rs4790841 | -0.2846 | 0.0742 | 1.26E-04 |
| rs4886039 | -0.2806 | 0.0741 | 1.52E-04 |
| rs4921301 | -0.2829 | 0.0741 | 1.35E-04 |
| rs4958702 | -0.2828 | 0.0742 | 1.37E-04 |
| rs4962424 | -0.2866 | 0.0740 | 1.08E-04 |
| rs541577 | -0.2863 | 0.0740 | 1.11E-04 |
| rs543874 | -0.3027 | 0.0747 | 5.10E-05 |
| rs55637757 | -0.2818 | 0.0740 | 1.41E-04 |
| rs55655049 | -0.2803 | 0.0740 | 1.52E-04 |
| rs55726687 | -0.2944 | 0.0733 | 5.93E-05 |
| rs55966114 | -0.2805 | 0.0740 | 1.52E-04 |
| rs56288810 | -0.2876 | 0.0740 | 1.01E-04 |
| rs56803094 | -0.2822 | 0.0741 | 1.40E-04 |
| rs57636386 | -0.2817 | 0.0743 | 1.50E-04 |
| rs58084604 | -0.2751 | 0.0748 | 2.36E-04 |
| rs6029180 | -0.2813 | 0.0741 | 1.46E-04 |
| rs6050446 | -0.2836 | 0.0742 | 1.33E-04 |

| | | | |
|---|---|---|---|
| rs60764613 | -0.2800 | 0.0740 | 1.55E-04 |
| rs61649432 | -0.2796 | 0.0739 | 1.55E-04 |
| rs61813324 | -0.2812 | 0.0740 | 1.45E-04 |
| rs62037365 | -0.2709 | 0.0741 | 2.56E-04 |
| rs62104483 | -0.2759 | 0.0740 | 1.93E-04 |
| rs62246314 | -0.2831 | 0.0741 | 1.33E-04 |
| rs62261725 | -0.2772 | 0.0740 | 1.80E-04 |
| rs62396185 | -0.2710 | 0.0739 | 2.45E-04 |
| rs62477684 | -0.2863 | 0.0742 | 1.13E-04 |
| rs6445198 | -0.2836 | 0.0741 | 1.31E-04 |
| rs649648 | -0.2863 | 0.0740 | 1.10E-04 |
| rs6575340 | -0.2830 | 0.0742 | 1.38E-04 |
| rs6601527 | -0.2906 | 0.0738 | 8.24E-05 |
| rs6656785 | -0.2816 | 0.0741 | 1.45E-04 |
| rs66679256 | -0.2825 | 0.0742 | 1.41E-04 |
| rs668799 | -0.2662 | 0.0726 | 2.45E-04 |
| rs6699744 | -0.2796 | 0.0742 | 1.63E-04 |
| rs6739755 | -0.2867 | 0.0742 | 1.11E-04 |
| rs67602344 | -0.2760 | 0.0735 | 1.74E-04 |
| rs67807996 | -0.2878 | 0.0738 | 9.72E-05 |
| rs67844506 | -0.2817 | 0.0742 | 1.46E-04 |
| rs6802087 | -0.2842 | 0.0741 | 1.26E-04 |
| rs6861649 | -0.2796 | 0.0740 | 1.57E-04 |
| rs705145 | -0.2784 | 0.0739 | 1.64E-04 |
| rs7094644 | -0.2898 | 0.0737 | 8.42E-05 |
| rs7116641 | -0.2855 | 0.0743 | 1.21E-04 |
| rs7124681 | -0.2844 | 0.0743 | 1.31E-04 |
| rs713586 | -0.2814 | 0.0748 | 1.68E-04 |
| rs7149242 | -0.2863 | 0.0740 | 1.10E-04 |
| rs7201895 | -0.2805 | 0.0741 | 1.53E-04 |
| rs7218014 | -0.2730 | 0.0737 | 2.12E-04 |
| rs72917533 | -0.2836 | 0.0741 | 1.31E-04 |
| rs73213484 | -0.2812 | 0.0741 | 1.48E-04 |
| rs745249 | -0.2849 | 0.0741 | 1.21E-04 |
| rs7498044 | -0.2840 | 0.0741 | 1.27E-04 |
| rs750090 | -0.2845 | 0.0741 | 1.24E-04 |
| rs756717 | -0.2834 | 0.0741 | 1.32E-04 |
| rs76040172 | -0.2759 | 0.0738 | 1.84E-04 |
| rs764729 | -0.2885 | 0.0738 | 9.36E-05 |
| rs7649970 | -0.2843 | 0.0741 | 1.26E-04 |
| rs765876 | -0.2826 | 0.0741 | 1.37E-04 |
| rs770082 | -0.2890 | 0.0740 | 9.43E-05 |
| rs7723426 | -0.2900 | 0.0736 | 8.14E-05 |
| rs7755574 | -0.2831 | 0.0741 | 1.34E-04 |
| rs7811825 | -0.2914 | 0.0734 | 7.22E-05 |

| | | | |
|---|---|---|---|
| rs7845090 | -0.2849 | 0.0742 | 1.24E-04 |
| rs78784145 | -0.2856 | 0.0741 | 1.15E-04 |
| rs7930275 | -0.2782 | 0.0739 | 1.68E-04 |
| rs79535757 | -0.2808 | 0.0739 | 1.45E-04 |
| rs7978353 | -0.2863 | 0.0741 | 1.12E-04 |
| rs7987928 | -0.2810 | 0.0741 | 1.49E-04 |
| rs79921461 | -0.2832 | 0.0741 | 1.33E-04 |
| rs8087074 | -0.2869 | 0.0740 | 1.07E-04 |
| rs811715 | -0.2856 | 0.0741 | 1.16E-04 |
| rs815163 | -0.2865 | 0.0741 | 1.10E-04 |
| rs8192675 | -0.2821 | 0.0742 | 1.42E-04 |
| rs846545 | -0.2837 | 0.0741 | 1.30E-04 |
| rs854917 | -0.2801 | 0.0740 | 1.55E-04 |
| rs869400 | -0.2790 | 0.0741 | 1.66E-04 |
| rs879620 | -0.2739 | 0.0740 | 2.16E-04 |
| rs9289630 | -0.2836 | 0.0742 | 1.33E-04 |
| rs9320823 | -0.2805 | 0.0741 | 1.54E-04 |
| rs935166 | -0.2868 | 0.0741 | 1.08E-04 |
| rs9375702 | -0.2817 | 0.0741 | 1.44E-04 |
| rs9378684 | -0.2864 | 0.0740 | 1.09E-04 |
| rs9463511 | -0.2892 | 0.0740 | 9.23E-05 |
| rs9496567 | -0.2904 | 0.0734 | 7.66E-05 |
| rs9515455 | -0.2864 | 0.0740 | 1.10E-04 |
| rs9549099 | -0.2829 | 0.0741 | 1.36E-04 |
| rs9688977 | -0.2844 | 0.0742 | 1.26E-04 |
| rs9843653 | -0.2933 | 0.0741 | 7.64E-05 |
| rs9847186 | -0.2855 | 0.0741 | 1.17E-04 |
| rs9861443 | -0.2876 | 0.0738 | 9.65E-05 |
| All | -0.2835 | 0.0739 | 1.25E-04 |

**Whole body fat mass || id:UKB-a:265**

| SNP | b | se | p |
|---|---|---|---|
| rs10100245 | -0.2659 | 0.0703 | 1.55E-04 |
| rs10128597 | -0.2655 | 0.0703 | 1.57E-04 |
| rs10141106 | -0.2646 | 0.0702 | 1.64E-04 |
| rs10172196 | -0.2668 | 0.0702 | 1.45E-04 |
| rs10237306 | -0.2685 | 0.0701 | 1.29E-04 |
| rs10269774 | -0.2592 | 0.0700 | 2.14E-04 |
| rs10423928 | -0.2680 | 0.0704 | 1.39E-04 |
| rs10490530 | -0.2623 | 0.0701 | 1.83E-04 |
| rs10499014 | -0.2647 | 0.0702 | 1.63E-04 |
| rs10507483 | -0.2655 | 0.0702 | 1.56E-04 |
| rs1056441 | -0.2659 | 0.0702 | 1.53E-04 |
| rs10748028 | -0.2637 | 0.0702 | 1.72E-04 |
| rs10756714 | -0.2575 | 0.0701 | 2.40E-04 |
| rs10756798 | -0.2707 | 0.0700 | 1.10E-04 |
| rs10760724 | -0.2649 | 0.0702 | 1.62E-04 |
| rs10787738 | -0.2686 | 0.0701 | 1.27E-04 |
| rs10820852 | -0.2660 | 0.0702 | 1.51E-04 |
| rs10827289 | -0.2642 | 0.0702 | 1.68E-04 |
| rs10915840 | -0.2682 | 0.0701 | 1.30E-04 |
| rs10938397 | -0.2733 | 0.0705 | 1.06E-04 |
| rs10947793 | -0.2646 | 0.0702 | 1.65E-04 |
| rs10954772 | -0.2685 | 0.0701 | 1.29E-04 |
| rs10999460 | -0.2605 | 0.0700 | 1.97E-04 |
| rs11012732 | -0.2511 | 0.0700 | 3.33E-04 |
| rs11030119 | -0.2595 | 0.0704 | 2.27E-04 |
| rs11150745 | -0.2633 | 0.0702 | 1.78E-04 |
| rs111640872 | -0.2581 | 0.0701 | 2.31E-04 |
| rs11205303 | -0.2748 | 0.0702 | 9.02E-05 |
| rs11205617 | -0.2656 | 0.0703 | 1.57E-04 |
| rs113866544 | -0.2688 | 0.0702 | 1.28E-04 |
| rs11474838 | -0.2626 | 0.0701 | 1.81E-04 |
| rs11538 | -0.2673 | 0.0701 | 1.39E-04 |
| rs11580836 | -0.2579 | 0.0695 | 2.06E-04 |
| rs11603783 | -0.2692 | 0.0700 | 1.21E-04 |
| rs11642015 | -0.2608 | 0.0717 | 2.75E-04 |
| rs11656758 | -0.2554 | 0.0693 | 2.30E-04 |
| rs11695700 | -0.2650 | 0.0702 | 1.61E-04 |
| rs11725731 | -0.2647 | 0.0702 | 1.63E-04 |
| rs11764337 | -0.2692 | 0.0700 | 1.21E-04 |
| rs11782074 | -0.2677 | 0.0701 | 1.35E-04 |
| rs11782341 | -0.2700 | 0.0700 | 1.13E-04 |
| rs11786089 | -0.2631 | 0.0702 | 1.78E-04 |

| | | | |
|---|---|---|---|
| rs1182199 | -0.2708 | 0.0702 | 1.13E-04 |
| rs11824092 | -0.2692 | 0.0700 | 1.19E-04 |
| rs11857221 | -0.2616 | 0.0701 | 1.88E-04 |
| rs11958027 | -0.2653 | 0.0702 | 1.58E-04 |
| rs12031634 | -0.2638 | 0.0702 | 1.72E-04 |
| rs12103006 | -0.2651 | 0.0703 | 1.62E-04 |
| rs12128526 | -0.2556 | 0.0691 | 2.15E-04 |
| rs12140153 | -0.2693 | 0.0702 | 1.26E-04 |
| rs12144626 | -0.2702 | 0.0699 | 1.12E-04 |
| rs1218824 | -0.2659 | 0.0702 | 1.53E-04 |
| rs12254441 | -0.2610 | 0.0701 | 1.95E-04 |
| rs1229984 | -0.2641 | 0.0703 | 1.70E-04 |
| rs12330631 | -0.2669 | 0.0702 | 1.43E-04 |
| rs12339822 | -0.2603 | 0.0700 | 2.02E-04 |
| rs12367809 | -0.2698 | 0.0704 | 1.26E-04 |
| rs12459368 | -0.2643 | 0.0702 | 1.68E-04 |
| rs12475388 | -0.2667 | 0.0702 | 1.45E-04 |
| rs12477385 | -0.2681 | 0.0701 | 1.31E-04 |
| rs12578952 | -0.2614 | 0.0701 | 1.93E-04 |
| rs12610925 | -0.2713 | 0.0700 | 1.06E-04 |
| rs12616638 | -0.2686 | 0.0700 | 1.25E-04 |
| rs12619178 | -0.2658 | 0.0702 | 1.54E-04 |
| rs1268403 | -0.2660 | 0.0702 | 1.51E-04 |
| rs12877270 | -0.2626 | 0.0701 | 1.82E-04 |
| rs12887636 | -0.2663 | 0.0702 | 1.48E-04 |
| rs12890931 | -0.2630 | 0.0702 | 1.78E-04 |
| rs12926311 | -0.2610 | 0.0701 | 1.95E-04 |
| rs12987931 | -0.2671 | 0.0701 | 1.40E-04 |
| rs12992672 | -0.2759 | 0.0704 | 8.81E-05 |
| rs13056506 | -0.2684 | 0.0702 | 1.32E-04 |
| rs13062093 | -0.2654 | 0.0702 | 1.58E-04 |
| rs13067187 | -0.2637 | 0.0702 | 1.72E-04 |
| rs13135092 | -0.2535 | 0.0697 | 2.74E-04 |
| rs13156484 | -0.2640 | 0.0702 | 1.70E-04 |
| rs13174863 | -0.2632 | 0.0702 | 1.78E-04 |
| rs13333747 | -0.2644 | 0.0702 | 1.66E-04 |
| rs1335055 | -0.2630 | 0.0702 | 1.78E-04 |
| rs13427822 | -0.2637 | 0.0702 | 1.73E-04 |
| rs1383723 | -0.2653 | 0.0702 | 1.57E-04 |
| rs1441264 | -0.2629 | 0.0702 | 1.82E-04 |
| rs1446585 | -0.2873 | 0.0673 | 1.99E-05 |
| rs1470749 | -0.2672 | 0.0702 | 1.39E-04 |
| rs1477290 | -0.2705 | 0.0702 | 1.17E-04 |
| rs147730268 | -0.2723 | 0.0701 | 1.02E-04 |
| rs1491905 | -0.2568 | 0.0694 | 2.15E-04 |

| | | | |
|---|---|---|---|
| rs1503527 | -0.2663 | 0.0702 | 1.49E-04 |
| rs1568489 | -0.2707 | 0.0700 | 1.09E-04 |
| rs1652376 | -0.2586 | 0.0702 | 2.28E-04 |
| rs1653892 | -0.2679 | 0.0702 | 1.34E-04 |
| rs17014332 | -0.2684 | 0.0701 | 1.28E-04 |
| rs17024393 | -0.2656 | 0.0703 | 1.59E-04 |
| rs17218879 | -0.2621 | 0.0701 | 1.85E-04 |
| rs1724551 | -0.2645 | 0.0702 | 1.65E-04 |
| rs17265513 | -0.2711 | 0.0700 | 1.07E-04 |
| rs17770336 | -0.2607 | 0.0702 | 2.05E-04 |
| rs1782508 | -0.2653 | 0.0702 | 1.58E-04 |
| rs1783541 | -0.2694 | 0.0700 | 1.19E-04 |
| rs1834144 | -0.2549 | 0.0695 | 2.46E-04 |
| rs1840126 | -0.2592 | 0.0697 | 2.02E-04 |
| rs1879529 | -0.2678 | 0.0701 | 1.35E-04 |
| rs1881505 | -0.2644 | 0.0701 | 1.62E-04 |
| rs1927626 | -0.2629 | 0.0702 | 1.80E-04 |
| rs1928496 | -0.2642 | 0.0702 | 1.68E-04 |
| rs1964599 | -0.2652 | 0.0702 | 1.59E-04 |
| rs2016469 | -0.2616 | 0.0700 | 1.87E-04 |
| rs2072413 | -0.2624 | 0.0701 | 1.81E-04 |
| rs2102278 | -0.2605 | 0.0700 | 1.97E-04 |
| rs2172131 | -0.2610 | 0.0701 | 1.98E-04 |
| rs217672 | -0.2674 | 0.0701 | 1.38E-04 |
| rs2192527 | -0.2646 | 0.0703 | 1.66E-04 |
| rs2239647 | -0.2679 | 0.0702 | 1.36E-04 |
| rs2253310 | -0.2716 | 0.0701 | 1.06E-04 |
| rs2287214 | -0.2663 | 0.0703 | 1.50E-04 |
| rs2289379 | -0.2659 | 0.0702 | 1.53E-04 |
| rs2292238 | -0.2659 | 0.0702 | 1.52E-04 |
| rs2307111 | -0.2635 | 0.0704 | 1.81E-04 |
| rs2370982 | -0.2660 | 0.0703 | 1.56E-04 |
| rs2374947 | -0.2690 | 0.0700 | 1.21E-04 |
| rs2425024 | -0.2662 | 0.0702 | 1.50E-04 |
| rs2439823 | -0.2664 | 0.0703 | 1.50E-04 |
| rs245775 | -0.2673 | 0.0702 | 1.41E-04 |
| rs2479958 | -0.2660 | 0.0702 | 1.52E-04 |
| rs2482398 | -0.2711 | 0.0698 | 1.03E-04 |
| rs2494196 | -0.2645 | 0.0703 | 1.68E-04 |
| rs2499468 | -0.2657 | 0.0702 | 1.54E-04 |
| rs252749 | -0.2593 | 0.0700 | 2.13E-04 |
| rs2606228 | -0.2683 | 0.0701 | 1.29E-04 |
| rs2678204 | -0.2579 | 0.0701 | 2.35E-04 |
| rs2814943 | -0.2761 | 0.0704 | 8.74E-05 |
| rs28366156 | -0.2690 | 0.0701 | 1.26E-04 |

| | | | |
|---|---|---|---|
| rs2861685 | -0.2615 | 0.0701 | 1.93E-04 |
| rs28636067 | -0.2640 | 0.0702 | 1.69E-04 |
| rs2869950 | -0.2562 | 0.0698 | 2.43E-04 |
| rs2917705 | -0.2651 | 0.0702 | 1.60E-04 |
| rs2954021 | -0.2649 | 0.0702 | 1.62E-04 |
| rs314279 | -0.2618 | 0.0701 | 1.88E-04 |
| rs34045288 | -0.2668 | 0.0703 | 1.48E-04 |
| rs34292685 | -0.2641 | 0.0702 | 1.68E-04 |
| rs34517439 | -0.2631 | 0.0701 | 1.75E-04 |
| rs34769775 | -0.2639 | 0.0702 | 1.72E-04 |
| rs34966008 | -0.2666 | 0.0702 | 1.47E-04 |
| rs35589149 | -0.2610 | 0.0701 | 1.95E-04 |
| rs35851183 | -0.2611 | 0.0701 | 1.95E-04 |
| rs35882248 | -0.2603 | 0.0702 | 2.07E-04 |
| rs3730071 | -0.2685 | 0.0701 | 1.29E-04 |
| rs3737992 | -0.2703 | 0.0699 | 1.11E-04 |
| rs3766823 | -0.2657 | 0.0702 | 1.55E-04 |
| rs3784699 | -0.2601 | 0.0702 | 2.13E-04 |
| rs3803286 | -0.2694 | 0.0701 | 1.23E-04 |
| rs3814883 | -0.2547 | 0.0701 | 2.79E-04 |
| rs3826408 | -0.2717 | 0.0699 | 1.02E-04 |
| rs3864311 | -0.2671 | 0.0701 | 1.41E-04 |
| rs40071 | -0.2644 | 0.0702 | 1.67E-04 |
| rs412522 | -0.2652 | 0.0702 | 1.60E-04 |
| rs429358 | -0.2681 | 0.0702 | 1.35E-04 |
| rs4471907 | -0.2659 | 0.0702 | 1.52E-04 |
| rs4477562 | -0.2644 | 0.0703 | 1.68E-04 |
| rs4502882 | -0.2652 | 0.0702 | 1.59E-04 |
| rs4503172 | -0.2670 | 0.0702 | 1.42E-04 |
| rs4549080 | -0.2668 | 0.0702 | 1.44E-04 |
| rs4657796 | -0.2632 | 0.0702 | 1.77E-04 |
| rs4670172 | -0.2624 | 0.0701 | 1.83E-04 |
| rs4673617 | -0.2678 | 0.0701 | 1.35E-04 |
| rs4718964 | -0.2647 | 0.0703 | 1.65E-04 |
| rs4790841 | -0.2660 | 0.0703 | 1.55E-04 |
| rs4794222 | -0.2641 | 0.0702 | 1.68E-04 |
| rs4864201 | -0.2656 | 0.0702 | 1.55E-04 |
| rs4876611 | -0.2618 | 0.0702 | 1.93E-04 |
| rs4900471 | -0.2678 | 0.0701 | 1.34E-04 |
| rs4958702 | -0.2643 | 0.0702 | 1.67E-04 |
| rs543874 | -0.2815 | 0.0707 | 6.77E-05 |
| rs55637757 | -0.2633 | 0.0701 | 1.74E-04 |
| rs55714539 | -0.2649 | 0.0703 | 1.64E-04 |
| rs55726687 | -0.2753 | 0.0694 | 7.32E-05 |
| rs56288810 | -0.2689 | 0.0701 | 1.24E-04 |

| | | | |
|---|---|---|---|
| rs56361700 | -0.2621 | 0.0701 | 1.84E-04 |
| rs57636386 | -0.2631 | 0.0703 | 1.83E-04 |
| rs59066241 | -0.2664 | 0.0702 | 1.47E-04 |
| rs59104534 | -0.2656 | 0.0702 | 1.55E-04 |
| rs59227842 | -0.2672 | 0.0703 | 1.45E-04 |
| rs6029180 | -0.2629 | 0.0701 | 1.79E-04 |
| rs60898782 | -0.2680 | 0.0701 | 1.32E-04 |
| rs61649432 | -0.2612 | 0.0700 | 1.91E-04 |
| rs61813293 | -0.2642 | 0.0701 | 1.65E-04 |
| rs61903695 | -0.2655 | 0.0702 | 1.55E-04 |
| rs62025854 | -0.2651 | 0.0703 | 1.63E-04 |
| rs62037365 | -0.2525 | 0.0701 | 3.18E-04 |
| rs62178054 | -0.2655 | 0.0703 | 1.57E-04 |
| rs62246314 | -0.2646 | 0.0702 | 1.63E-04 |
| rs62261725 | -0.2590 | 0.0700 | 2.18E-04 |
| rs62343137 | -0.2660 | 0.0703 | 1.53E-04 |
| rs62396185 | -0.2502 | 0.0702 | 3.63E-04 |
| rs62407569 | -0.2695 | 0.0700 | 1.19E-04 |
| rs62499696 | -0.2671 | 0.0701 | 1.40E-04 |
| rs6545714 | -0.2687 | 0.0702 | 1.30E-04 |
| rs6561937 | -0.2630 | 0.0701 | 1.77E-04 |
| rs6567160 | -0.2577 | 0.0707 | 2.67E-04 |
| rs6575340 | -0.2644 | 0.0703 | 1.69E-04 |
| rs6598540 | -0.2620 | 0.0702 | 1.88E-04 |
| rs6601527 | -0.2706 | 0.0699 | 1.08E-04 |
| rs6699744 | -0.2611 | 0.0702 | 2.01E-04 |
| rs6717858 | -0.2623 | 0.0702 | 1.89E-04 |
| rs67536595 | -0.2649 | 0.0702 | 1.61E-04 |
| rs67913249 | -0.2679 | 0.0701 | 1.32E-04 |
| rs6861649 | -0.2608 | 0.0701 | 1.98E-04 |
| rs6875585 | -0.2730 | 0.0695 | 8.61E-05 |
| rs6898357 | -0.2628 | 0.0701 | 1.79E-04 |
| rs6973656 | -0.2681 | 0.0701 | 1.32E-04 |
| rs704061 | -0.2716 | 0.0699 | 1.03E-04 |
| rs7094644 | -0.2716 | 0.0698 | 9.97E-05 |
| rs7124681 | -0.2656 | 0.0704 | 1.61E-04 |
| rs713586 | -0.2624 | 0.0706 | 2.04E-04 |
| rs7171864 | -0.2699 | 0.0701 | 1.17E-04 |
| rs7193783 | -0.2650 | 0.0702 | 1.62E-04 |
| rs7238896 | -0.2612 | 0.0701 | 1.93E-04 |
| rs724016 | -0.2667 | 0.0705 | 1.56E-04 |
| rs7259070 | -0.2636 | 0.0702 | 1.72E-04 |
| rs72755233 | -0.2684 | 0.0699 | 1.23E-04 |
| rs72820274 | -0.2650 | 0.0702 | 1.61E-04 |
| rs72906282 | -0.2643 | 0.0702 | 1.66E-04 |

| | | | |
|---|---|---|---|
| rs72917533 | -0.2651 | 0.0702 | 1.60E-04 |
| rs72976986 | -0.2572 | 0.0696 | 2.17E-04 |
| rs73041988 | -0.2650 | 0.0703 | 1.62E-04 |
| rs7313748 | -0.2689 | 0.0701 | 1.24E-04 |
| rs73142879 | -0.2630 | 0.0702 | 1.80E-04 |
| rs73213484 | -0.2627 | 0.0702 | 1.82E-04 |
| rs7460093 | -0.2620 | 0.0700 | 1.83E-04 |
| rs750090 | -0.2659 | 0.0702 | 1.52E-04 |
| rs75412871 | -0.2641 | 0.0702 | 1.68E-04 |
| rs75557510 | -0.2695 | 0.0701 | 1.21E-04 |
| rs75641275 | -0.2639 | 0.0702 | 1.70E-04 |
| rs76040172 | -0.2577 | 0.0698 | 2.24E-04 |
| rs7613261 | -0.2670 | 0.0703 | 1.44E-04 |
| rs7649970 | -0.2662 | 0.0703 | 1.52E-04 |
| rs765875 | -0.2639 | 0.0702 | 1.70E-04 |
| rs7845090 | -0.2666 | 0.0703 | 1.50E-04 |
| rs7933085 | -0.2690 | 0.0701 | 1.24E-04 |
| rs7987928 | -0.2623 | 0.0702 | 1.85E-04 |
| rs80135947 | -0.2533 | 0.0698 | 2.87E-04 |
| rs801738 | -0.2536 | 0.0695 | 2.61E-04 |
| rs8087074 | -0.2680 | 0.0701 | 1.32E-04 |
| rs8112818 | -0.2649 | 0.0703 | 1.63E-04 |
| rs8118253 | -0.2640 | 0.0702 | 1.69E-04 |
| rs815163 | -0.2676 | 0.0702 | 1.37E-04 |
| rs8192675 | -0.2637 | 0.0702 | 1.73E-04 |
| rs846546 | -0.2651 | 0.0702 | 1.59E-04 |
| rs854917 | -0.2618 | 0.0701 | 1.89E-04 |
| rs869400 | -0.2611 | 0.0701 | 1.96E-04 |
| rs879620 | -0.2554 | 0.0701 | 2.69E-04 |
| rs9320823 | -0.2619 | 0.0703 | 1.93E-04 |
| rs9375702 | -0.2632 | 0.0702 | 1.78E-04 |
| rs9515455 | -0.2681 | 0.0701 | 1.32E-04 |
| rs9569933 | -0.2597 | 0.0699 | 2.03E-04 |
| rs9584855 | -0.2677 | 0.0701 | 1.35E-04 |
| rs9610311 | -0.2634 | 0.0701 | 1.73E-04 |
| rs9843007 | -0.2644 | 0.0702 | 1.66E-04 |
| rs9843653 | -0.2740 | 0.0702 | 9.49E-05 |
| rs9861443 | -0.2692 | 0.0699 | 1.17E-04 |
| rs9968060 | -0.2665 | 0.0702 | 1.47E-04 |
| All | -0.2650 | 0.0700 | 1.53E-04 |

**Leg fat percentage (right) || id:UKB-a:274**

| SNP | b | se | p |
|---|---|---|---|
| rs1013402 | -0.4623 | 0.1261 | 2.46E-04 |
| rs10209821 | -0.4767 | 0.1259 | 1.53E-04 |
| rs10423928 | -0.4775 | 0.1262 | 1.55E-04 |
| rs1046080 | -0.4381 | 0.1250 | 4.57E-04 |
| rs10499014 | -0.4707 | 0.1260 | 1.87E-04 |
| rs1056441 | -0.4729 | 0.1259 | 1.73E-04 |
| rs10732335 | -0.4561 | 0.1260 | 2.95E-04 |
| rs10756798 | -0.4835 | 0.1255 | 1.17E-04 |
| rs10774018 | -0.4697 | 0.1259 | 1.91E-04 |
| rs10787738 | -0.4777 | 0.1257 | 1.44E-04 |
| rs10835186 | -0.4640 | 0.1256 | 2.20E-04 |
| rs10883026 | -0.4704 | 0.1260 | 1.89E-04 |
| rs10938397 | -0.4853 | 0.1264 | 1.23E-04 |
| rs10995427 | -0.4633 | 0.1256 | 2.24E-04 |
| rs11012732 | -0.4430 | 0.1257 | 4.27E-04 |
| rs11150745 | -0.4680 | 0.1259 | 2.02E-04 |
| rs111743285 | -0.4604 | 0.1255 | 2.43E-04 |
| rs11187838 | -0.4764 | 0.1261 | 1.57E-04 |
| rs11264489 | -0.4726 | 0.1259 | 1.74E-04 |
| rs1127100 | -0.4679 | 0.1259 | 2.03E-04 |
| rs112852122 | -0.4845 | 0.1250 | 1.07E-04 |
| rs11474838 | -0.4665 | 0.1258 | 2.08E-04 |
| rs115584674 | -0.4713 | 0.1257 | 1.78E-04 |
| rs11642015 | -0.4631 | 0.1281 | 3.00E-04 |
| rs11688365 | -0.4758 | 0.1257 | 1.54E-04 |
| rs11695700 | -0.4710 | 0.1259 | 1.83E-04 |
| rs117068593 | -0.4761 | 0.1258 | 1.54E-04 |
| rs11782341 | -0.4803 | 0.1254 | 1.28E-04 |
| rs11824092 | -0.4804 | 0.1254 | 1.28E-04 |
| rs11841757 | -0.4822 | 0.1255 | 1.22E-04 |
| rs11866219 | -0.4736 | 0.1259 | 1.70E-04 |
| rs12042959 | -0.4708 | 0.1260 | 1.86E-04 |
| rs12049202 | -0.4807 | 0.1255 | 1.28E-04 |
| rs12055234 | -0.4792 | 0.1256 | 1.36E-04 |
| rs12103006 | -0.4709 | 0.1259 | 1.85E-04 |
| rs12124126 | -0.4638 | 0.1259 | 2.29E-04 |
| rs12140153 | -0.4774 | 0.1258 | 1.48E-04 |
| rs12148513 | -0.4812 | 0.1249 | 1.16E-04 |
| rs1225004 | -0.4714 | 0.1260 | 1.83E-04 |
| rs1225404 | -0.4831 | 0.1251 | 1.13E-04 |
| rs12292614 | -0.4715 | 0.1259 | 1.81E-04 |
| rs1229984 | -0.4695 | 0.1260 | 1.95E-04 |

| | | | |
|---|---|---|---|
| rs12316080 | -0.4748 | 0.1259 | 1.63E-04 |
| rs12339822 | -0.4614 | 0.1256 | 2.40E-04 |
| rs12355619 | -0.4671 | 0.1258 | 2.05E-04 |
| rs12367809 | -0.4772 | 0.1260 | 1.52E-04 |
| rs12438864 | -0.4740 | 0.1260 | 1.68E-04 |
| rs12459676 | -0.4751 | 0.1258 | 1.60E-04 |
| rs12462975 | -0.4619 | 0.1254 | 2.29E-04 |
| rs12477385 | -0.4785 | 0.1257 | 1.40E-04 |
| rs12503232 | -0.4550 | 0.1254 | 2.87E-04 |
| rs12548773 | -0.4745 | 0.1259 | 1.64E-04 |
| rs12583872 | -0.4721 | 0.1259 | 1.78E-04 |
| rs12619178 | -0.4727 | 0.1260 | 1.75E-04 |
| rs12679106 | -0.4765 | 0.1260 | 1.56E-04 |
| rs12716979 | -0.4356 | 0.1217 | 3.45E-04 |
| rs12806052 | -0.4710 | 0.1260 | 1.86E-04 |
| rs12890931 | -0.4669 | 0.1259 | 2.08E-04 |
| rs1296328 | -0.4686 | 0.1259 | 1.99E-04 |
| rs13056506 | -0.4759 | 0.1258 | 1.55E-04 |
| rs13062093 | -0.4723 | 0.1261 | 1.79E-04 |
| rs13135092 | -0.4497 | 0.1249 | 3.18E-04 |
| rs13427822 | -0.4684 | 0.1259 | 2.00E-04 |
| rs136309 | -0.4766 | 0.1257 | 1.50E-04 |
| rs138556772 | -0.4670 | 0.1251 | 1.90E-04 |
| rs1441264 | -0.4672 | 0.1259 | 2.07E-04 |
| rs1446585 | -0.5163 | 0.1203 | 1.78E-05 |
| rs1460940 | -0.4730 | 0.1262 | 1.78E-04 |
| rs1471740 | -0.4749 | 0.1259 | 1.62E-04 |
| rs1477290 | -0.4806 | 0.1258 | 1.34E-04 |
| rs147730268 | -0.4812 | 0.1256 | 1.27E-04 |
| rs1486921 | -0.4694 | 0.1259 | 1.92E-04 |
| rs1490382 | -0.4677 | 0.1258 | 2.02E-04 |
| rs1503527 | -0.4739 | 0.1259 | 1.68E-04 |
| rs1524445 | -0.4702 | 0.1259 | 1.89E-04 |
| rs1538742 | -0.4680 | 0.1260 | 2.04E-04 |
| rs1568489 | -0.4803 | 0.1254 | 1.28E-04 |
| rs1608113 | -0.4722 | 0.1255 | 1.68E-04 |
| rs1609303 | -0.4714 | 0.1260 | 1.83E-04 |
| rs1653892 | -0.4771 | 0.1258 | 1.49E-04 |
| rs17024393 | -0.4713 | 0.1260 | 1.83E-04 |
| rs17222916 | -0.4703 | 0.1259 | 1.87E-04 |
| rs17522122 | -0.4757 | 0.1258 | 1.56E-04 |
| rs1783541 | -0.4801 | 0.1255 | 1.31E-04 |
| rs1788808 | -0.4618 | 0.1258 | 2.42E-04 |
| rs1881934 | -0.4681 | 0.1259 | 2.00E-04 |
| rs1928496 | -0.4696 | 0.1260 | 1.93E-04 |

| | | | |
|---|---|---|---|
| rs2033530 | -0.4779 | 0.1257 | 1.43E-04 |
| rs2034768 | -0.4683 | 0.1260 | 2.01E-04 |
| rs208477 | -0.4740 | 0.1258 | 1.65E-04 |
| rs2089054 | -0.4730 | 0.1259 | 1.71E-04 |
| rs2161796 | -0.4536 | 0.1246 | 2.72E-04 |
| rs2172131 | -0.4630 | 0.1258 | 2.32E-04 |
| rs2241743 | -0.4658 | 0.1258 | 2.13E-04 |
| rs228443 | -0.4709 | 0.1259 | 1.84E-04 |
| rs2285639 | -0.4742 | 0.1259 | 1.67E-04 |
| rs2287894 | -0.4742 | 0.1258 | 1.64E-04 |
| rs2289379 | -0.4731 | 0.1260 | 1.73E-04 |
| rs2307111 | -0.4684 | 0.1261 | 2.03E-04 |
| rs2318543 | -0.4721 | 0.1259 | 1.77E-04 |
| rs2319849 | -0.4746 | 0.1256 | 1.58E-04 |
| rs2370982 | -0.4727 | 0.1261 | 1.78E-04 |
| rs2398861 | -0.4585 | 0.1255 | 2.59E-04 |
| rs241459 | -0.4730 | 0.1260 | 1.75E-04 |
| rs2436772 | -0.4751 | 0.1259 | 1.62E-04 |
| rs2606228 | -0.4773 | 0.1256 | 1.45E-04 |
| rs2644135 | -0.4577 | 0.1256 | 2.69E-04 |
| rs2702123 | -0.4733 | 0.1259 | 1.70E-04 |
| rs2711117 | -0.4835 | 0.1248 | 1.07E-04 |
| rs2721965 | -0.4648 | 0.1259 | 2.22E-04 |
| rs2725370 | -0.4793 | 0.1257 | 1.37E-04 |
| rs2802295 | -0.4794 | 0.1254 | 1.32E-04 |
| rs2814967 | -0.4771 | 0.1261 | 1.55E-04 |
| rs2861685 | -0.4645 | 0.1258 | 2.22E-04 |
| rs2867131 | -0.4884 | 0.1257 | 1.03E-04 |
| rs28726372 | -0.4625 | 0.1250 | 2.16E-04 |
| rs2955242 | -0.4592 | 0.1246 | 2.29E-04 |
| rs2963457 | -0.4756 | 0.1258 | 1.57E-04 |
| rs34117444 | -0.4705 | 0.1259 | 1.87E-04 |
| rs34234296 | -0.4742 | 0.1258 | 1.64E-04 |
| rs34424 | -0.4705 | 0.1261 | 1.90E-04 |
| rs34769775 | -0.4689 | 0.1259 | 1.96E-04 |
| rs350832 | -0.4528 | 0.1253 | 3.01E-04 |
| rs362307 | -0.4734 | 0.1258 | 1.68E-04 |
| rs3749473 | -0.4671 | 0.1259 | 2.06E-04 |
| rs3764002 | -0.4718 | 0.1264 | 1.90E-04 |
| rs3770781 | -0.4557 | 0.1249 | 2.63E-04 |
| rs3784699 | -0.4622 | 0.1260 | 2.44E-04 |
| rs3803286 | -0.4790 | 0.1258 | 1.40E-04 |
| rs4073582 | -0.4508 | 0.1245 | 2.95E-04 |
| rs429358 | -0.4782 | 0.1260 | 1.48E-04 |
| rs4322261 | -0.4695 | 0.1259 | 1.92E-04 |

| | | | |
|---|---|---|---|
| rs4467770 | -0.4709 | 0.1260 | 1.86E-04 |
| rs4471907 | -0.4732 | 0.1259 | 1.71E-04 |
| rs4503172 | -0.4761 | 0.1259 | 1.56E-04 |
| rs450779 | -0.4757 | 0.1258 | 1.55E-04 |
| rs45593734 | -0.4588 | 0.1248 | 2.38E-04 |
| rs4682711 | -0.4756 | 0.1258 | 1.57E-04 |
| rs4690324 | -0.4649 | 0.1256 | 2.15E-04 |
| rs4718964 | -0.4704 | 0.1260 | 1.89E-04 |
| rs4737526 | -0.4642 | 0.1257 | 2.20E-04 |
| rs4759318 | -0.4758 | 0.1257 | 1.54E-04 |
| rs4762962 | -0.4685 | 0.1259 | 1.98E-04 |
| rs4790292 | -0.4718 | 0.1261 | 1.83E-04 |
| rs4894808 | -0.4660 | 0.1257 | 2.08E-04 |
| rs497418 | -0.4679 | 0.1258 | 2.01E-04 |
| rs4981693 | -0.4726 | 0.1259 | 1.74E-04 |
| rs501444 | -0.4680 | 0.1259 | 2.01E-04 |
| rs513280 | -0.4846 | 0.1250 | 1.05E-04 |
| rs525101 | -0.4762 | 0.1258 | 1.54E-04 |
| rs529200 | -0.4687 | 0.1259 | 1.96E-04 |
| rs538656 | -0.4597 | 0.1261 | 2.67E-04 |
| rs539515 | -0.4856 | 0.1259 | 1.14E-04 |
| rs55714539 | -0.4712 | 0.1261 | 1.87E-04 |
| rs56375099 | -0.4678 | 0.1259 | 2.02E-04 |
| rs56399737 | -0.4673 | 0.1258 | 2.05E-04 |
| rs56803094 | -0.4687 | 0.1258 | 1.96E-04 |
| rs57803 | -0.4769 | 0.1256 | 1.47E-04 |
| rs59104534 | -0.4723 | 0.1259 | 1.76E-04 |
| rs59227842 | -0.4748 | 0.1261 | 1.66E-04 |
| rs6021948 | -0.4668 | 0.1259 | 2.09E-04 |
| rs6103254 | -0.4694 | 0.1259 | 1.93E-04 |
| rs61826877 | -0.4768 | 0.1257 | 1.49E-04 |
| rs62011698 | -0.4649 | 0.1257 | 2.16E-04 |
| rs62261725 | -0.4591 | 0.1257 | 2.60E-04 |
| rs62443626 | -0.4696 | 0.1259 | 1.91E-04 |
| rs62444907 | -0.4759 | 0.1258 | 1.55E-04 |
| rs6561575 | -0.4800 | 0.1254 | 1.30E-04 |
| rs6561937 | -0.4664 | 0.1259 | 2.13E-04 |
| rs6575340 | -0.4699 | 0.1260 | 1.92E-04 |
| rs6674970 | -0.4800 | 0.1254 | 1.28E-04 |
| rs670104 | -0.4682 | 0.1259 | 1.99E-04 |
| rs6711584 | -0.4659 | 0.1258 | 2.13E-04 |
| rs6717858 | -0.4658 | 0.1260 | 2.20E-04 |
| rs6720868 | -0.4621 | 0.1259 | 2.42E-04 |
| rs6723465 | -0.4697 | 0.1259 | 1.91E-04 |
| rs6739755 | -0.4757 | 0.1259 | 1.58E-04 |

| | | | |
|---|---|---|---|
| rs6861649 | -0.4630 | 0.1257 | 2.30E-04 |
| rs6888037 | -0.4903 | 0.1259 | 9.90E-05 |
| rs6925197 | -0.4620 | 0.1255 | 2.31E-04 |
| rs6977416 | -0.4744 | 0.1258 | 1.63E-04 |
| rs6992604 | -0.4749 | 0.1258 | 1.60E-04 |
| rs7006629 | -0.4737 | 0.1259 | 1.68E-04 |
| rs7027304 | -0.4701 | 0.1260 | 1.92E-04 |
| rs704061 | -0.4813 | 0.1253 | 1.23E-04 |
| rs7117842 | -0.4666 | 0.1259 | 2.10E-04 |
| rs7124681 | -0.4728 | 0.1263 | 1.82E-04 |
| rs7164782 | -0.4702 | 0.1261 | 1.93E-04 |
| rs719802 | -0.4803 | 0.1258 | 1.34E-04 |
| rs7206608 | -0.4743 | 0.1259 | 1.64E-04 |
| rs7250833 | -0.4724 | 0.1260 | 1.78E-04 |
| rs7259070 | -0.4687 | 0.1258 | 1.93E-04 |
| rs72677847 | -0.4707 | 0.1259 | 1.85E-04 |
| rs73213501 | -0.4699 | 0.1259 | 1.90E-04 |
| rs7323 | -0.4692 | 0.1260 | 1.97E-04 |
| rs7498044 | -0.4721 | 0.1259 | 1.77E-04 |
| rs7498665 | -0.4533 | 0.1253 | 2.96E-04 |
| rs75499503 | -0.4643 | 0.1260 | 2.30E-04 |
| rs75557510 | -0.4806 | 0.1257 | 1.32E-04 |
| rs7591494 | -0.4758 | 0.1258 | 1.56E-04 |
| rs76040172 | -0.4571 | 0.1252 | 2.62E-04 |
| rs7649970 | -0.4746 | 0.1262 | 1.69E-04 |
| rs76824303 | -0.4716 | 0.1259 | 1.80E-04 |
| rs7700107 | -0.4485 | 0.1242 | 3.06E-04 |
| rs77162980 | -0.4675 | 0.1260 | 2.06E-04 |
| rs7719067 | -0.4698 | 0.1259 | 1.90E-04 |
| rs7774 | -0.4689 | 0.1259 | 1.96E-04 |
| rs7826830 | -0.4578 | 0.1249 | 2.46E-04 |
| rs7831711 | -0.4657 | 0.1257 | 2.12E-04 |
| rs7893571 | -0.4818 | 0.1252 | 1.18E-04 |
| rs7896727 | -0.4772 | 0.1258 | 1.49E-04 |
| rs79219212 | -0.4769 | 0.1257 | 1.47E-04 |
| rs7978070 | -0.4683 | 0.1259 | 1.98E-04 |
| rs7987928 | -0.4660 | 0.1258 | 2.13E-04 |
| rs8073510 | -0.4518 | 0.1250 | 3.02E-04 |
| rs8112818 | -0.4709 | 0.1260 | 1.86E-04 |
| rs819168 | -0.4691 | 0.1260 | 1.96E-04 |
| rs874758 | -0.4698 | 0.1259 | 1.91E-04 |
| rs878206 | -0.4677 | 0.1259 | 2.03E-04 |
| rs879620 | -0.4574 | 0.1254 | 2.65E-04 |
| rs9320823 | -0.4650 | 0.1261 | 2.28E-04 |
| rs9508857 | -0.4693 | 0.1259 | 1.93E-04 |

| | | | |
|---|---|---|---|
| rs9515455 | -0.4793 | 0.1259 | 1.40E-04 |
| rs972283 | -0.4803 | 0.1258 | 1.35E-04 |
| rs9803921 | -0.4778 | 0.1256 | 1.43E-04 |
| rs9839081 | -0.4740 | 0.1259 | 1.66E-04 |
| rs9843653 | -0.4880 | 0.1259 | 1.06E-04 |
| rs9880079 | -0.4809 | 0.1252 | 1.23E-04 |
| rs9902386 | -0.4814 | 0.1257 | 1.28E-04 |
| rs9906944 | -0.4677 | 0.1260 | 2.05E-04 |
| All | -0.4709 | 0.1255 | 1.75E-04 |

**Arm fat mass (right) || id:UKB-a:283**

| SNP | b | se | p |
|---|---|---|---|
| rs10100245 | -0.272 | 0.072 | 1.64E-04 |
| rs10187101 | -0.274 | 0.072 | 1.40E-04 |
| rs10236214 | -0.267 | 0.072 | 2.01E-04 |
| rs10237306 | -0.274 | 0.072 | 1.37E-04 |
| rs10237317 | -0.270 | 0.072 | 1.77E-04 |
| rs10259620 | -0.276 | 0.072 | 1.18E-04 |
| rs10269774 | -0.265 | 0.072 | 2.19E-04 |
| rs10404726 | -0.271 | 0.072 | 1.66E-04 |
| rs10423928 | -0.274 | 0.072 | 1.48E-04 |
| rs10492229 | -0.273 | 0.072 | 1.50E-04 |
| rs10499014 | -0.270 | 0.072 | 1.73E-04 |
| rs1056441 | -0.271 | 0.072 | 1.63E-04 |
| rs10745787 | -0.268 | 0.072 | 1.93E-04 |
| rs10756798 | -0.277 | 0.072 | 1.17E-04 |
| rs10760724 | -0.271 | 0.072 | 1.72E-04 |
| rs10787738 | -0.274 | 0.072 | 1.36E-04 |
| rs10873302 | -0.275 | 0.072 | 1.31E-04 |
| rs10938398 | -0.277 | 0.072 | 1.22E-04 |
| rs10947793 | -0.270 | 0.072 | 1.74E-04 |
| rs10954772 | -0.274 | 0.072 | 1.38E-04 |
| rs10995427 | -0.267 | 0.072 | 2.00E-04 |
| rs11001963 | -0.269 | 0.072 | 1.85E-04 |
| rs11012732 | -0.257 | 0.072 | 3.35E-04 |
| rs11030119 | -0.265 | 0.072 | 2.43E-04 |
| rs11042030 | -0.271 | 0.072 | 1.67E-04 |
| rs11150745 | -0.269 | 0.072 | 1.89E-04 |
| rs11152135 | -0.271 | 0.072 | 1.69E-04 |
| rs11165643 | -0.268 | 0.072 | 1.99E-04 |
| rs11205277 | -0.279 | 0.072 | 1.02E-04 |
| rs11205617 | -0.271 | 0.072 | 1.67E-04 |
| rs11245480 | -0.274 | 0.072 | 1.38E-04 |
| rs112566467 | -0.269 | 0.072 | 1.84E-04 |
| rs1127100 | -0.269 | 0.072 | 1.86E-04 |
| rs113182412 | -0.272 | 0.072 | 1.53E-04 |
| rs113230003 | -0.270 | 0.072 | 1.77E-04 |
| rs113866544 | -0.275 | 0.072 | 1.37E-04 |
| rs11474838 | -0.268 | 0.072 | 1.89E-04 |
| rs11515071 | -0.264 | 0.072 | 2.40E-04 |
| rs11538 | -0.273 | 0.072 | 1.47E-04 |
| rs11609659 | -0.276 | 0.072 | 1.18E-04 |
| rs11642015 | -0.267 | 0.074 | 2.93E-04 |
| rs11757278 | -0.275 | 0.072 | 1.30E-04 |

| | | | |
|---|---|---|---|
| rs11782074 | -0.273 | 0.072 | 1.44E-04 |
| rs11782341 | -0.276 | 0.072 | 1.19E-04 |
| rs11786089 | -0.269 | 0.072 | 1.88E-04 |
| rs1182143 | -0.276 | 0.072 | 1.19E-04 |
| rs11882409 | -0.270 | 0.072 | 1.76E-04 |
| rs11920781 | -0.273 | 0.072 | 1.50E-04 |
| rs12001634 | -0.272 | 0.072 | 1.55E-04 |
| rs12037698 | -0.268 | 0.072 | 1.91E-04 |
| rs12072739 | -0.266 | 0.072 | 2.08E-04 |
| rs12103006 | -0.271 | 0.072 | 1.72E-04 |
| rs12128526 | -0.261 | 0.071 | 2.28E-04 |
| rs12140153 | -0.275 | 0.072 | 1.32E-04 |
| rs1218822 | -0.271 | 0.072 | 1.63E-04 |
| rs12254441 | -0.267 | 0.072 | 2.04E-04 |
| rs12316080 | -0.272 | 0.072 | 1.54E-04 |
| rs12339822 | -0.266 | 0.072 | 2.13E-04 |
| rs12367809 | -0.276 | 0.072 | 1.32E-04 |
| rs12435171 | -0.273 | 0.072 | 1.47E-04 |
| rs12442480 | -0.269 | 0.072 | 1.85E-04 |
| rs12578952 | -0.267 | 0.072 | 2.05E-04 |
| rs1263627 | -0.276 | 0.072 | 1.18E-04 |
| rs12692738 | -0.272 | 0.072 | 1.60E-04 |
| rs1286065 | -0.271 | 0.072 | 1.67E-04 |
| rs12877270 | -0.268 | 0.072 | 1.94E-04 |
| rs12885458 | -0.262 | 0.071 | 2.33E-04 |
| rs12887636 | -0.272 | 0.072 | 1.55E-04 |
| rs1293395 | -0.273 | 0.072 | 1.46E-04 |
| rs12992672 | -0.283 | 0.072 | 8.72E-05 |
| rs13056506 | -0.273 | 0.072 | 1.46E-04 |
| rs13062093 | -0.271 | 0.072 | 1.68E-04 |
| rs13076052 | -0.273 | 0.072 | 1.41E-04 |
| rs13135092 | -0.259 | 0.071 | 2.94E-04 |
| rs13174863 | -0.269 | 0.072 | 1.91E-04 |
| rs1335055 | -0.269 | 0.072 | 1.87E-04 |
| rs13427822 | -0.269 | 0.072 | 1.83E-04 |
| rs1406778 | -0.273 | 0.072 | 1.50E-04 |
| rs1441264 | -0.268 | 0.072 | 1.95E-04 |
| rs1446585 | -0.293 | 0.069 | 2.15E-05 |
| rs1449630 | -0.268 | 0.072 | 1.95E-04 |
| rs1470749 | -0.273 | 0.072 | 1.48E-04 |
| rs1471338 | -0.273 | 0.072 | 1.52E-04 |
| rs1477290 | -0.276 | 0.072 | 1.23E-04 |
| rs147730268 | -0.278 | 0.072 | 1.10E-04 |
| rs1503526 | -0.271 | 0.072 | 1.66E-04 |
| rs1516725 | -0.268 | 0.072 | 1.96E-04 |

| | | | |
|---|---|---|---|
| rs1524445 | -0.270 | 0.072 | 1.76E-04 |
| rs1582931 | -0.270 | 0.072 | 1.83E-04 |
| rs1652376 | -0.264 | 0.072 | 2.36E-04 |
| rs17024393 | -0.271 | 0.072 | 1.68E-04 |
| rs1724551 | -0.270 | 0.072 | 1.75E-04 |
| rs17627049 | -0.275 | 0.072 | 1.27E-04 |
| rs17716502 | -0.267 | 0.072 | 2.08E-04 |
| rs17770336 | -0.266 | 0.072 | 2.20E-04 |
| rs1782508 | -0.271 | 0.072 | 1.68E-04 |
| rs1783541 | -0.275 | 0.072 | 1.28E-04 |
| rs1840126 | -0.265 | 0.072 | 2.16E-04 |
| rs1901241 | -0.274 | 0.072 | 1.37E-04 |
| rs1927626 | -0.268 | 0.072 | 1.90E-04 |
| rs1928496 | -0.270 | 0.072 | 1.78E-04 |
| rs1979440 | -0.270 | 0.072 | 1.73E-04 |
| rs2016469 | -0.267 | 0.072 | 2.01E-04 |
| rs2102278 | -0.266 | 0.072 | 2.08E-04 |
| rs2161796 | -0.261 | 0.071 | 2.52E-04 |
| rs217672 | -0.273 | 0.072 | 1.47E-04 |
| rs2239647 | -0.274 | 0.072 | 1.43E-04 |
| rs2253310 | -0.277 | 0.072 | 1.15E-04 |
| rs2289379 | -0.272 | 0.072 | 1.62E-04 |
| rs2292238 | -0.272 | 0.072 | 1.60E-04 |
| rs2307111 | -0.269 | 0.072 | 1.92E-04 |
| rs2318543 | -0.271 | 0.072 | 1.65E-04 |
| rs2404324 | -0.273 | 0.072 | 1.43E-04 |
| rs2425024 | -0.272 | 0.072 | 1.59E-04 |
| rs2439823 | -0.272 | 0.072 | 1.59E-04 |
| rs245775 | -0.273 | 0.072 | 1.48E-04 |
| rs2479958 | -0.272 | 0.072 | 1.62E-04 |
| rs2494196 | -0.270 | 0.072 | 1.77E-04 |
| rs2499468 | -0.272 | 0.072 | 1.63E-04 |
| rs252749 | -0.265 | 0.072 | 2.22E-04 |
| rs254024 | -0.273 | 0.072 | 1.45E-04 |
| rs2678204 | -0.263 | 0.072 | 2.56E-04 |
| rs273505 | -0.271 | 0.072 | 1.67E-04 |
| rs2814943 | -0.281 | 0.072 | 9.72E-05 |
| rs28366156 | -0.275 | 0.072 | 1.33E-04 |
| rs2861685 | -0.267 | 0.072 | 2.04E-04 |
| rs286818 | -0.270 | 0.072 | 1.75E-04 |
| rs2903738 | -0.274 | 0.072 | 1.37E-04 |
| rs2954021 | -0.271 | 0.072 | 1.72E-04 |
| rs34045288 | -0.273 | 0.072 | 1.56E-04 |
| rs34045894 | -0.274 | 0.072 | 1.34E-04 |
| rs34292685 | -0.270 | 0.072 | 1.79E-04 |

| | | | |
|---|---|---|---|
| rs34517439 | -0.269 | 0.072 | 1.85E-04 |
| rs34769775 | -0.270 | 0.072 | 1.82E-04 |
| rs34811474 | -0.270 | 0.072 | 1.75E-04 |
| rs34966008 | -0.272 | 0.072 | 1.56E-04 |
| rs35537311 | -0.272 | 0.072 | 1.60E-04 |
| rs35679975 | -0.269 | 0.072 | 1.91E-04 |
| rs35851183 | -0.267 | 0.072 | 2.04E-04 |
| rs35882248 | -0.266 | 0.072 | 2.16E-04 |
| rs362307 | -0.272 | 0.072 | 1.58E-04 |
| rs3729793 | -0.267 | 0.072 | 1.98E-04 |
| rs3770821 | -0.273 | 0.072 | 1.53E-04 |
| rs3803286 | -0.275 | 0.072 | 1.31E-04 |
| rs3826408 | -0.278 | 0.072 | 1.05E-04 |
| rs390192 | -0.277 | 0.072 | 1.09E-04 |
| rs3914188 | -0.269 | 0.072 | 1.85E-04 |
| rs429358 | -0.274 | 0.072 | 1.44E-04 |
| rs4402589 | -0.260 | 0.072 | 3.07E-04 |
| rs4474229 | -0.269 | 0.072 | 1.86E-04 |
| rs4477562 | -0.270 | 0.072 | 1.79E-04 |
| rs4502882 | -0.271 | 0.072 | 1.69E-04 |
| rs4549080 | -0.273 | 0.072 | 1.53E-04 |
| rs4673617 | -0.274 | 0.072 | 1.42E-04 |
| rs4740383 | -0.262 | 0.071 | 2.26E-04 |
| rs4757144 | -0.275 | 0.072 | 1.27E-04 |
| rs4776970 | -0.270 | 0.072 | 1.93E-04 |
| rs4777541 | -0.272 | 0.072 | 1.54E-04 |
| rs4790841 | -0.272 | 0.072 | 1.64E-04 |
| rs4832298 | -0.265 | 0.072 | 2.14E-04 |
| rs4856721 | -0.272 | 0.072 | 1.56E-04 |
| rs4864201 | -0.271 | 0.072 | 1.65E-04 |
| rs4886039 | -0.268 | 0.072 | 1.96E-04 |
| rs4921301 | -0.270 | 0.072 | 1.76E-04 |
| rs4926726 | -0.274 | 0.072 | 1.38E-04 |
| rs4958702 | -0.270 | 0.072 | 1.78E-04 |
| rs541577 | -0.274 | 0.072 | 1.42E-04 |
| rs543874 | -0.289 | 0.073 | 6.65E-05 |
| rs55637757 | -0.269 | 0.072 | 1.83E-04 |
| rs55655049 | -0.267 | 0.072 | 1.98E-04 |
| rs55726687 | -0.282 | 0.071 | 7.54E-05 |
| rs56288810 | -0.275 | 0.072 | 1.33E-04 |
| rs56803094 | -0.269 | 0.072 | 1.82E-04 |
| rs57636386 | -0.269 | 0.072 | 1.94E-04 |
| rs58084604 | -0.262 | 0.073 | 3.08E-04 |
| rs59104534 | -0.271 | 0.072 | 1.65E-04 |
| rs60764613 | -0.267 | 0.072 | 2.04E-04 |

| | | | |
|---|---|---|---|
| rs61813324 | -0.268 | 0.072 | 1.88E-04 |
| rs61826865 | -0.274 | 0.072 | 1.40E-04 |
| rs62037365 | -0.258 | 0.072 | 3.34E-04 |
| rs62104483 | -0.263 | 0.072 | 2.50E-04 |
| rs62178054 | -0.271 | 0.072 | 1.67E-04 |
| rs62246314 | -0.270 | 0.072 | 1.73E-04 |
| rs62261725 | -0.264 | 0.072 | 2.33E-04 |
| rs62343137 | -0.272 | 0.072 | 1.62E-04 |
| rs62396185 | -0.258 | 0.072 | 3.14E-04 |
| rs62477684 | -0.274 | 0.072 | 1.46E-04 |
| rs649648 | -0.274 | 0.072 | 1.42E-04 |
| rs6575340 | -0.270 | 0.072 | 1.79E-04 |
| rs6601527 | -0.278 | 0.072 | 1.05E-04 |
| rs6656785 | -0.269 | 0.072 | 1.87E-04 |
| rs66679256 | -0.270 | 0.072 | 1.83E-04 |
| rs668799 | -0.253 | 0.070 | 3.20E-04 |
| rs6699744 | -0.267 | 0.072 | 2.12E-04 |
| rs6739755 | -0.274 | 0.072 | 1.45E-04 |
| rs67602344 | -0.263 | 0.071 | 2.22E-04 |
| rs6861649 | -0.267 | 0.072 | 2.04E-04 |
| rs7030732 | -0.272 | 0.072 | 1.57E-04 |
| rs704061 | -0.278 | 0.072 | 1.08E-04 |
| rs7094644 | -0.277 | 0.072 | 1.08E-04 |
| rs7116641 | -0.273 | 0.072 | 1.57E-04 |
| rs7124681 | -0.271 | 0.072 | 1.70E-04 |
| rs713586 | -0.268 | 0.073 | 2.20E-04 |
| rs7141420 | -0.274 | 0.072 | 1.41E-04 |
| rs7201895 | -0.268 | 0.072 | 1.98E-04 |
| rs7218014 | -0.260 | 0.072 | 2.75E-04 |
| rs7259070 | -0.269 | 0.072 | 1.84E-04 |
| rs72740701 | -0.275 | 0.072 | 1.26E-04 |
| rs72917533 | -0.271 | 0.072 | 1.70E-04 |
| rs73213484 | -0.268 | 0.072 | 1.93E-04 |
| rs745249 | -0.272 | 0.072 | 1.57E-04 |
| rs750090 | -0.272 | 0.072 | 1.61E-04 |
| rs756717 | -0.271 | 0.072 | 1.72E-04 |
| rs7584391 | -0.271 | 0.072 | 1.71E-04 |
| rs76040172 | -0.263 | 0.072 | 2.36E-04 |
| rs7649970 | -0.271 | 0.072 | 1.63E-04 |
| rs765876 | -0.270 | 0.072 | 1.78E-04 |
| rs7811825 | -0.279 | 0.071 | 9.11E-05 |
| rs7845090 | -0.272 | 0.072 | 1.60E-04 |
| rs78719460 | -0.267 | 0.072 | 2.03E-04 |
| rs7915723 | -0.268 | 0.072 | 1.98E-04 |
| rs7917983 | -0.275 | 0.072 | 1.23E-04 |

| | | | |
|---|---|---|---|
| rs7925100 | -0.269 | 0.072 | 1.87E-04 |
| rs7930275 | -0.265 | 0.072 | 2.19E-04 |
| rs79535757 | -0.268 | 0.072 | 1.89E-04 |
| rs7987928 | -0.268 | 0.072 | 1.93E-04 |
| rs801738 | -0.261 | 0.071 | 2.48E-04 |
| rs8026927 | -0.272 | 0.072 | 1.53E-04 |
| rs8087074 | -0.274 | 0.072 | 1.38E-04 |
| rs815163 | -0.274 | 0.072 | 1.43E-04 |
| rs8192675 | -0.269 | 0.072 | 1.85E-04 |
| rs845084 | -0.269 | 0.072 | 1.88E-04 |
| rs846545 | -0.271 | 0.072 | 1.69E-04 |
| rs854917 | -0.267 | 0.072 | 2.00E-04 |
| rs879620 | -0.261 | 0.072 | 2.85E-04 |
| rs9289630 | -0.271 | 0.072 | 1.72E-04 |
| rs9291822 | -0.269 | 0.072 | 1.84E-04 |
| rs9320823 | -0.268 | 0.072 | 2.00E-04 |
| rs935166 | -0.274 | 0.072 | 1.40E-04 |
| rs9375702 | -0.269 | 0.072 | 1.87E-04 |
| rs947088 | -0.272 | 0.072 | 1.55E-04 |
| rs9515455 | -0.274 | 0.072 | 1.42E-04 |
| rs9549099 | -0.270 | 0.072 | 1.76E-04 |
| rs9688977 | -0.271 | 0.072 | 1.64E-04 |
| rs9843007 | -0.270 | 0.072 | 1.76E-04 |
| rs9843653 | -0.280 | 0.072 | 9.88E-05 |
| rs9847186 | -0.273 | 0.072 | 1.51E-04 |
| rs9861443 | -0.275 | 0.072 | 1.24E-04 |
| rs9883422 | -0.266 | 0.072 | 2.04E-04 |
| rs9968060 | -0.272 | 0.072 | 1.56E-04 |
| All | -0.271 | 0.072 | 1.62E-04 |

**Tea intake || id UKB-b 6066**

| SNP | b | se | p |
|---|---|---|---|
| rs10741694 | 0.5939 | 0.1624 | 2.56E-04 |
| rs10752269 | 0.6029 | 0.1599 | 1.62E-04 |
| rs10764990 | 0.5980 | 0.1611 | 2.05E-04 |
| rs11164870 | 0.5363 | 0.1437 | 1.89E-04 |
| rs1156588 | 0.6057 | 0.1595 | 1.46E-04 |
| rs11587444 | 0.6225 | 0.1542 | 5.40E-05 |
| rs12591786 | 0.5712 | 0.1612 | 3.94E-04 |
| rs13282783 | 0.5895 | 0.1623 | 2.80E-04 |
| rs132904 | 0.5531 | 0.1565 | 4.08E-04 |
| rs1481012 | 0.5836 | 0.1631 | 3.47E-04 |
| rs17245213 | 0.5741 | 0.1614 | 3.75E-04 |
| rs17576658 | 0.5826 | 0.1622 | 3.27E-04 |
| rs17685 | 0.6065 | 0.1631 | 2.01E-04 |
| rs2117137 | 0.5928 | 0.1621 | 2.55E-04 |
| rs2273447 | 0.5765 | 0.1622 | 3.78E-04 |
| rs2279844 | 0.5752 | 0.1615 | 3.68E-04 |
| rs2351187 | 0.6002 | 0.1605 | 1.84E-04 |
| rs2472297 | 0.6177 | 0.1885 | 1.05E-03 |
| rs2478875 | 0.6001 | 0.1626 | 2.24E-04 |
| rs2645929 | 0.5877 | 0.1623 | 2.93E-04 |
| rs34619 | 0.6085 | 0.1578 | 1.15E-04 |
| rs4410790 | 0.4841 | 0.1780 | 6.54E-03 |
| rs4808193 | 0.5832 | 0.1627 | 3.39E-04 |
| rs4817505 | 0.5514 | 0.1569 | 4.40E-04 |
| rs56188862 | 0.5876 | 0.1630 | 3.12E-04 |
| rs56348300 | 0.6011 | 0.1605 | 1.80E-04 |
| rs57462170 | 0.5882 | 0.1622 | 2.88E-04 |
| rs57631352 | 0.5978 | 0.1610 | 2.05E-04 |
| rs6829 | 0.5810 | 0.1622 | 3.40E-04 |
| rs713598 | 0.5986 | 0.1627 | 2.33E-04 |
| rs72797284 | 0.6012 | 0.1618 | 2.02E-04 |
| rs7757102 | 0.5547 | 0.1547 | 3.37E-04 |
| rs9302428 | 0.6063 | 0.1589 | 1.36E-04 |
| rs9624470 | 0.5876 | 0.1653 | 3.80E-04 |
| rs9648476 | 0.5752 | 0.1617 | 3.74E-04 |
| rs977474 | 0.5900 | 0.1603 | 2.33E-04 |
| rs9937354 | 0.5730 | 0.1619 | 4.00E-04 |
| All | 0.5852 | 0.1592 | 2.37E-04 |

**Arm fat percentage (left) || id:UKB-a:286**

| SNP | b | se | p |
|---|---|---|---|
| rs10100245 | -0.4147 | 0.1030 | 5.68E-05 |
| rs1013293 | -0.4022 | 0.1025 | 8.72E-05 |
| rs10259620 | -0.4230 | 0.1025 | 3.66E-05 |
| rs10278040 | -0.4119 | 0.1029 | 6.21E-05 |
| rs10423928 | -0.4170 | 0.1030 | 5.17E-05 |
| rs10459574 | -0.4195 | 0.1025 | 4.30E-05 |
| rs10499014 | -0.4124 | 0.1028 | 6.07E-05 |
| rs10745785 | -0.4095 | 0.1028 | 6.75E-05 |
| rs10754132 | -0.4101 | 0.1029 | 6.67E-05 |
| rs10756714 | -0.4009 | 0.1029 | 9.74E-05 |
| rs10756798 | -0.4230 | 0.1026 | 3.72E-05 |
| rs10766077 | -0.4179 | 0.1025 | 4.57E-05 |
| rs10774018 | -0.4120 | 0.1029 | 6.23E-05 |
| rs1078455 | -0.4189 | 0.1026 | 4.42E-05 |
| rs10938398 | -0.4222 | 0.1030 | 4.17E-05 |
| rs10954772 | -0.4175 | 0.1027 | 4.80E-05 |
| rs10999460 | -0.4057 | 0.1026 | 7.61E-05 |
| rs11012732 | -0.3940 | 0.1025 | 1.21E-04 |
| rs1101563 | -0.4055 | 0.1026 | 7.71E-05 |
| rs11030119 | -0.4057 | 0.1031 | 8.33E-05 |
| rs11042030 | -0.4135 | 0.1029 | 5.86E-05 |
| rs11078687 | -0.3938 | 0.1006 | 9.12E-05 |
| rs11099020 | -0.4148 | 0.1029 | 5.50E-05 |
| rs11113445 | -0.4147 | 0.1029 | 5.55E-05 |
| rs11150745 | -0.4106 | 0.1029 | 6.57E-05 |
| rs11205277 | -0.4267 | 0.1026 | 3.22E-05 |
| rs11205617 | -0.4136 | 0.1029 | 5.85E-05 |
| rs1127100 | -0.4107 | 0.1028 | 6.50E-05 |
| rs1128249 | -0.4079 | 0.1030 | 7.47E-05 |
| rs113019802 | -0.4042 | 0.1025 | 8.00E-05 |
| rs113230003 | -0.4121 | 0.1029 | 6.22E-05 |
| rs113603865 | -0.4118 | 0.1029 | 6.25E-05 |
| rs114712833 | -0.4145 | 0.1028 | 5.51E-05 |
| rs11642015 | -0.4092 | 0.1051 | 9.87E-05 |
| rs11782341 | -0.4206 | 0.1025 | 4.03E-05 |
| rs11786089 | -0.4100 | 0.1028 | 6.67E-05 |
| rs11795066 | -0.4103 | 0.1028 | 6.62E-05 |
| rs118136827 | -0.4106 | 0.1028 | 6.46E-05 |
| rs11839227 | -0.4062 | 0.1025 | 7.44E-05 |
| rs11866219 | -0.4170 | 0.1031 | 5.22E-05 |
| rs12001437 | -0.4059 | 0.1024 | 7.43E-05 |
| rs12037698 | -0.4096 | 0.1028 | 6.75E-05 |

| | | | |
|---|---|---|---|
| rs12049202 | -0.4205 | 0.1025 | 4.10E-05 |
| rs12144626 | -0.4219 | 0.1025 | 3.84E-05 |
| rs12339822 | -0.4054 | 0.1027 | 7.86E-05 |
| rs12367809 | -0.4201 | 0.1031 | 4.58E-05 |
| rs12619178 | -0.4139 | 0.1029 | 5.72E-05 |
| rs1266921 | -0.4148 | 0.1028 | 5.50E-05 |
| rs12679106 | -0.4162 | 0.1029 | 5.21E-05 |
| rs12885458 | -0.4009 | 0.1017 | 8.07E-05 |
| rs12920259 | -0.3874 | 0.0995 | 9.93E-05 |
| rs1293446 | -0.4158 | 0.1028 | 5.24E-05 |
| rs12997625 | -0.4205 | 0.1024 | 4.00E-05 |
| rs13056506 | -0.4173 | 0.1028 | 4.94E-05 |
| rs13062093 | -0.4134 | 0.1029 | 5.87E-05 |
| rs13107325 | -0.3595 | 0.1001 | 3.27E-04 |
| rs13174863 | -0.4104 | 0.1029 | 6.64E-05 |
| rs1322842 | -0.4139 | 0.1028 | 5.70E-05 |
| rs13420048 | -0.4168 | 0.1027 | 4.93E-05 |
| rs1441264 | -0.4098 | 0.1029 | 6.87E-05 |
| rs1446585 | -0.4445 | 0.0986 | 6.51E-06 |
| rs1503526 | -0.4139 | 0.1029 | 5.78E-05 |
| rs1514173 | -0.4112 | 0.1028 | 6.38E-05 |
| rs1516725 | -0.4098 | 0.1029 | 6.79E-05 |
| rs153664 | -0.4135 | 0.1028 | 5.82E-05 |
| rs1568489 | -0.4201 | 0.1024 | 4.12E-05 |
| rs1652376 | -0.4056 | 0.1026 | 7.77E-05 |
| rs16951304 | -0.4141 | 0.1031 | 5.89E-05 |
| rs16953563 | -0.4138 | 0.1028 | 5.73E-05 |
| rs17024393 | -0.4143 | 0.1031 | 5.84E-05 |
| rs1724557 | -0.4145 | 0.1028 | 5.54E-05 |
| rs17342242 | -0.4134 | 0.1028 | 5.83E-05 |
| rs1786978 | -0.4143 | 0.1028 | 5.59E-05 |
| rs1881505 | -0.4118 | 0.1027 | 6.05E-05 |
| rs1884902 | -0.4184 | 0.1028 | 4.75E-05 |
| rs1927626 | -0.4095 | 0.1029 | 6.87E-05 |
| rs1928496 | -0.4117 | 0.1029 | 6.25E-05 |
| rs2002023 | -0.4128 | 0.1029 | 6.00E-05 |
| rs2020942 | -0.4177 | 0.1027 | 4.76E-05 |
| rs2035936 | -0.4089 | 0.1028 | 6.90E-05 |
| rs2043016 | -0.4095 | 0.1028 | 6.78E-05 |
| rs2052607 | -0.3994 | 0.1021 | 9.18E-05 |
| rs2124499 | -0.4155 | 0.1028 | 5.31E-05 |
| rs2172131 | -0.4071 | 0.1027 | 7.42E-05 |
| rs217672 | -0.4158 | 0.1027 | 5.16E-05 |
| rs2182633 | -0.4172 | 0.1027 | 4.82E-05 |
| rs2239647 | -0.4172 | 0.1029 | 4.99E-05 |

| | | | |
|---|---|---|---|
| rs2279574 | -0.4260 | 0.1026 | 3.31E-05 |
| rs2281819 | -0.4065 | 0.1025 | 7.35E-05 |
| rs2306593 | -0.4214 | 0.1025 | 3.92E-05 |
| rs2307111 | -0.4109 | 0.1030 | 6.63E-05 |
| rs2318543 | -0.4138 | 0.1029 | 5.76E-05 |
| rs2356505 | -0.4184 | 0.1026 | 4.55E-05 |
| rs2370982 | -0.4143 | 0.1030 | 5.78E-05 |
| rs2374947 | -0.4196 | 0.1025 | 4.29E-05 |
| rs2384054 | -0.4136 | 0.1037 | 6.67E-05 |
| rs2398861 | -0.4054 | 0.1024 | 7.52E-05 |
| rs240461 | -0.4126 | 0.1028 | 6.04E-05 |
| rs2439823 | -0.4151 | 0.1030 | 5.53E-05 |
| rs245775 | -0.4168 | 0.1029 | 5.09E-05 |
| rs2470531 | -0.4143 | 0.1028 | 5.61E-05 |
| rs2494196 | -0.4125 | 0.1030 | 6.25E-05 |
| rs2500406 | -0.4007 | 0.1018 | 8.32E-05 |
| rs2515963 | -0.4135 | 0.1029 | 5.83E-05 |
| rs2569881 | -0.4177 | 0.1027 | 4.72E-05 |
| rs2606228 | -0.4182 | 0.1026 | 4.61E-05 |
| rs2623259 | -0.4173 | 0.1027 | 4.83E-05 |
| rs2678204 | -0.4033 | 0.1027 | 8.60E-05 |
| rs273505 | -0.4142 | 0.1030 | 5.79E-05 |
| rs2815764 | -0.4131 | 0.1030 | 6.03E-05 |
| rs28418580 | -0.4028 | 0.1023 | 8.23E-05 |
| rs28483178 | -0.4120 | 0.1031 | 6.39E-05 |
| rs2855818 | -0.4001 | 0.1022 | 9.08E-05 |
| rs2861685 | -0.4083 | 0.1027 | 7.04E-05 |
| rs2914231 | -0.4196 | 0.1026 | 4.30E-05 |
| rs2943650 | -0.4117 | 0.1030 | 6.40E-05 |
| rs2954021 | -0.4128 | 0.1029 | 6.02E-05 |
| rs301806 | -0.4044 | 0.1024 | 7.81E-05 |
| rs3134404 | -0.4205 | 0.1024 | 4.01E-05 |
| rs34045894 | -0.4178 | 0.1026 | 4.66E-05 |
| rs34292685 | -0.4116 | 0.1029 | 6.28E-05 |
| rs34331990 | -0.4069 | 0.1026 | 7.28E-05 |
| rs34338229 | -0.4091 | 0.1027 | 6.85E-05 |
| rs34417222 | -0.4202 | 0.1026 | 4.24E-05 |
| rs34483452 | -0.4215 | 0.1029 | 4.18E-05 |
| rs34580448 | -0.4150 | 0.1029 | 5.49E-05 |
| rs34769775 | -0.4113 | 0.1029 | 6.37E-05 |
| rs34811474 | -0.4121 | 0.1028 | 6.12E-05 |
| rs34882821 | -0.4080 | 0.1027 | 7.08E-05 |
| rs35851183 | -0.4071 | 0.1027 | 7.37E-05 |
| rs3734554 | -0.4168 | 0.1029 | 5.10E-05 |
| rs3803286 | -0.4183 | 0.1027 | 4.64E-05 |

| | | | |
|---|---|---|---|
| rs3826408 | -0.4238 | 0.1024 | 3.52E-05 |
| rs3856595 | -0.4107 | 0.1028 | 6.48E-05 |
| rs3923501 | -0.4124 | 0.1029 | 6.10E-05 |
| rs40067 | -0.4124 | 0.1029 | 6.15E-05 |
| rs4073582 | -0.4003 | 0.1017 | 8.25E-05 |
| rs429358 | -0.4169 | 0.1028 | 5.05E-05 |
| rs4402589 | -0.4006 | 0.1026 | 9.43E-05 |
| rs4450871 | -0.4105 | 0.1026 | 6.32E-05 |
| rs4477562 | -0.4120 | 0.1029 | 6.23E-05 |
| rs4519214 | -0.4141 | 0.1028 | 5.66E-05 |
| rs4547132 | -0.4133 | 0.1028 | 5.85E-05 |
| rs4549080 | -0.4157 | 0.1028 | 5.29E-05 |
| rs4549685 | -0.4106 | 0.1029 | 6.62E-05 |
| rs4643373 | -0.4141 | 0.1030 | 5.76E-05 |
| rs4727630 | -0.4173 | 0.1027 | 4.88E-05 |
| rs4752182 | -0.4194 | 0.1025 | 4.24E-05 |
| rs4764949 | -0.4079 | 0.1027 | 7.07E-05 |
| rs4790841 | -0.4149 | 0.1030 | 5.67E-05 |
| rs4876611 | -0.4083 | 0.1029 | 7.27E-05 |
| rs538656 | -0.4035 | 0.1032 | 9.27E-05 |
| rs541577 | -0.4168 | 0.1027 | 4.96E-05 |
| rs543874 | -0.4365 | 0.1034 | 2.44E-05 |
| rs55637757 | -0.4106 | 0.1027 | 6.40E-05 |
| rs56288810 | -0.4182 | 0.1026 | 4.60E-05 |
| rs56399737 | -0.4099 | 0.1028 | 6.71E-05 |
| rs57221746 | -0.4114 | 0.1028 | 6.30E-05 |
| rs57636386 | -0.4104 | 0.1030 | 6.76E-05 |
| rs57800857 | -0.4117 | 0.1029 | 6.32E-05 |
| rs58120873 | -0.4097 | 0.1027 | 6.67E-05 |
| rs59066241 | -0.4150 | 0.1028 | 5.42E-05 |
| rs59104534 | -0.4137 | 0.1028 | 5.75E-05 |
| rs60764613 | -0.4079 | 0.1027 | 7.14E-05 |
| rs61037750 | -0.4175 | 0.1030 | 5.03E-05 |
| rs61787785 | -0.4116 | 0.1028 | 6.29E-05 |
| rs61813324 | -0.4097 | 0.1027 | 6.58E-05 |
| rs61903695 | -0.4137 | 0.1028 | 5.76E-05 |
| rs62017600 | -0.4249 | 0.1021 | 3.18E-05 |
| rs62037365 | -0.3948 | 0.1029 | 1.25E-04 |
| rs62178054 | -0.4134 | 0.1029 | 5.86E-05 |
| rs62244189 | -0.4171 | 0.1026 | 4.81E-05 |
| rs62261725 | -0.4047 | 0.1026 | 8.01E-05 |
| rs62407779 | -0.4200 | 0.1031 | 4.59E-05 |
| rs62477684 | -0.4171 | 0.1029 | 5.05E-05 |
| rs635634 | -0.4182 | 0.1029 | 4.87E-05 |
| rs6545714 | -0.4186 | 0.1029 | 4.73E-05 |

| | | | |
|---|---|---|---|
| rs6548237 | -0.4252 | 0.1030 | 3.64E-05 |
| rs6561937 | -0.4097 | 0.1028 | 6.74E-05 |
| rs6575340 | -0.4121 | 0.1030 | 6.25E-05 |
| rs6601527 | -0.4221 | 0.1023 | 3.71E-05 |
| rs66679256 | -0.4115 | 0.1029 | 6.40E-05 |
| rs66833742 | -0.4003 | 0.1022 | 8.99E-05 |
| rs6754974 | -0.4177 | 0.1026 | 4.69E-05 |
| rs6861649 | -0.4070 | 0.1027 | 7.37E-05 |
| rs6927268 | -0.4171 | 0.1027 | 4.87E-05 |
| rs705158 | -0.4100 | 0.1028 | 6.65E-05 |
| rs7124681 | -0.4164 | 0.1035 | 5.73E-05 |
| rs7133378 | -0.4167 | 0.1030 | 5.26E-05 |
| rs7201895 | -0.4086 | 0.1028 | 7.02E-05 |
| rs7246182 | -0.4135 | 0.1028 | 5.80E-05 |
| rs7259070 | -0.4109 | 0.1028 | 6.38E-05 |
| rs73213484 | -0.4099 | 0.1028 | 6.67E-05 |
| rs73307079 | -0.4147 | 0.1028 | 5.50E-05 |
| rs745249 | -0.4151 | 0.1028 | 5.42E-05 |
| rs7498044 | -0.4136 | 0.1028 | 5.77E-05 |
| rs750090 | -0.4142 | 0.1028 | 5.63E-05 |
| rs75499503 | -0.4079 | 0.1029 | 7.42E-05 |
| rs7580600 | -0.4066 | 0.1028 | 7.65E-05 |
| rs7591494 | -0.4172 | 0.1028 | 4.96E-05 |
| rs76040172 | -0.4024 | 0.1023 | 8.44E-05 |
| rs7609467 | -0.4125 | 0.1028 | 6.05E-05 |
| rs7613261 | -0.4158 | 0.1029 | 5.33E-05 |
| rs764729 | -0.4204 | 0.1024 | 4.06E-05 |
| rs7649970 | -0.4148 | 0.1030 | 5.61E-05 |
| rs7719067 | -0.4120 | 0.1029 | 6.20E-05 |
| rs7739003 | -0.4182 | 0.1031 | 4.99E-05 |
| rs7755574 | -0.4123 | 0.1029 | 6.12E-05 |
| rs778094 | -0.4110 | 0.1028 | 6.40E-05 |
| rs78801969 | -0.4157 | 0.1027 | 5.17E-05 |
| rs7930275 | -0.4057 | 0.1026 | 7.61E-05 |
| rs79535757 | -0.4087 | 0.1026 | 6.78E-05 |
| rs7972728 | -0.4212 | 0.1026 | 4.02E-05 |
| rs7987928 | -0.4092 | 0.1028 | 6.85E-05 |
| rs80135947 | -0.3976 | 0.1023 | 1.01E-04 |
| rs8042404 | -0.4147 | 0.1028 | 5.51E-05 |
| rs8078135 | -0.4136 | 0.1028 | 5.78E-05 |
| rs8087074 | -0.4178 | 0.1027 | 4.74E-05 |
| rs8134638 | -0.4128 | 0.1028 | 5.98E-05 |
| rs815163 | -0.4169 | 0.1028 | 4.98E-05 |
| rs8192675 | -0.4110 | 0.1029 | 6.45E-05 |
| rs846545 | -0.4131 | 0.1029 | 5.93E-05 |

| | | | |
|---|---|---|---|
| rs879620 | -0.4006 | 0.1026 | 9.50E-05 |
| rs9320823 | -0.4087 | 0.1029 | 7.17E-05 |
| rs9342196 | -0.4081 | 0.1027 | 7.08E-05 |
| rs935166 | -0.4173 | 0.1027 | 4.86E-05 |
| rs9522183 | -0.4152 | 0.1028 | 5.38E-05 |
| rs9522279 | -0.4178 | 0.1027 | 4.78E-05 |
| rs9584855 | -0.4174 | 0.1028 | 4.87E-05 |
| rs9641499 | -0.4129 | 0.1029 | 6.00E-05 |
| rs9688977 | -0.4138 | 0.1028 | 5.75E-05 |
| rs972283 | -0.4201 | 0.1028 | 4.38E-05 |
| rs9843653 | -0.4248 | 0.1028 | 3.57E-05 |
| rs9861440 | -0.4191 | 0.1025 | 4.35E-05 |
| All | -0.4127 | 0.1025 | 5.66E-05 |

**Table S3.** Reverse causality analyses

| Exposure | method | No. of SNPs | beta | se | p-value | Outcome |
|---|---|---|---|---|---|---|
| Chang et al. 2017 | MR Egger | 18 | 0.019 | 0.020 | 0.350 | Arm fat percentage (right) \|\| id:UKB-a:282 |
| Chang et al. 2017 | Weighted median | 18 | 0.012 | 0.007 | 0.070 | Arm fat percentage (right) \|\| id:UKB-a:282 |
| Chang et al. 2017 | Inverse variance weighted | 18 | 0.015 | 0.008 | 0.066 | Arm fat percentage (right) \|\| id:UKB-a:282 |
| Chang et al. 2017 | MR Egger | 18 | 0.026 | 0.021 | 0.233 | Arm fat percentage (left) \|\| id:UKB-a:286 |
| Chang et al. 2017 | Weighted median | 18 | 0.016 | 0.007 | 0.015 | Arm fat percentage (left) \|\| id:UKB-a:286 |
| Chang et al. 2017 | Inverse variance weighted | 18 | 0.015 | 0.008 | 0.062 | Arm fat percentage (left) \|\| id:UKB-a:286 |
| Chang et al. 2017 | MR Egger | 18 | 0.035 | 0.029 | 0.247 | Arm fat mass (right) \|\| id:UKB-a:283 |
| Chang et al. 2017 | Weighted median | 18 | 0.016 | 0.008 | 0.053 | Arm fat mass (right) \|\| id:UKB-a:283 |
| Chang et al. 2017 | Inverse variance weighted | 18 | 0.018 | 0.012 | 0.115 | Arm fat mass (right) \|\| id:UKB-a:283 |
| Chang et al. 2017 | MR Egger | 18 | 0.008 | 0.016 | 0.626 | Leg fat percentage (right) \|\| id:UKB-a:274 |
| Chang et al. 2017 | Weighted median | 18 | 0.008 | 0.005 | 0.141 | Leg fat percentage (right) \|\| id:UKB-a:274 |
| Chang et al. 2017 | Inverse variance weighted | 18 | 0.012 | 0.006 | 0.051 | Leg fat percentage (right) \|\| id:UKB-a:274 |
| Chang et al. 2017 | MR Egger | 18 | 0.045 | 0.029 | 0.142 | Whole body fat mass \|\| id:UKB-a:265 |
| Chang et al. 2017 | Weighted median | 18 | 0.028 | 0.008 | 0.001 | Whole body fat mass \|\| id:UKB-a:265 |
| Chang et al. 2017 | Inverse variance weighted | 18 | 0.022 | 0.012 | 0.060 | Whole body fat mass \|\| id:UKB-a:265 |
| Chang et al. 2017 | MR Egger | 18 | 0.037 | 0.030 | 0.235 | Arm fat mass (left) \|\| id:UKB-a:287 |
| Chang et al. 2017 | Weighted median | 18 | 0.021 | 0.008 | 0.011 | Arm fat mass (left) \|\| id:UKB-a:287 |
| Chang et al. 2017 | Inverse variance weighted | 18 | 0.019 | 0.012 | 0.104 | Arm fat mass (left) \|\| id:UKB-a:287 |
| Chang et al. 2017 | MR Egger | 18 | 0.058 | 0.030 | 0.071 | Trunk fat mass \|\| id:UKB-a:291 |
| Chang et al. 2017 | Weighted median | 18 | 0.038 | 0.009 | 0.000 | Trunk fat mass \|\| id:UKB-a:291 |
| Chang et al. 2017 | Inverse variance weighted | 18 | 0.027 | 0.012 | 0.028 | Trunk fat mass \|\| id:UKB-a:291 |
| Chang et al. 2017 | MR Egger | 18 | -0.008 | 0.013 | 0.545 | Time spent watching television (TV) \|\| id:UKB-b:5192 |
| Chang et al. 2017 | Weighted median | 18 | -0.010 | 0.005 | 0.052 | Time spent watching television (TV) \|\| id:UKB-b:5192 |
| Chang et al. 2017 | Inverse variance weighted | 18 | -0.005 | 0.005 | 0.306 | Time spent watching television (TV) \|\| id:UKB-b:5192 |
| Chang et al. 2017 | MR Egger | 18 | 0.049 | 0.026 | 0.078 | Trunk fat percentage \|\| id:UKB-a:290 |
| Chang et al. 2017 | Weighted median | 18 | 0.036 | 0.010 | 0.000 | Trunk fat percentage \|\| id:UKB-a:290 |
| Chang et al. 2017 | Inverse variance weighted | 18 | 0.026 | 0.010 | 0.014 | Trunk fat percentage \|\| id:UKB-a:290 |
| Chang et al. 2017 | MR Egger | 18 | 0.081 | 0.034 | 0.030 | Forced vital capacity (FVC) Best measure \|\| id:UKB-a:232 |
| Chang et al. 2017 | Weighted median | 18 | 0.009 | 0.009 | 0.355 | Forced vital capacity (FVC) Best measure \|\| id:UKB-a:232 |
| Chang et al. 2017 | Inverse variance weighted | 18 | 0.025 | 0.015 | 0.094 | Forced vital capacity (FVC) Best measure \|\| id:UKB-a:232 |
| Chang et al. 2017 | MR Egger | 18 | -0.010 | 0.018 | 0.584 | Tea intake \|\| id:UKB-b:6066 |
| Chang et al. 2017 | Weighted median | 18 | -0.003 | 0.007 | 0.689 | Tea intake \|\| id:UKB-b:6066 |

| | | | | | | |
|---|---|---|---|---|---|---|
| Chang et al. 2017 | Inverse variance weighted | 18 | -0.008 | 0.007 | 0.257 | Tea intake \|\| id:UKB-b:6066 |
| Chang et al. 2017 | MR Egger | 18 | -0.006 | 0.032 | 0.859 | Impedance of leg (right) \|\| id:UKB-a:270 |
| Chang et al. 2017 | Weighted median | 18 | 0.005 | 0.010 | 0.636 | Impedance of leg (right) \|\| id:UKB-a:270 |
| Chang et al. 2017 | Inverse variance weighted | 18 | 0.004 | 0.013 | 0.752 | Impedance of leg (right) \|\| id:UKB-a:270 |

id, specific code attributed to each trait by MR Base; se, standard error; No. of SNPs, number of SNPs

**Table S4.** Mendelian randomization analyses of exposures significantly associated to PD at IVW uncorrected p-value < 0.05 vs Nalls et al., 2019 (only clinically diagnosed cases)

| Exposure | no. of SNPs | beta | se | p-value | FDR |
|---|---|---|---|---|---|
| Hip circumference \|\| id:UKB-a:388 | 278 | -0.2122 | 0.072 | 0.003 | 0.7529 |
| Frequency of tiredness / lethargy in last 2 weeks \|\| id:UKB-b:929 | 39 | -0.9405 | 0.362 | 0.009 | 0.7529 |
| Hair/balding pattern: Pattern 3 \|\| id:UKB-a:302 | 37 | 1.0982 | 0.442 | 0.013 | 0.7529 |
| Fed-up feelings \|\| id:UKB-a:49 | 31 | -2.3210 | 0.956 | 0.015 | 0.7529 |
| Leg fat mass (left) \|\| id:UKB-a:279 | 276 | -0.2282 | 0.097 | 0.018 | 0.7529 |
| Frequency of friend/family visits \|\| id:UKB-b:5379 | 20 | -0.9743 | 0.420 | 0.020 | 0.7529 |
| Trunk fat mass \|\| id:UKB-a:291 | 281 | -0.1673 | 0.073 | 0.021 | 0.7529 |
| Non-cancer illness code self-reported: hypertension \|\| id:UKB-a:61 | 153 | -0.5528 | 0.242 | 0.022 | 0.7529 |
| Type 2 diabetes \|\| id:25 | 25 | 0.0767 | 0.035 | 0.026 | 0.7529 |
| Forced expiratory volume in 1-second (FEV1) Best measure \|\| id:UKB-a:231 | 141 | 0.3129 | 0.144 | 0.029 | 0.7529 |
| Guilty feelings \|\| id:UKB-a:240 | 17 | -3.2359 | 1.490 | 0.030 | 0.7529 |
| Whole body fat mass \|\| id:UKB-a:265 | 276 | -0.1587 | 0.073 | 0.031 | 0.7529 |
| Overall health rating \|\| id:UKB-a:251 | 54 | -1.3154 | 0.611 | 0.031 | 0.7529 |
| Leg fat mass (right) \|\| id:UKB-a:275 | 276 | -0.2070 | 0.096 | 0.032 | 0.7529 |
| Treatment/medication code: levothyroxine sodium \|\| id:UKB-a:190 | 54 | -1.6648 | 0.792 | 0.035 | 0.7529 |
| Non-cancer illness code self-reported: migraine \|\| id:UKB-a:78 | 13 | -4.0746 | 1.977 | 0.039 | 0.7529 |
| Cancer code self-reported: prostate cancer \|\| id:UKB-a:57 | 12 | 8.8486 | 4.301 | 0.040 | 0.7529 |
| Fasting insulin \|\| id:779 | 14 | 0.8836 | 0.433 | 0.041 | 0.7529 |
| Mouth/teeth dental problems: Mouth ulcers \|\| id:UKB-a:427 | 29 | -1.1501 | 0.565 | 0.042 | 0.7529 |
| Trunk fat percentage \|\| id:UKB-a:290 | 233 | -0.1839 | 0.091 | 0.042 | 0.7529 |
| Vascular/heart problems diagnosed by doctor: None of the above \|\| id:UKB-a:435 | 142 | 0.4590 | 0.229 | 0.045 | 0.7529 |
| Medication for pain relief constipation heartburn: None of the above \|\| id:UKB-a:453 | 15 | 3.0620 | 1.536 | 0.046 | 0.7529 |
| Leg fat percentage (right) \|\| id:UKB-a:274 | 245 | -0.2654 | 0.135 | 0.050 | 0.7529 |
| Sleeplessness / insomnia \|\| id:UKB-b:3957 | 39 | 0.7305 | 0.372 | 0.050 | 0.7529 |

id, specific code attributed to each trait by MR Base; se, standard error; No. of SNPs, number of SNPs; MR, Mendelian randomization; NA: not applicable due to limited no. of SNPs

**Table S5.** Mendelian randomization analyses of exposures significantly associated to PD at IVW p-value < 0.05 vs Nalls et al., 2014

| **Exposure** | **no. of SNPs** | **beta** | **se** | **p-value** | **FDR** |
|---|---|---|---|---|---|
| Arm fat percentage (right) \|\| id:UKB-a:282 | 213 | -0.447 | 0.099 | 5.91E-06 | 0.0024 |
| Trunk fat percentage \|\| id:UKB-a:290 | 215 | -0.352 | 0.087 | 5.30E-05 | 0.0082 |
| Whole body fat mass \|\| id:UKB-a:265 | 257 | -0.283 | 0.072 | 7.46E-05 | 0.0082 |
| Arm fat percentage (left) \|\| id:UKB-a:286 | 232 | -0.398 | 0.101 | 8.17E-05 | 0.0082 |
| Arm fat mass (right) \|\| id:UKB-a:283 | 247 | -0.282 | 0.073 | 1.17E-04 | 0.0093 |
| Arm fat mass (left) \|\| id:UKB-a:287 | 247 | -0.280 | 0.074 | 1.68E-04 | 0.0096 |
| Trunk fat mass \|\| id:UKB-a:291 | 256 | -0.266 | 0.071 | 1.84E-04 | 0.0096 |
| Leg fat percentage (right) \|\| id:UKB-a:274 | 228 | -0.485 | 0.130 | 1.91E-04 | 0.0096 |
| Leg fat percentage (left) \|\| id:UKB-a:278 | 226 | -0.431 | 0.133 | 1.20E-03 | 0.0471 |
| Leg fat mass (right) \|\| id:UKB-a:275 | 257 | -0.295 | 0.091 | 1.21E-03 | 0.0471 |
| Weight \|\| id:UKB-a:249 | 303 | -0.229 | 0.071 | 1.29E-03 | 0.0488 |

id, specific code attributed to each trait by MR Base; se, standard error; No. of SNPs, number of SNPs; MR, Mendelian randomization; NA: not applicable due to limited no. of SNPs

**Mendelian Randomization assumptions**

1) The instrumental variables are associated with the exposure (measured using $R^2$ and a first-stage F statistic)

2) The instrument variables are not associated with confounders of the exposure-outcome association

3) The instrument variables are not associated with the outcome via any alternative path other than one that passes through the exposure (our risk factor of interest) (Didelez and Sheehan 2007). Violation of the third assumption can arise from horizontal pleiotropy or from heterogeneity. Pleiotropy occurs when the SNPs are directly related to the outcome itself or affect certain characteristics that are risk factors for the outcome independent of the exposure. Heterogeneity refers to the situations where SNPs are associated with population substructure or with many correlated traits.

**IPDGC consortium members and affiliations:**

**United Kingdom:**
Alastair J Noyce (Preventive Neurology Unit, Wolfson Institute of Preventive Medicine, QMUL, London, UK and Department of Molecular Neuroscience, UCL, London, UK),
Rauan Kaiyrzhanov (Department of Molecular Neuroscience, UCL Institute of Neurology, London, UK),
Ben Middlehurst (Institute of Translational Medicine, University of Liverpool, Liverpool, UK),
Demis A Kia (UCL Genetics Institute; and Department of Molecular Neuroscience, UCL Institute of Neurology, London, UK),
Manuela Tan (Department of Clinical Neuroscience, University College London, London, UK),
Henry Houlden (Department of Molecular Neuroscience, UCL Institute of Neurology, London, UK),
Huw R Morris (Department of Clinical Neuroscience, University College London, London, UK),
Helene Plun-Favreau (Department of Molecular Neuroscience, UCL Institute of Neurology, London, UK),
Peter Holmans (Biostatistics & Bioinformatics Unit, Institute of Psychological Medicine and Clinical Neuroscience, MRC Centre for Neuropsychiatric Genetics & Genomics, Cardiff, UK),
John Hardy (Department of Molecular Neuroscience, UCL Institute of Neurology, London, UK),
Daniah Trabzuni (Department of Molecular Neuroscience, UCL Institute of Neurology, London, UK; Department of Genetics, King Faisal Specialist Hospital and Research Centre, Riyadh, 11211 Saudi Arabia),
Jose Bras (UK Dementia Research Institute at UCL and Department of Molecular Neuroscience, UCL Institute of Neurology, London, UK),
John Quinn PhD (Institute of Translational Medicine, University of Liverpool, Liverpool, UK),
Kin Y Mok (Department of Molecular Neuroscience, UCL Institute of Neurology, London, UK),
Kerri J. Kinghorn (Institute of Healthy Ageing, University College London, London, UK),
Kimberley Billingsley (Institute of Translational Medicine, University of Liverpool, Liverpool, UK),
Nicholas W Wood (UCL Genetics Institute; and Department of Molecular Neuroscience, UCL Institute of Neurology, London, UK),
Patrick Lewis (University of Reading, Reading, UK),
Sebastian Schreglmann (Department of Molecular Neuroscience, UCL Institute of Neurology, London, UK),
Rita Guerreiro (UK Dementia Research Institute at UCL and Department of Molecular Neuroscience, UCL Institute of Neurology, London, UK),
Ruth Lovering (University College London, London, UK),
Lea R'Bibo (Department of Molecular Neuroscience, UCL Institute of Neurology, London, UK),
Claudia Manzoni (University of Reading, Reading, UK),
Mie Rizig (Department of Molecular Neuroscience, UCL Institute of Neurology, London, UK),
Mina Ryten (Department of Molecular Neuroscience, UCL Institute of Neurology, London, UK),
Sebastian Guelfi (Department of Molecular Neuroscience, UCL Institute of Neurology, London, UK),
Valentina Escott-Price (MRC Centre for Neuropsychiatric Genetics and Genomics, Cardiff University School of Medicine, Cardiff, UK),
Viorica Chelban (Department of Molecular Neuroscience, UCL Institute of Neurology, London, UK),
Thomas Foltynie (UCL Institute of Neurology, London, UK),
Nigel Williams (MRC Centre for Neuropsychiatric Genetics and Genomics, Cardiff, UK), Karen E. Morrison (Faculty of Medicine, University of Southampton, UK), Carl Clarke (University of Birmingham, Birmingham, UK and Sandwell and West Birmingham Hospitals NHS Trust, Birmingham, UK).

**France:**
Alexis Brice (Institut du Cerveau et de la Moelle épinière, ICM, Inserm U 1127, CNRS, UMR 7225, Sorbonne Universités, UPMC University Paris 06, UMR S 1127, AP-HP, Pitié-Salpêtrière Hospital, Paris, France),
Fabrice Danjou (Institut du Cerveau et de la Moelle épinière, ICM, Inserm U 1127, CNRS, UMR 7225, Sorbonne Universités, UPMC University Paris 06, UMR S 1127, AP-HP, Pitié-Salpêtrière Hospital, Paris, France),
Suzanne Lesage (Institut du Cerveau et de la Moelle épinière, ICM, Inserm U 1127, CNRS, UMR 7225, Sorbonne Universités, UPMC University Paris 06, UMR S 1127, AP-HP, Pitié-Salpêtrière Hospital, Paris, France),

Jean-Christophe Corvol (Institut du Cerveau et de la Moelle épinière, ICM, Inserm U 1127, CNRS, UMR 7225, Sorbonne Universités, UPMC University Paris 06, UMR S 1127, Centre d'Investigation Clinique Pitié Neurosciences CIC-1422, AP-HP, Pitié-Salpêtrière Hospital, Paris, France),
Maria Martinez (INSERM UMR 1220; and Paul Sabatier University, Toulouse, France),

**Germany:**
Claudia Schulte (Department for Neurodegenerative Diseases, Hertie Institute for Clinical Brain Research, University of Tübingen, and DZNE, German Center for Neurodegenerative Diseases, Tübingen, Germany),
Kathrin Brockmann (Department for Neurodegenerative Diseases, Hertie Institute for Clinical Brain Research, University of Tübingen, and DZNE, German Center for Neurodegenerative Diseases, Tübingen, Germany),
Javier Simón-Sánchez (Department for Neurodegenerative Diseases, Hertie Institute for Clinical Brain Research, University of Tübingen, and DZNE, German Center for Neurodegenerative Diseases, Tübingen, Germany),
Peter Heutink (DZNE, German Center for Neurodegenerative Diseases and Department for Neurodegenerative Diseases, Hertie Institute for Clinical Brain Research, University of Tübingen, Tübingen, Germany),
Patrizia Rizzu (DZNE, German Center for Neurodegenerative Diseases),
Manu Sharma (Centre for Genetic Epidemiology, Institute for Clinical Epidemiology and Applied Biometry, University of Tubingen, Germany),
Thomas Gasser (Department for Neurodegenerative Diseases, Hertie Institute for Clinical Brain Research, and DZNE, German Center for Neurodegenerative Diseases, Tübingen, Germany),

**United States of America:**
Aude Nicolas (Laboratory of Neurogenetics, National Institute on Aging, Bethesda, MD, USA),
Mark R Cookson (Laboratory of Neurogenetics, National Institute on Aging, Bethesda, USA),
Sara Bandres-Ciga (Laboratory of Neurogenetics, National Institute on Aging, Bethesda, MD, USA),
Cornelis Blauwendraat (National Institute on Aging and National Institute of Neurological Disorders and Stroke, USA),
David W. Craig (Department of Translational Genomics, Keck School of Medicine, University of Southern California, Los Angeles, USA),
Faraz Faghri (Laboratory of Neurogenetics, National Institute on Aging, Bethesda, USA; Department of Computer Science, University of Illinois at Urbana-Champaign, Urbana, IL, USA),
J Raphael Gibbs (Laboratory of Neurogenetics, National Institute on Aging, National Institutes of Health, Bethesda, MD, USA),
Dena G. Hernandez (Laboratory of Neurogenetics, National Institute on Aging, Bethesda, MD, USA),
Kendall Van Keuren-Jensen (Neurogenomics Division, TGen, Phoenix, AZ USA),
Joshua M. Shulman (Departments of Neurology, Neuroscience, and Molecular & Human Genetics, Baylor College of Medicine, Houston, Texas, USA; Jan and Dan Duncan Neurological Research Institute, Texas Children's Hospital, Houston, Texas, USA),
Hirotaka Iwaki (Laboratory of Neurogenetics, National Institute on Aging, Bethesda, MD, USA),
Hampton L. Leonard (Laboratory of Neurogenetics, National Institute on Aging, Bethesda, MD, USA),
Mike A. Nalls (Laboratory of Neurogenetics, National Institute on Aging, Bethesda, USA; CEO/Consultant Data Tecnica International, Glen Echo, MD, USA),
Laurie Robak (Baylor College of Medicine, Houston, Texas, USA),
Steven Lubbe (Ken and Ruth Davee Department of Neurology, Northwestern University Feinberg School of Medicine, Chicago, IL, USA),
Steven Finkbeiner (Departments of Neurology and Physiology, University of California, San Francisco; Gladstone Institute of Neurological Disease; Taube/Koret Center for Neurodegenerative Disease Research, San Francisco, CA, USA),
Niccolo E. Mencacci (Northwestern University Feinberg School of Medicine, Chicago, IL, USA),
Codrin Lungu (National Institutes of Health Division of Clinical Research, NINDS, National Institutes of Health, Bethesda, MD, USA),
Andrew B Singleton (Laboratory of Neurogenetics, National Institute on Aging, Bethesda, MD, USA),
Sonja W. Scholz (Neurodegenerative Diseases Research Unit, National Institute of Neurological Disorders and Stroke, Bethesda, MD, USA),

Xylena Reed (Laboratory of Neurogenetics, National Institute on Aging, Bethesda, MD, USA).
Roy N. Alcalay (Department of Neurology, College of Physicians and Surgeons, Columbia University Medical Center, New York, NY, USA, Taub Institute for Research on Alzheimer's Disease and the Aging Brain, College of Physicians and Surgeons, Columbia University Medical Center, New York, NY, USA)

**Canada:**
Ziv Gan-Or (Montreal Neurological Institute and Hospital, Department of Neurology & Neurosurgery, Department of Human Genetics, McGill University, Montréal, QC, H3A 0G4, Canada),
Guy A. Rouleau (Montreal Neurological Institute and Hospital, Department of Neurology & Neurosurgery, Department of Human Genetics, McGill University, Montréal, QC, H3A 0G4, Canada), Lynne Krohn (Montreal Neurological Institute and Hospital, Department of Neurology & Neurosurgery, Department of Human Genetics, McGill University, Montréal, QC, H3A 0G4, Canada)

**The Netherlands:**
Jacobus J van Hilten (Department of Neurology, Leiden University Medical Center, Leiden, Netherlands),
Johan Marinus (Department of Neurology, Leiden University Medical Center, Leiden, Netherlands)

**Spain:**
Astrid D. Adarmes-Gómez (Instituto de Biomedicina de Sevilla (IBiS), Hospital Universitario Virgen del Rocío/CSIC/Universidad de Sevilla, Seville),
Miquel Aguilar (Fundació Docència i Recerca Mútua de Terrassa and Movement Disorders Unit, Department of Neurology, University Hospital Mutua de Terrassa, Terrassa, Barcelona.),
Ignacio Alvarez (Fundació Docència i Recerca Mútua de Terrassa and Movement Disorders Unit, Department of Neurology, University Hospital Mutua de Terrassa, Terrassa, Barcelona.),
Victoria Alvarez (Hospital Universitario Central de Asturias, Oviedo),
Francisco Javier Barrero (Hospital Universitario San Cecilio de Granada, Universidad de Granada),
Jesús Alberto Bergareche Yarza (Instituto de Investigación Sanitaria Biodonostia, San Sebastián),
Inmaculada Bernal-Bernal (Instituto de Biomedicina de Sevilla (IBiS), Hospital Universitario Virgen del Rocío/CSIC/Universidad de Sevilla, Seville),
Marta Blazquez (Hospital Universitario Central de Asturias, Oviedo),
Marta Bonilla-Toribio (Instituto de Biomedicina de Sevilla (IBiS), Hospital Universitario Virgen del Rocío/CSIC/Universidad de Sevilla, Seville),
Juan A. Botía (Universidad de Murcia, Murcia),
María Teresa Boungiorno (Fundació Docència i Recerca Mútua de Terrassa and Movement Disorders Unit, Department of Neurology, University Hospital Mutua de Terrassa, Terrassa, Barcelona.)
Dolores Buiza-Rueda (Instituto de Biomedicina de Sevilla (IBiS), Hospital Universitario Virgen del Rocío/CSIC/Universidad de Sevilla, Seville),
Ana Cámara (Hospital Clinic de Barcelona),
Fátima Carrillo (Instituto de Biomedicina de Sevilla (IBiS), Hospital Universitario Virgen del Rocío/CSIC/Universidad de Sevilla, Seville),
Mario Carrión-Claro (Instituto de Biomedicina de Sevilla (IBiS), Hospital Universitario Virgen del Rocío/CSIC/Universidad de Sevilla, Seville),
Debora Cerdan (Hospital General de Segovia, Segovia),
Jordi Clarimón (Memory Unit, Department of Neurology, IIB Sant Pau, Hospital de la Santa Creu i Sant Pau, Universitat Autònoma de Barcelona and Centro de Investigación Biomédica en Red en Enfermedades Neurodegenerativas (CIBERNED), Madrid)
Yaroslau Compta (Hospital Clinic de Barcelona),
Beatríz de la Casa (Hospital Gregorio Marañon, Madrid)
Monica Diez-Fairen (Fundació Docència i Recerca Mútua de Terrassa and Movement Disorders Unit, Department of Neurology, University Hospital Mutua de Terrassa, Terrassa, Barcelona.),
Oriol Dols-Icardo (Memory Unit, Department of Neurology, IIB Sant Pau, Hospital de la Santa Creu i Sant Pau, Universitat Autònoma de Barcelona, Barcelona, and Centro de Investigación Biomédica en Red en Enfermedades Neurodegenerativas (CIBERNED), Madrid),
Jacinto Duarte (Hospital General de Segovia, Segovia),
Raquel Duran (Centro de Investigacion Biomedica, Universidad de Granada, Granada),

Francisco Escamilla-Sevilla (Hospital Universitario Virgen de las Nieves, Instituto de Investigación Biosanitaria de Granada, Granada),
Mario Ezquerra (Hospital Clinic de Barcelona)
Cici Feliz (Departmento de Neurologia, Instituto de Investigación Sanitaria Fundación Jiménez Díaz, Madrid, Spain),
Manel Fernández (Hospital Clinic de Barcelona),
Rubén Fernández-Santiago (Hospital Clinic de Barcelona),
Ciara Garcia (Hospital Universitario Central de Asturias, Oviedo),
Pedro García-Ruiz (Instituto de Investigación Sanitaria Fundación Jiménez Díaz, Madrid),
Pilar Gómez-Garre (Instituto de Biomedicina de Sevilla (IBiS), Hospital Universitario Virgen del Rocío/CSIC/Universidad de Sevilla, Seville),
Maria Jose Gomez Heredia (Hospital Universitario Virgen de la Victoria, Malaga),
Isabel Gonzalez-Aramburu (Hospital Universitario Marqués de Valdecilla-IDIVAL, Santander) ,
Ana Gorostidi Pagola (Instituto de Investigación Sanitaria Biodonostia, San Sebastián),
Janet Hoenicka (Institut de Recerca Sant Joan de Déu, Barcelona),
Jon Infante (Hospital Universitario Marqués de Valdecilla-IDIVAL and University of Cantabria, Santander, and Centro de Investigación Biomédica en Red en Enfermedades Neurodegenerativas (CIBERNED),
Silvia Jesús (Instituto de Biomedicina de Sevilla (IBiS), Hospital Universitario Virgen del Rocío/CSIC/Universidad de Sevilla, Seville),
Adriano Jimenez-Escrig (Hospital Universitario Ramón y Cajal, Madrid),
Jaime Kulisevsky (Movement Disorders Unit, Department of Neurology, IIB Sant Pau, Hospital de la Santa Creu i Sant Pau, Universitat Autònoma de Barcelona, Barcelona, and Centro de Investigación Biomédica en Red en Enfermedades Neurodegenerativas (CIBERNED)),
Miguel A. Labrador-Espinosa (Instituto de Biomedicina de Sevilla (IBiS), Hospital Universitario Virgen del Rocío/CSIC/Universidad de Sevilla, Seville),
Jose Luis Lopez-Sendon (Hospital Universitario Ramón y Cajal, Madrid),
Adolfo López de Munain Arregui (Instituto de Investigación Sanitaria Biodonostia, San Sebastián),
Daniel Macias (Instituto de Biomedicina de Sevilla (IBiS), Hospital Universitario Virgen del Rocío/CSIC/Universidad de Sevilla, Seville),
Irene Martínez Torres (Department of Neurology, Instituto de Investigación Sanitaria La Fe, Hospital Universitario y Politécnico La Fe, Valencia),
Juan Marín (Movement Disorders Unit, Department of Neurology, IIB Sant Pau, Hospital de la Santa Creu i Sant Pau, Universitat Autònoma de Barcelona, Barcelona, and Centro de Investigación Biomédica en Red en Enfermedades Neurodegenerativas (CIBERNED)),
Maria Jose Marti (Hospital Clinic Barcelona),
Juan Carlos Martínez-Castrillo (Instituto Ramón y Cajal de Investigación Sanitaria, Hospital Universitario Ramón y Cajal, Madrid),
Carlota Méndez-del-Barrio (Instituto de Biomedicina de Sevilla (IBiS), Hospital Universitario Virgen del Rocío/CSIC/Universidad de Sevilla, Seville),
Manuel Menéndez González (Hospital Universitario Central de Asturias, Oviedo),
Marina Mata (Department of Neurology, Hospital Universitario Infanta Sofía, Madrid, Spain)
Adolfo Mínguez (Hospital Universitario Virgen de las Nieves, Granada, Instituto de Investigación Biosanitaria de Granada),
Pablo Mir (Instituto de Biomedicina de Sevilla (IBiS), Hospital Universitario Virgen del Rocío/CSIC/Universidad de Sevilla, Seville),
Elisabet Mondragon Rezola (Instituto de Investigación Sanitaria Biodonostia, San Sebastián),
Esteban Muñoz (Hospital Clinic Barcelona),
Javier Pagonabarraga (Movement Disorders Unit, Department of Neurology, IIB Sant Pau, Hospital de la Santa Creu i Sant Pau, Universitat Autònoma de Barcelona, Barcelona, and Centro de Investigación Biomédica en Red en Enfermedades Neurodegenerativas (CIBERNED)),
Berta Pascual-Sedano (Unidad de Trastornos del Movimiento, Hospital Sant Pau, Barcelona)
Pau Pastor (Fundació Docència i Recerca Mútua de Terrassa and Movement Disorders Unit, Department of Neurology, University Hospital Mutua de Terrassa, Terrassa, Barcelona.),
Francisco Perez Errazquin (Hospital Universitario Virgen de la Victoria, Malaga),
Teresa Periñán-Tocino (Instituto de Biomedicina de Sevilla (IBiS), Hospital Universitario Virgen del Rocío/CSIC/Universidad de Sevilla, Seville),

Javier Ruiz-Martínez (Hospital Universitario Donostia, Instituto de Investigación Sanitaria Biodonostia, San Sebastián),
Clara Ruz (Centro de Investigacion Biomedica, Universidad de Granada, Granada),
Antonio Sanchez Rodriguez (Hospital Universitario Marqués de Valdecilla-IDIVAL, Santander),
María Sierra (Hospital Universitario Marqués de Valdecilla-IDIVAL, Santander),
Esther Suarez-Sanmartin (Hospital Universitario Central de Asturias, Oviedo),
Cesar Tabernero (Hospital General de Segovia, Segovia),
Juan Pablo Tartari (Fundació Docència i Recerca Mútua de Terrassa and Movement Disorders Unit, Department of Neurology, University Hospital Mutua de Terrassa, Terrassa, Barcelona),
Cristina Tejera-Parrado (Instituto de Biomedicina de Sevilla (IBiS), Hospital Universitario Virgen del Rocío/CSIC/Universidad de Sevilla, Seville),
Eduard Tolosa (Hospital Clinic Barcelona),
Francesc Valldeoriola (Hospital Clinic Barcelona),
Laura Vargas-González (Instituto de Biomedicina de Sevilla (IBiS), Hospital Universitario Virgen del Rocío/CSIC/Universidad de Sevilla, Seville),
Lydia Vela (Department of Neurology, Hospital Universitario Fundación Alcorcón, Madrid),
Francisco Vives (Centro de Investigacion Biomedica, Universidad de Granada, Granada).

**Austria:**
Alexander Zimprich (Department of Neurology, Medical University of Vienna, Austria)

**Norway:**
Lasse Pihlstrom (Department of Neurology, Oslo University Hospital, Oslo, Norway),
Mathias Toft (Department of Neurology and Institute of Clinical Medicine, Oslo University Hospital, Oslo, Norway)

**Estonia**:
Sulev Koks (Department of Pathophysiology, University of Tartu, Tartu, Estonia; Department of Reproductive Biology, Estonian University of Life Sciences, Tartu, Estonia),
Pille Taba (Department of Neurology and Neurosurgery, University of Tartu, Tartu, Estonia)

**Israel:**
Sharon Hassin-Baer (The Movement Disorders Institute, Department of Neurology and Sagol Neuroscience Center, Chaim Sheba Medical Center, Tel-Hashomer, 5262101, Ramat Gan, Israel, Sackler Faculty of Medicine, Tel Aviv University, Tel Aviv, Israel)
